# ReALLEN: structural variation discovery in cancer genome by sensitive analysis of single-end reads

**DOI:** 10.1101/506329

**Authors:** Ryo Kanno, Daisuke Tanaka, Hideaki Nanamiya, Takao Isogai

## Abstract

**Background:** The structural abnormalities in chromosomes are important issues in cancer genomics. Next generation sequencing technologies have big potentials to detect the structural variations precisely and comprehensively. Nevertheless, it is still difficult problem to detect large structural variations from short read sequence data. Major efforts have been achieved with paired-end reads, since discordant pairs directly reflect the existence of large rearrangement. Furthermore, approaches to detect structural variations from single-end reads are still worthwhile challenge because they allow wide choices of sequencing platforms.

**Result:** We present ReALLEN, a series of tools to detect genomic rearrangement with base-pair resolution from single-end reads provided by next generation sequencing. We examined the performance of ReALLEN using simulated dataset and real dataset sequenced by Ion Torrent systems. In most cases on the simulated dataset, ReALLEN showed nearly 100% precision and better sensitivity than other major tools. Notably, ReALLEN showed stable scores even if it was on some unfavorable conditions, for example, low coverage or small variant size. On the real dataset sequenced by Ion Torrent systems, ReALLEN accurately found an insertional translocation that was crucial for the diagnosis of chronic myeloid leukemia.

**Conclusion:** ReALLEN is useful to researchers in finding genomic rearrangements. It will contribute to discovery of cancer-specific fusion proteins, precise diagnosis of known types of cancers, and understanding of genetic diseases caused by abnormal chromosomes.

## Background

Normal cells could postnatally turn into cancer cells by acquisition of somatic mutations in their genome [1]. The mutations that affect large sequence alteration (typically > 50 bp) are called structural variations (SVs), and are distinguished from single nucleotide variations (SNVs) [2, 3]. The somatic rearrangement of chromosomes that produces deleterious fusion protein, for example, BCR-ABL, PML-RARα, or EML4-ALK, is known to promote carcinogenesis [4]. Although some fusion proteins have been identified by traditional approaches [3, 4, 5], systematic information for various types of cancer diagnosis is still insufficient. The comprehensive discovery of SVs in cancer genome will make it possible to find novel fusion proteins, and will contribute to the targeted cancer therapies. Next generation sequencing (NGS) technologies enable us to get vast amount of genetic information with lower cost, and more and more whole genome sequencing (WGS) data are accumulating [6, 7]. Nevertheless, there is no gold standard method for the discovery of SVs from short read data.

Four distinct approaches have been reported to detect SV signatures from NGS data: Read-pair (RP), Read-count (RC), Split-read (SR) and de novo assemble (AS) [2, 8]. BreakDancer, DELLY, LUMPY, and PRISM are able to detect SVs from paired-end (PE) reads with RP or complexed methods of RP and SR [9, 10, 11, 12]. HiSeq systems (Illumina) produce PE data [7], and properly cooperate with these tools. On the other hand, other systems that produce only single-end (SE) data, including the Ion Proton (Thermo Fisher Scientific), cannot utilize them effectively. SR is generally lower at the sensitivity than RP, because SR can find only breakpoint (BP) that crosses just over a read, whereas RP can find BP in large gap between paired reads. However, it is an advantage that SR yields base-pair resolution. Besides, SR becomes a more powerful approach as the readable length by NGS gets longer. Socrates, CREST, Gustaf, and SPLITREAD detect SVs from SE data with SR-based methods [13, 14, 15, 16, 17].

We present new software, ReALLEN (<u>Re</u>arrangement <u>A</u>nalysis by <u>L</u>ong-<u>L</u>eap <u>E</u>stimation on <u>N</u>GS data), for the discovery of SVs with base-pair resolution from SE data. Notably, it has three features: (i) ReALLEN is specifically designed for cancer genome data. The content of reads with somatic mutations is usually very little in cancer sample, because most cancer tissues heterogeneously include both normal DNA and cancer DNA. Thus, high sensitivity is necessary to detect the weak SV signatures in cancer samples. ReALLEN improves the sensitivity by integrating two different SR-based algorisms, soft-clip split [18] and fixed-length split. (ii) ReALLEN is able to find both small SVs (deletions, inversions, and tandem duplications range from 25 bp to 400 bp) and large SVs (inter-chromosomal and intra-chromosomal translocations; deletions, inversions, and tandem duplications more than 400 bp). The existing RP-based methods have more or less problems to find the small SVs, although such SVs are important in coding regions. (iii) ReALLEN is optimized for SE data. Because ReALLEN has been developed for Ion Torrent platforms at the beginning, all parameters has been designed to provide sufficient result without PE data. In spite of the primal purpose, ReALLEN will work well on various sequencing platforms, considering that it showed sufficient scores on simulated dataset as well as on real dataset sequenced by Ion Torrent systems.

## Implementation

The main algorithm of ReALLEN was implemented in Ruby. Startup scripts for easy-run were implemented in Bash script. ReALLEN requires aligned sequence data in SAM or BAM format as input. ReALLEN outputs the position of SVs in CSV format with annotations or in simple BLAST-like format [19]. ReALLEN is designed to work in cooperation with any mapping software although we recommend BWA-mem [20, 21] and TMAP (Thermo Fisher Scientific). References and annotations for hg19 (UCSC human genome 19) are included in build-in package. We show the workflow of ReALLEN in Figure 1. The steps are as follows:

1. Mapping filter. At the first step, ReALLEN removes well-mapped reads to achieve fast calculation because these reads rarely have BPs. ReALLEN counts mismatch alignments (insertion, deletion, and soft-clip) of each read, and discards reads in which the count does not exceed threshold. Applying larger threshold values at this step increases the entire speed of the calculation while it slightly decreases the sensitivity.
2. Soft-clip split and realignment. If a read has long soft-clipped sequence, the read is separated into the soft-clipped end and the opposite end. Both ends of the read are realigned by BWA-mem.
3. Fixed-length split and realignment. If a read does not have enough length of soft-clipped sequence, the read is separated into two ends with fixed length. The reads with incorrect alignment are rescued at this step. Both ends of the read are also realigned by BWA-mem.
4. Mapping uniqueness filter. If a read is not uniquely mapped to the reference by the realignment, the read is removed. The uniqueness is evaluated by the difference of the scores between primary alignment and secondary alignment. This step is significantly effective to reduce false positive, especially when we target the sequence which has repetitive regions. Low quality mappings are also removed at this step.
5. Pairing test The filtered reads are paired with the opposite ends of the original reads. The pairing test is performed to find discordant pairs. If two reads of a pair are located on different chromosomes or inconsistent orientations, the pair is discordant. Besides, if the gap (L_gap_) is larger than the given size (the minimum SV size), the pair is discordant. L_gap_ is defines as

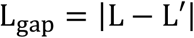

where L is the length of the original read and L’ is the distance between the realigned terminals of the read pair. The discordant pairs are the candidates of the BPs.
6. Search accurate positions of breakpoints. Since the accurate positions of BPs cannot be identified by fixed-length split algorithm, ReALLEN calculates them by split mapping algorithm [11, 16]. The original reads are mapped to the reference around the putative BPs in a way that allows a jump from one region to another region.
7. Clustering and counting filter. The discordant pairs are classified by modified k-means clustering. The radius (r) of clusters is given instead of number of clusters. A discordant pair, defined as an element, has six values: the chromosome (c_1_), the position (p_1_), and the orientation (d_1_) of one end (left-end) and the chromosome (c_2_), the position (p_2_), and the orientation (d_2_) of the opposite end (right-end). The order of chromosomes is uniquely defined (e.g. chromosome 1 < chromosome 2). The left-end and the right-end are swapped if C_2_ is smaller than c_1_ or p_2_ is smaller than p_1_. Each element is assigned to the cluster that firstly satisfies the following conditions:

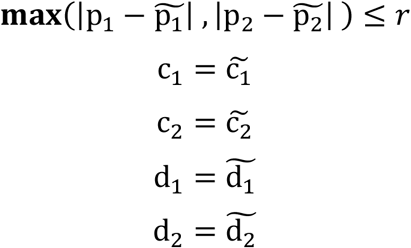

where 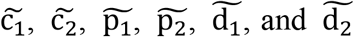 are median values for c_1_, c_2_, p_1_, p_2_, d_1_, and d_2_ of all elements in that cluster, respectively. If no cluster satisfies the conditions, the element is assigned to a new cluster. These processes are repeated until all elements are classified. After the classification, the cluster that have less than N elements are removed (N is a threshold value). When we suppose that error read is aligned to a random position, the possibility to align N error reads to an identical position is (1/s)^N-1^, where s is size of reference. Therefore, taking more than 1 for N, the false positive rate would be suppressed even if the size of reference increased.
8. Local detection. This optional step is effective to find specific fusion proteins. Ideally, a rearrangement which produces abnormal protein is supposed to occur within exons or around exons. Besides, the limited number of genes can be targeted for cancer analysis [22]. ReALLEN keeps only BPs that connect the selected regions when turning on this option, and enables us to focus on the important genes. We will decrease the false positive rate significantly if we have well-chosen list of target genes.
9. Annotation and secondary filtering. We prepare gene names, exon regions, and cytobands as annotations for hg19. All annotations can be attached to the output and viewed in CSV table. Simple estimation of SV type and secondary filtering can be applied at this step.

**Figure 1.**
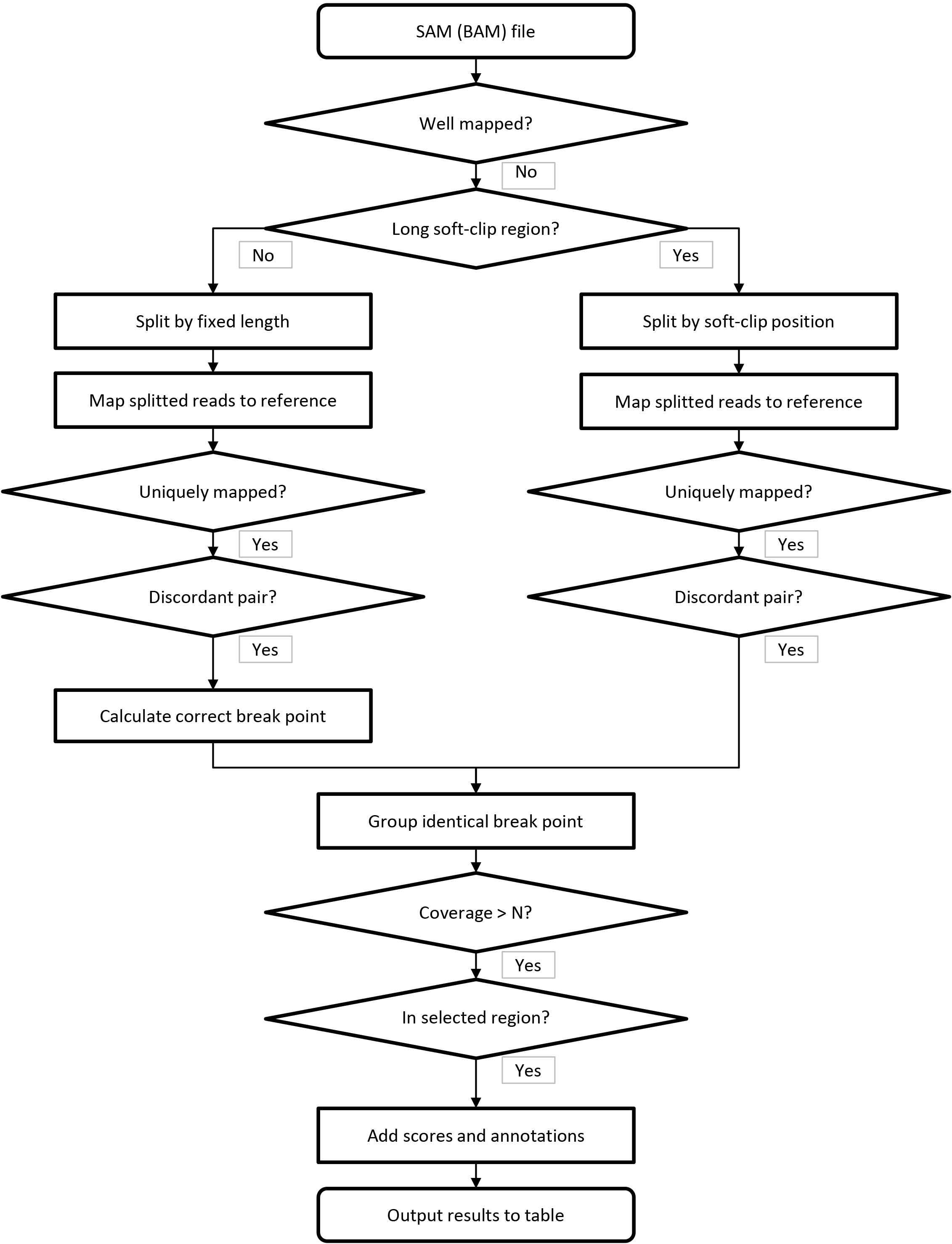
ReALLEN Workflow. SE reads are separated to produce artificial pairs by soft-clip split or fixed-length split. Every pairs are remapped, filtered, and classified to find discordant pairs. These steps are displayed with common flowchart symbols.

## Results and Discussion

### Preparation of simulated dataset

In order to reduce CPU time for the calculation, we used short partial sequences of hg19 for simulated dataset. The regions were carefully chosen not to bring sequencing bias after a preliminary test (data not shown). The artificial sequences (test sequence #1) with inter-chromosomal and intra-chromosomal translocations were generated as follows: we randomly chose 20 positions on the partial sequences of hg19 (60,000,000-61,000,000 on chromosome 16 and 22,000,000-23,000,000 on chromosome 21), divided them into pieces at the positions, and randomly connected the pieces to reconstruct two rearranged sequences. 8 patterns of the sequences were individually created in this way.

The artificial sequences (test sequence #2) with specific SV types were generated as follows: we inserted SVs of each type and each size category at 10 positions that we randomly chose on the partial sequence of hg19 (60,000,000-61,000,000 on chromosome 16). The SV types were deletion, inversion, and tandem duplication. The size categories were 25-49 bp, 50-99 bp, 100-199 bp, 200-399 bp, 400-799 bp, and 800-1599 bp. 8 patterns of the sequences were individually created in this way.

Dataset #1 and dataset #2 were generated from test sequence #1 and test sequence #2, respectively. The artificial sequences were mixed with the corresponding normal sequences at a ratio of 50%, which imitated autosomal heterozygous germline event. By the simulation using wgsim [23], the read data were generated at each condition (coverage, read type, and read length). The coverages were 5×, 10×, 20×, and 50×. The read types and the read lengths were 100 bp PE, 200 bp SE, 300 bp SE, and 400 bp SE. The insert size of PE was 300 bp (sigma=50). The generated reads were mapped to hg19 by BWA-mem with default parameters.

Dataset #3 was generated from test sequence #1. The parameters of wgsim were the same as dataset #1. The reads in dataset #3 were aligned by BWA-mem with different parameters. Match scores (‘-A’ option on BWA-mem) were 1 and 3 for Alignment #1 and Alignment #2, respectively. The larger match score for Alignment #2 allowed incorrect mapping with less soft-clipping.

### Comparison with other SR-based tools

We examined the performance of ReALLEN and other SR-based tools: CREST [14], Socrates [13], and Lumpy (only SR mode) [10]. SVs called by these tools on the simulated dataset (dataset #1, #2, and #3) were classified into true positive (TP), which agreed with the correct BP, and remaining false positive (FP). A tolerance of 500 bp was allowed even if the called position by each tool was shifted from the correct BP. Redundant calls for a single event were counted as one TP. The BPs not identified were counted as false negative (FN). Precision, Recall, and F-value were defined as follows:

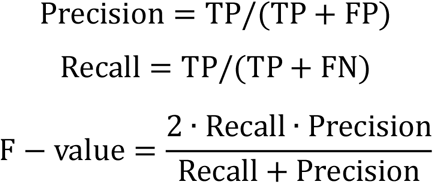

Mean shift was average distance between the correct BP and the called position. Precisions, Recalls, F-values, and Mean shifts on the simulated dataset were calculated (Table S1, Table S2, and Table S3).

The performance of ReALLEN on the low coverage conditions was compared with those of the other SR-based tools. Figure 2A shows the performance to find large translocations on dataset #1. The entire graphs indicate a trend that the higher coverage and the longer read length bring better performance. On the coverage of 50×, all tools showed F-values close to 1.0 with 200 bp, 300 bp, and 400 bp reads (Fig. 2A). This implies that there are no significant difference among these tools on the high coverage conditions. On the other hand, ReALLEN showed the highest F-values on the coverages of 5× and 10× with any of the read lengths. The difference was obvious on dataset #2. Figure 2B shows the performance to find SVs of the specific types and the size categories on dataset #2 on the coverage of 10× with 300 bp reads. ReALLEN showed the highest F-values for the small deletions and the small tandem duplications, without significant reduction compared to large SVs. The F-values of CREST and Lumpy (SR) were clearly decreased for the small deletions (< 100 bp) and the small tandem duplications (< 200 bp). The effect was weaker for Socrates than CREST or Lumpy (SR) although it was not a negligible quantity. For the small inversions, all of these tools showed lower F-values. Overall, these SR-based tools called highly precise positions of BPs with mean shifts less than 2 bp (Table S1 and Table S2). Precisions by these tools were nearly 1.0 in almost every cases (Table S1 and Table S2). As a result of comparison with each tool, the performance of ReALLEN on the low coverage conditions was better than those of the other SR-based tools.

**Figure 2.**
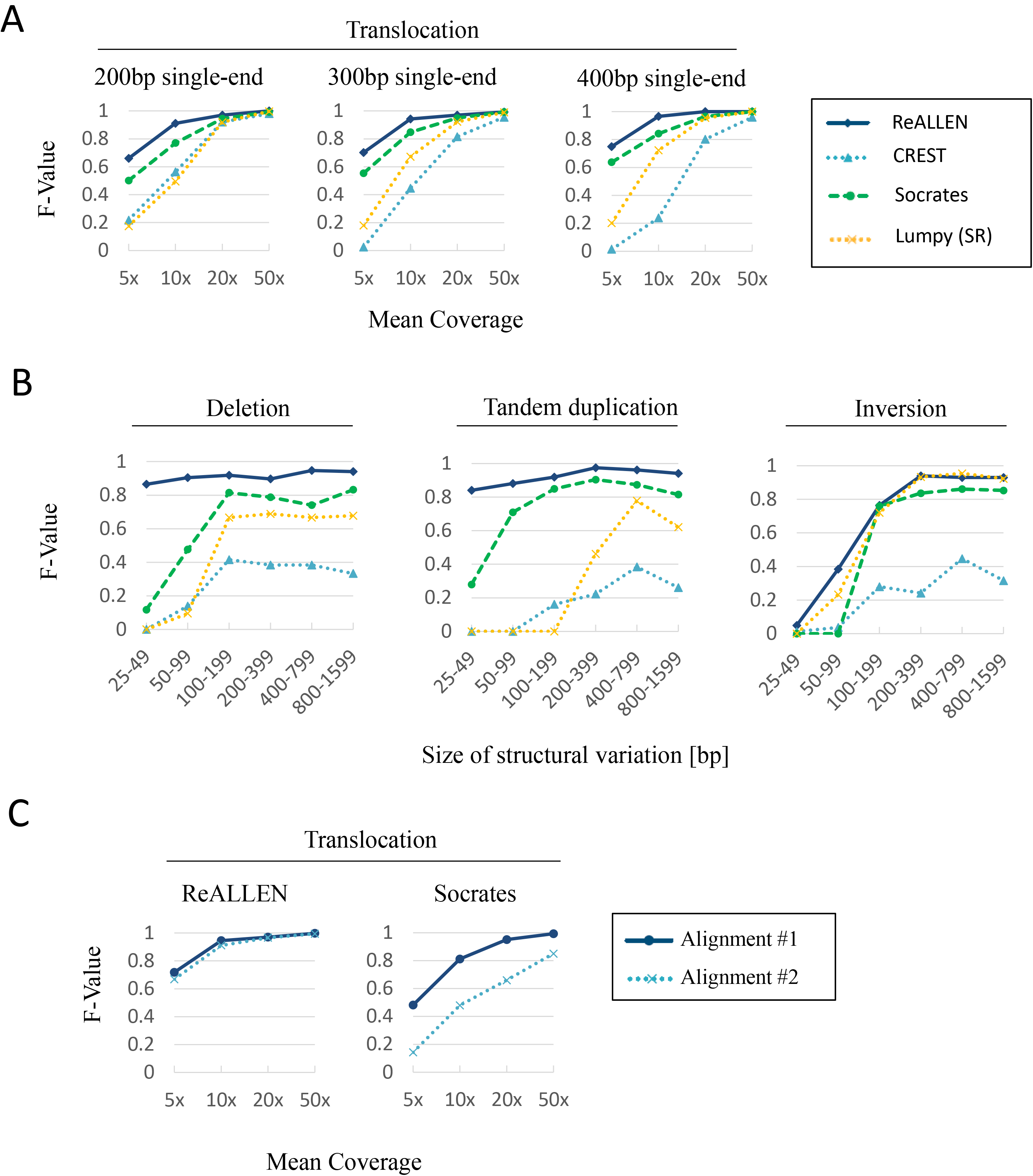
Evaluation of ReALLEN and the other SR-based tools. (A) F-values by Socrates, CREST, Lumpy (only SR mode), and ReALLEN for 200 bp SE, 300 bp SE, and 400 bp SE on the simulated translocations (dataset #1). The coverages are 5×, 10×, 20×, and 50×. (B) F-values by Socrates, CREST, Lumpy (only SR mode), and ReALLEN for 300 bp SE on the specific types of simulated SVs (dataset #2). The inserted SVs are deletions, inversions, and tandem duplications ranged from 25 bp to 1600 bp. The coverage is 10×. (C) F-values by Socrates and ReALLEN for 300 bp SE on the simulated translocations (dataset #3). Alignment #2 (match score=3) brings longer and less precise match than Alignment #1 (match score=1). The coverages are 5×, 10×, 20×, and 50×.

The effect of misalignment was examined in Figure 2C. Socrates showed the second best score to find translocations, deletions, and tandem duplications on dataset #1 and dataset #2. One of the difference between ReALLEN and Socrates is the usage of fixed-length split. This approach will affect the dependency on the first alignment. It is indicated that the performance of simple soft-clip split algorithm is affected by the result of aligner [13]. Figure 2C shows the performance on two different mappings with the same read data (dataset #3). By increasing the match score, the mapping was more tolerant of mismatches and indels. This imitated, or emphasized, the situation where aligner needed to allow discordant bases in alignment due to sequencing errors. When we increased the match score for the mapping, the F-values of Socrates clearly decreased, while the F-values of ReALLEN were nearly unchanged (Alignment #2 in Fig. 2C). Additionally, we tested dataset with high error rate (Fig. S1A and Table S4). The reduction of the performance of ReALLEN was smaller than that of Socrates on the dataset with 5% error rate. We examined the effect of the fixed-length split step, by comparing the results with or without this step (Fig. S1B and Table S3). We found that the performance for dataset#3 without the fixed-length split step was slightly lower than that with this step. Although it is difficult to evaluate which process actually contributed to the improvement of the performance, the fixed-length split step certainly affected the rescue of the inaccurate mappings. These results suggest that ReALLEN, which uses the combination of two algorithms, is rather independent on misalignment and resistant to sequencing errors.

### Comparison with RP-based tools

- The average read length is important for the performance, and it is dependent on the sequencing platforms. Generally, single-end platforms are able to sequence longer reads than pair-end platforms [7]. Therefore, we compared the scores of long SE reads (300 bp) by ReALLEN with the scores of short PE reads (100 bp) by Delly [9] and Lumpy (SR+RP mode) [10] on the simulated dataset (Table S1 and Table S2). When we compared the F-values for large translocations on dataset #1, there was no significant difference on the coverages of 20× and 50×, and ReALLEN showed a little better score on the coverage of 5× than the other tools (Fig. 3A). The result suggests that ReALLEN is comparable in performance to the RP-based tools. Figure 3B is focused on the specific types and the size categories on the coverage of 10×. All tools correctly called SVs more than 400 bp. However, Delly and Lumpy were much worse at calling the deletions ranged from 25 bp to 200 bp than ReALLEN. Delly and Lumpy rarely called the tandem duplications ranged from 25 bp to 400 bp.

**Figure 3.**
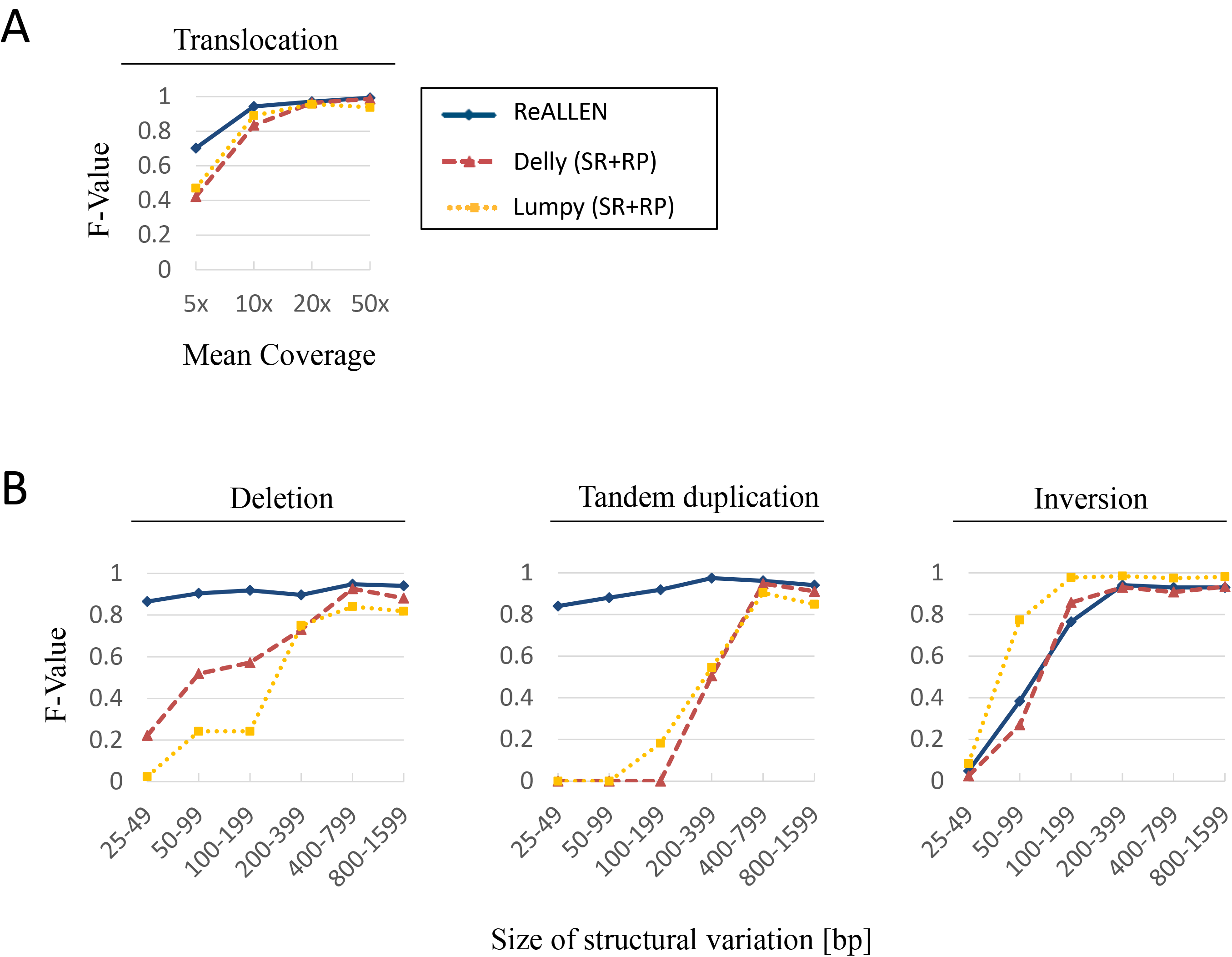
Evaluation of ReALLEN and the RP-based tools. (A) F-values by Delly and Lumpy (SR+PE mode) for 100 bp PE and ReALLEN for 300 bp SE on the simulated translocations (dataset #1). The coverages are 5×, 10×, 20×, and 50×. The insert size of PE is 300 bp (sigma=50). (B) F-values by Delly and Lumpy (SR+PE mode) for 100 bp PE and by ReALLEN for 300 bp SE on the specific types of simulated SVs (dataset #2). The inserted SVs are deletions, inversions, and tandem duplications ranged from 25 bp to 1600 bp. The coverage is 10×. The insert size of PE is 300 bp (sigma=50).

According to the low performance, RP-based methods with short PE data is inadequate for the discovery of small SVs. SR-based methods with long SE data have advantage in this respect. As a result of comparison with each tool, ReALLEN could sensitively find small SVs in contrast to the RP-based tools.

### Preparation of real dataset

The real dataset was obtained as follows: The fragment libraries of the CML sample (Supplier: ILSbio; Barcode: ILS27378; Sample ID: BM; Format: Frozen; Sample Type: Tumor; Organ Type: Bone Marrow; Species: Homo sapiens; Sex: F; Age: 50; Clinical Diagnosis: Leukemia; Pathological Diagnosis: Suggestive of Chronic Myeloid Leukemia; Description of Diagnosis: CML; Cancer Cell %: n/a; Consolidated STAGE: n/a; Grade: n/a) were constructed with Ion Xpress Plus Fragment Library kit (Thermo Fisher Scientific). Emulsion PCR and chip loading were performed with the Ion Chef system (Thermo Fisher Scientific). The five runs of sequencing (two runs with the 540 chip and three runs with the 530 chip) were performed by the Ion S5XL (Thermo Fisher Scientific). The Ion S5XL with the 530 chip produced long (about 300 bp) SE data, and that with the 540 chip produced relatively short (about 200 bp) SE data (Table S5). The sequence data were processed by the standard Ion Torrent software including TMAP. Because Socrates and Lumpy did not work on the direct output of TMAP, the reads for these tools were reprocessed by BWA-mem instead of TMAP. All read data of the five runs were merged for the test (Table S5).

### Performance test with real dataset

We examined the performance of ReALLEN using the real dataset of the CML sample sequenced by Ion Torrent systems. The SVs on the dataset were detected by ReALLEN, Socrates, CREST, and Lumpy (only SR mode). According to the output of these tools, we found inter-chromosomal translocations, t(9;22)(q34.1;q11.2). All of the tools called the translocation from 23,632,280 in chromosome 22 to 133,708,253 in chromosome 9 (Fig. 4A). The translocation corresponded to well-known rearrangement which produced BCR-ABL1 fusion protein found in CML patients [2, 24, 25]. In addition, some of the tools called the translocation from 136,052,205 in chromosome 9 to 23,632,280 in chromosome 22, and the translocation from 133,708,255 in chromosome 9 to 136,052,210 in chromosome 9. These three translocations indicated that a fragment of 2.3 million base pair on chromosome 9 was inserted into chromosome 22 (Fig. 4B). Only ReALLEN called all of the three BPs, while other tools called the deficient parts of them (Fig. 4C and Fig. S2). Even if the dataset did not have plenty of read data (the coverage was 14.8×; Fig. S3), ReALLEN showed excellent sensitivity to find the three BPs necessary for identifying the large insertional translocation in the CML sample.

**Figure 4.**
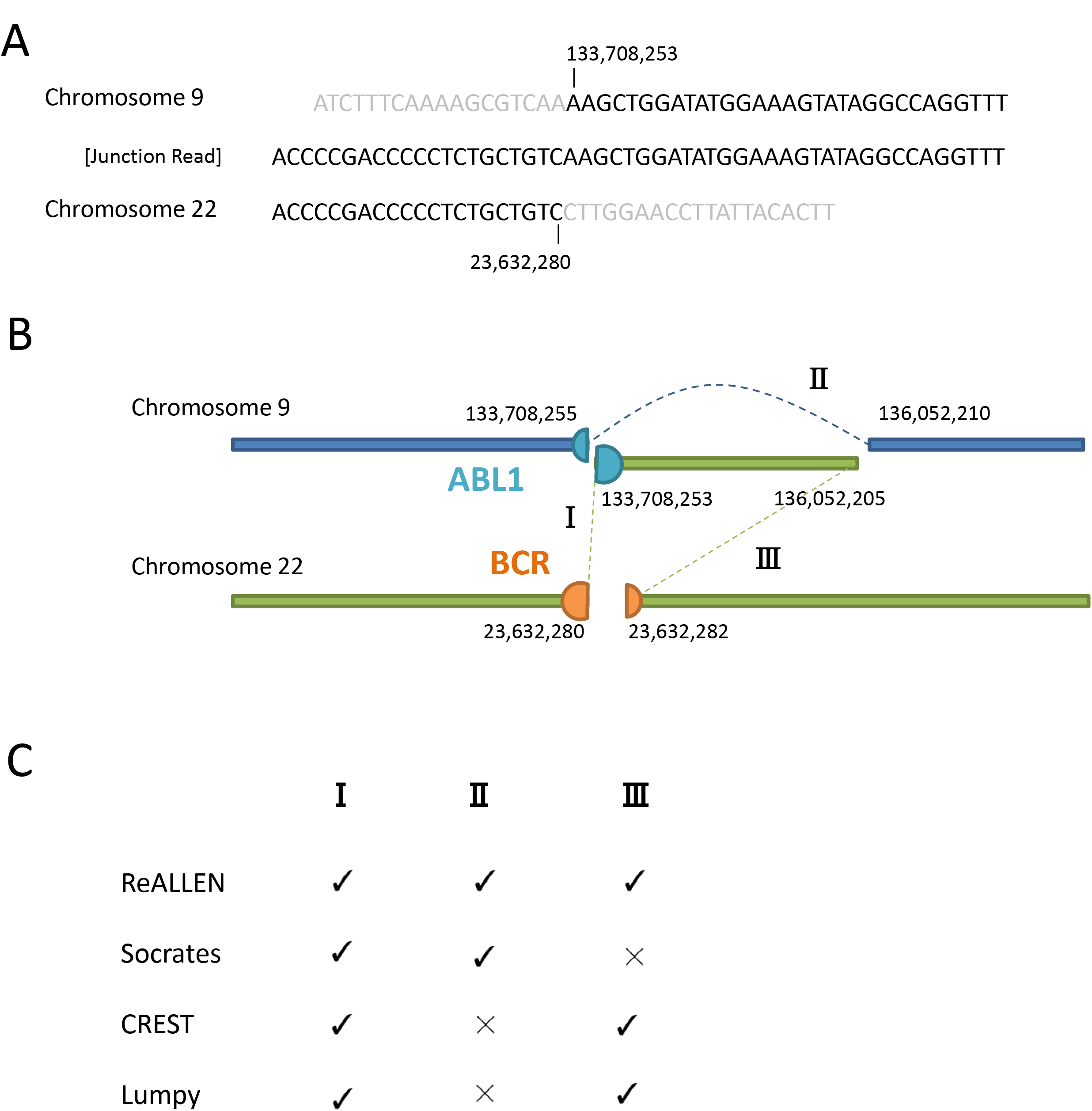
The translocation found in the CML sample. (A) The junction sequence around the translocation found in the CML sample. Either ends of the sequence across the BP are matched to the region of ABL1 (ENST00000372348) and the region of BCR (ENST00000305877). (B) Schematic view of the 2.3 M bp insertion identified with a combination of the three translocations in the CML sample. The three translocations are labeled as I, II, or III. (C) The discovery of the three SVs by ReALLEN and the other tools. Each translocation represented as I, II, or III is corresponding to the label in Figure 4B.

We investigated the detail of individual SVs called by these tools. ReALLEN called total 18,168 SVs in the sample (Table S6). After secondary filtering, it kept 6,019 SVs. Socrates called 2,978 SVs (paired results). The total number of calls by ReALLEN, Socrates, and their overlap are summarized in Figure 5A and Figure 5B. Most SVs uniquely detected by ReALLEN were the deletions less than 400 bp and the tandem duplications less than 100 bp (Fig. 5B). The result implies two things: Firstly, these small deletions and tandem duplications are major population of SVs in the actual cancer sample. Secondary, ReALLEN has potential to find SVs of these sizes with superiority to other existing tools. When we randomly sampled SVs called by ReALLEN and Socrates, their calls were judged substantially correct by manual validation (Table S7 and Fig. S4). Therefore, the large number of calls by ReALLEN reflects not low precision but high sensitivity.

**Figure 5.**
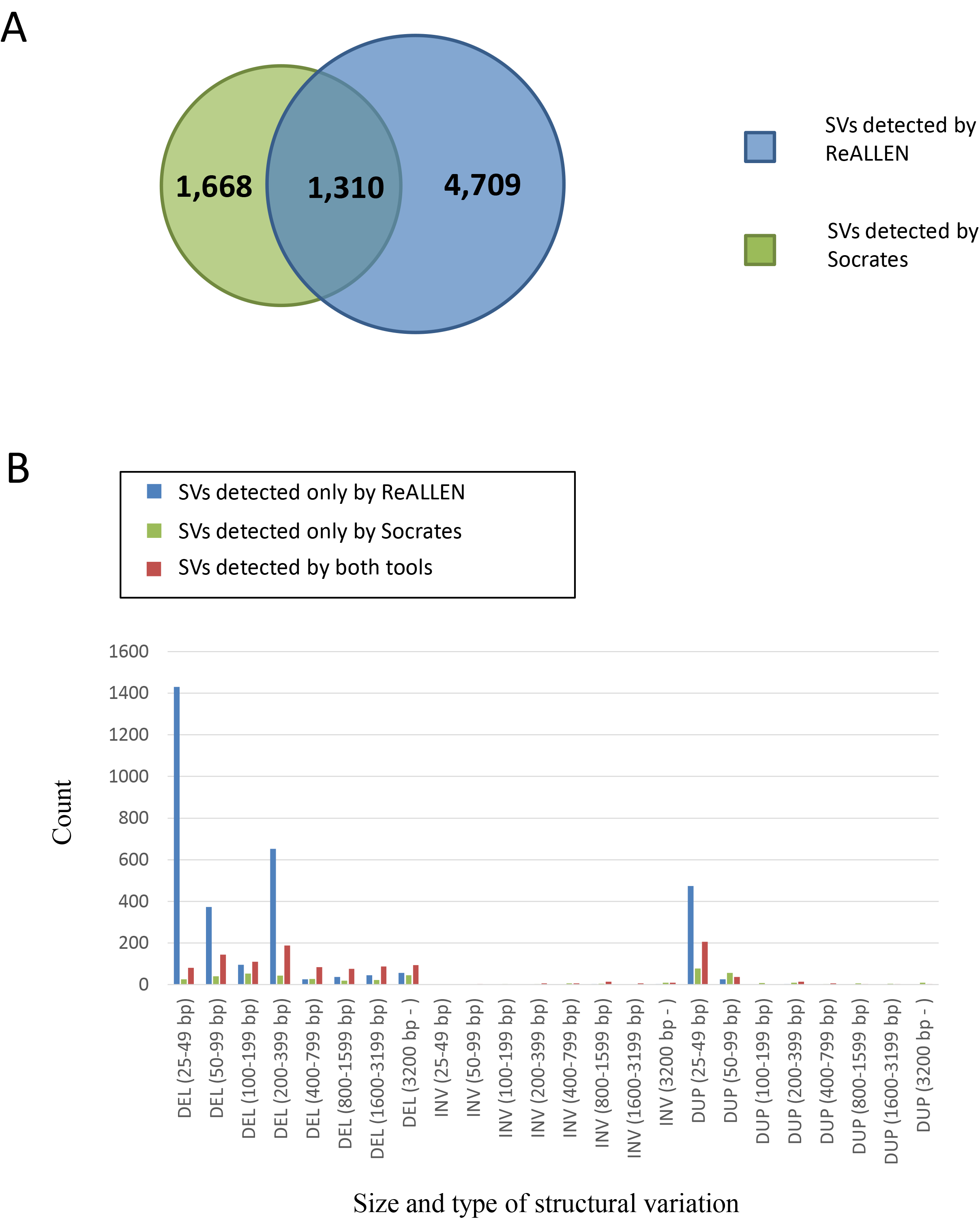
Number of SVs found in the CML sample by ReALLEN and Socrates. (A) Venn diagram that indicates the total number of SVs detected by ReALLEN, Socrates, and their overlap in the CML sample. Two calls are considered to overlap if they have the same orientation and the locations within 20 bp. (B) Number of SVs for each type and each size category detected by ReALLEN, Socrates, and their overlap in the CML sample. The types and the size categories are the same as dataset #2.

### Performance test with public dataset

In addition to our dataset, we examined the performance of ReALLEN using public dataset. We downloaded the benchmark SV calls [26, 27] for one individual in 1000 genome project (NA12878) [28, 29] from National Institute of Standards and Technology (NIST). The read data sequenced by the Ion Proton (ID: SRR1238539; Number of Bases: 32.9 G; Coverage: 11×; Mean Read Length: 176.8 bp; Read Type: single-end) were downloaded from Sequence Read Archive (SRA) [30] of National Center of Biotechnology Information (NCBI). We detected SVs by ReALLEN and Socrates, and extracted the deletions more than 50 bp for the comparison since the NIST benchmark SV calls did not contain deletions less than 50 bp (Fig. S5A). We counted the overlap of these calls (Fig. S5B). As the SVs included in the NIST benchmark SV calls were counted as TP, ReALLEN and Socrates showed almost equivalent precision (83.2% and 83.1%, respectively) whereas ReALLEN called 1.3 times more SVs (762 calls) than Socrates (579 calls). The result also supports the advantage of ReALLEN.

## Conclusions

In this study, we presented ReALLEN and evaluated its performance. ReALLEN showed the best score among the SR-based tools especially for detecting small SVs on low coverage data. Both RP-based methods and SR-based methods have their limitations and advantage. With long (300 bp) SE data, ReALLEN could find large translocations as effectively as the RP-based tools with short (100 bp) PE data. Moreover, the performance of ReALLEN to find small deletion and small tandem duplications was significantly superior to the RP-based tools. Consequently, ReALLEN worked very stably in most cases of our test.

ReALLEN found the CML-specific fusion gene, BCR-ABL1, on the real dataset sequenced by Ion Torrent systems. The result demonstrates the potential of ReALLEN to find such important translocations in cancer samples. Although Ion Torrent platforms are relatively low-cost, the output has a few peculiarities so that the original aligner, TMAP, is just suitable to their process. Since ReALLEN hardly depends on the first alignment, it is able to work in preferable cooperation with Ion Torrent systems. The high productivity of these platforms in combination with ReALLEN will contribute to discovery of cancer-specific fusion proteins, precise diagnosis of known types of cancers, and understanding of genetic diseases caused by abnormal chromosomes.

## Availability and Requirement

Project name: ReALLEN

Project home page: https://github.com/cannaryo/reallen/

Operating system: 64-bit Linux

Programming language: Ruby 2.2.3, bash shell script Other requirement: Ruby 2.2.3 or higher, Samtools, BWA

License: MIT

Any restrictions to use by non-academics: No

**List of Abbreviations**
NGS: Next Generation Sequencing
SV: Structural variation
SNV: single nucleotide variations
RP: Read-pair
RC: Read-count
SR: Split-read
AS: de novo assemble
WGS: Whole Genome Sequencing
TP: True positive
FP: False positive
FN: False negative
BP: breakpoint
CML: chronic myeloid leukemia.

## Declarations

### Ethics approval and consent to participate

The human DNA sample used in this study was collected within U.S. and International ethical guidelines. The sample was collected under Institutional Review Board (IRB) approved protocols. The detailed information was described in ILSBIO’s LETTER OF CONFIRMATION.

### Consent for publication

Not applicable.

### Availability of data and materials

The datasets used and/or analyzed during the current study available from the corresponding author on reasonable request.

### Competing interests

No competing interests.

### Funding

This work was supported by the Ministry of Economy, Trade and Industry of Japan.

### Author’s contributions

RK designed, implemented, and tested the software and wrote the manuscript. DT and HN acquired the sequence data that helped the improvement of the software. TI supervised the project and gave suggestions about the study. All authors read and approved the final manuscript.

## Acknowledgements

We thank M. Takahashi, T. Goto, M. Kawabata, and Y. Hashimoto for contribution to technical assistance. We also thank S. Watanabe, J. Imai, E. Ito, and M. Morisawa for efficient discussion.

