## Supplementary material for "ReALLEN: structural variation discovery in cancer genome by sensitive analysis of single-end reads": Figure S1 and Figure S2

A

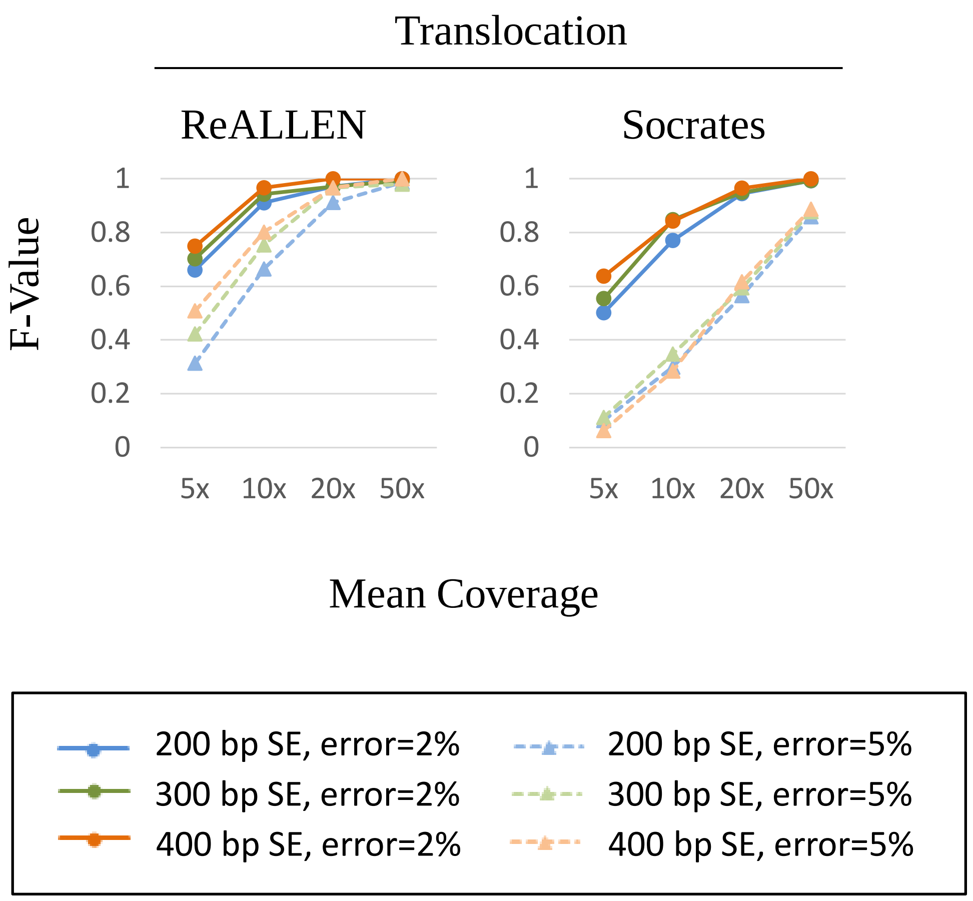


B

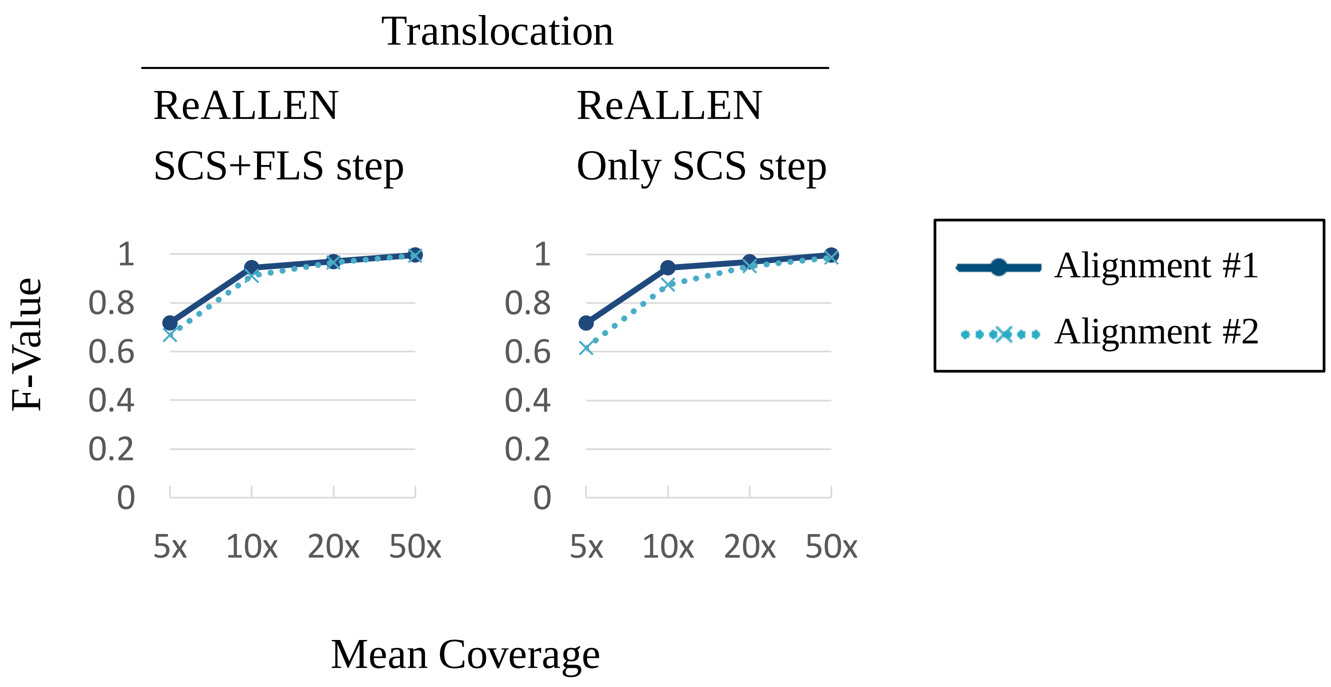


**Figure S1. Evaluation of ReALLEN and Socrates for different mappings.**
(A) Dataset #4 was generated by the same process as dataset #1, except error rate. When generating simulated reads, wgsim add 5% error on dataset #4, while dataset #1, #2, and #3 were processed with default error parameters (error rate=2%). F-values by Socrates and ReALLEN for 200 bp SE, 300 bp SE, and 400 bp SE on dataset #4 are plotted. (B) F-values calculated by ReALLEN with (left graph) or without (right graph) the fixed-length split step on dataset #3 are plotted. Alignment #1 and Alignment #2 were mapped by BWA with different match scores (1 and 3, respectively). The read length is 300 bp (SE). SCS: soft-clip split. FLS: fixed-length split.

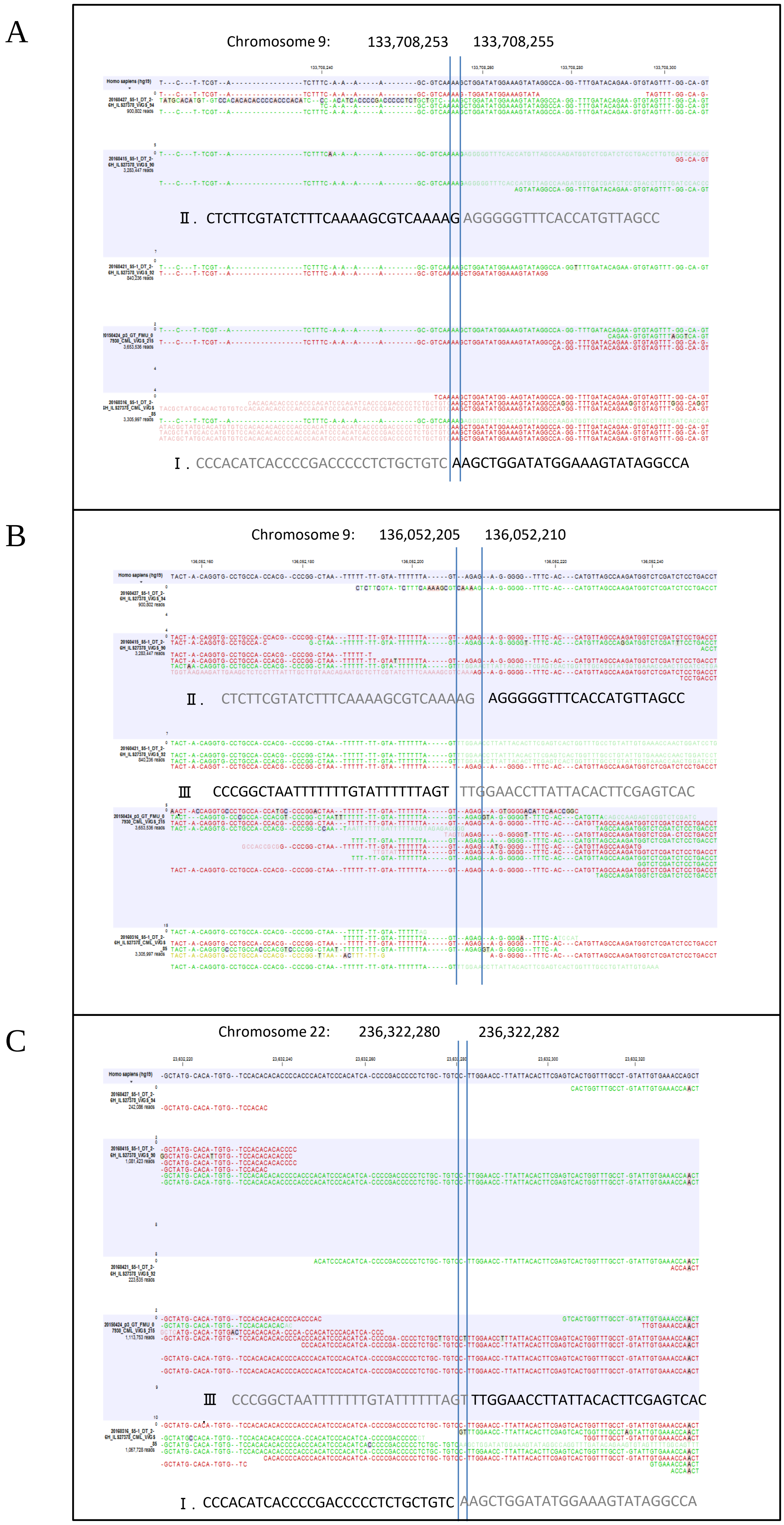


**Figure S2. The reads mappings of the CML sample.**

The reads mappings of the CML sample are visualized around the three BPs found in this study the by CLC Genomics Workbench (CLC bio). (A) The mapping around 133,708,253 and 133,708,255 in chromosome 9. (B) The mapping around 136,052,205 and 136,052,210 in chromosome 9. (C) The mapping around 236,322,380 and 236,322,282 in chromosome 22. Each sequence represented as I, II, or III is corresponding to the translocation labeled in Figure 4B. The matched sequences and the unmatched sequences are described with black and grey characters, respectively. These junction-spanning reads support the existence of the insertional translocation found by ReALLEN.
