## Supplementary material for "ReALLEN: structural variation discovery in cancer genome by sensitive analysis of single-end reads": Figure S3

**Figure S3. Confirmation of ReALLEN-called BPs in the mapped-reads data obtained by WGS.**
From the results of whole genome sequences as shown in Table S5, a set of BAM files were created by Torrent Suite Software ver.5.0.4 (Thermo Fisher Scientific). Next, these data were merged into a single file via CLC Genomics Workbench ver.7.5.1 (QIAGEN), and then the mapped-reads were visualized by the CLC Genomics Workbench. The mean coverage of the merged data was x14.8 (Table S5). The reads containing canonical BPs that were corresponding to the translocation I, II and III in Figure 4B, respectively, were shown by red arrows. The numbers of these reads were counted and were indicated in each panel. The reference sequence (hg19) of each region was shown at the bottom of each panel.
A, B : Translocation I (upper part) and translocation II (lower part) on chromosome 9.
C, D : Translocation III (upper part) and translocation II (lower part) on chromosome 9.
E : Translocation I on chromosome 22.
F : Read length histograms obtained by WGS.
The data from Run date 20160303 (upper panel) and 20160414 (lower panel) (Table S5) were shown. The details of each experimental procedure were also shown in left panel. Summaries of mean, median and mode read lengths in each WGS were indicated in upper part of the histogram.

A

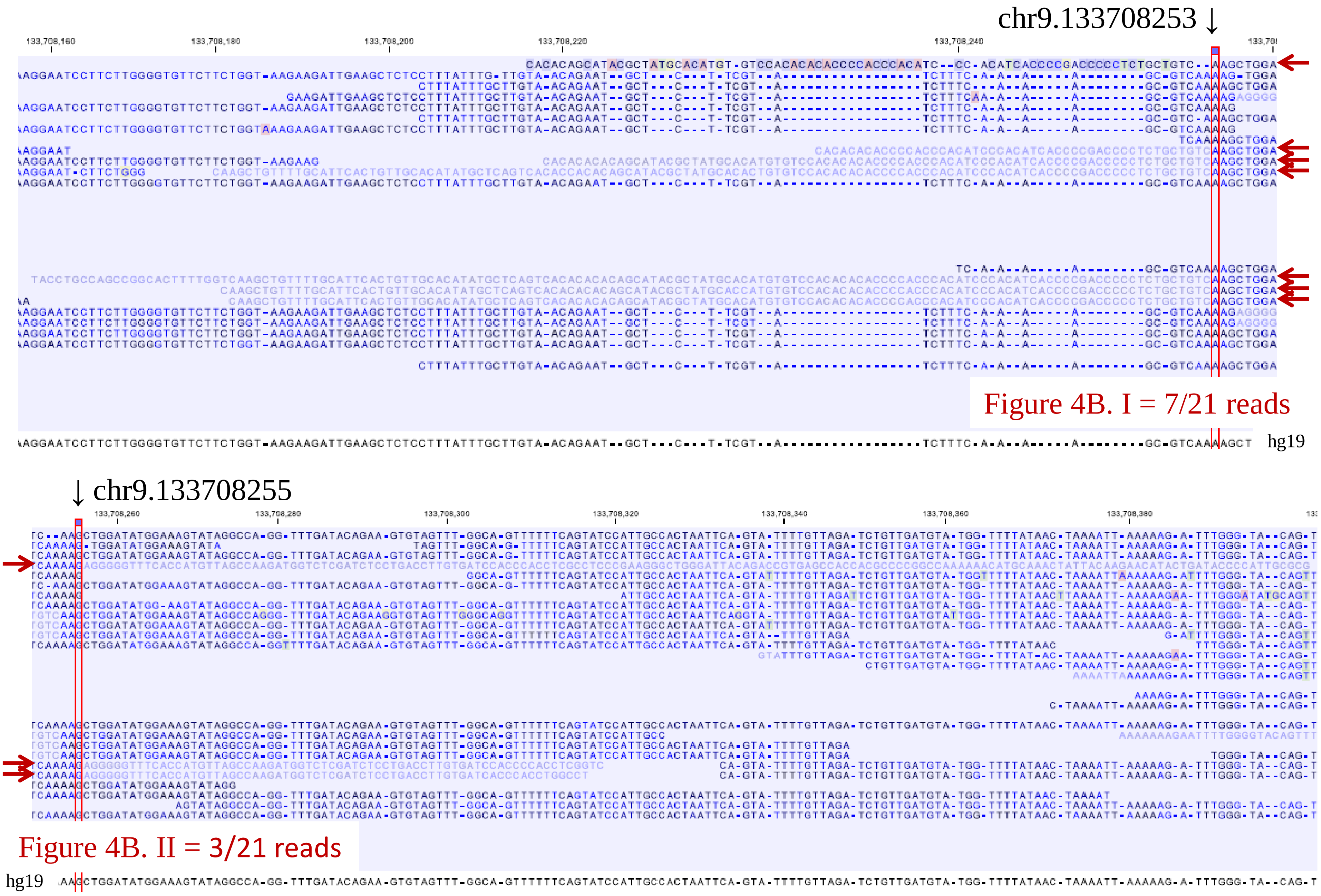
chr9.133708253
Figure 4B. I = 7/21 reads
hg19
chr9.133708255
Figure 4B. II = 3/21 reads
hg19

Figure S3 A.

B

Region of chr9.133708253～133708255

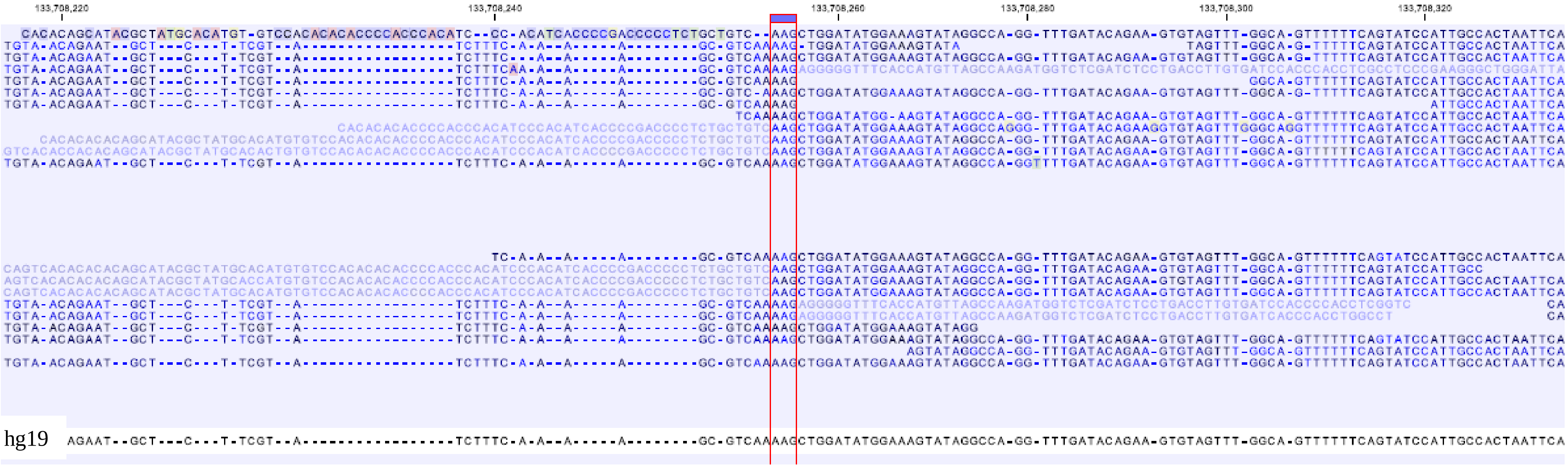


Figure S3 B.

C

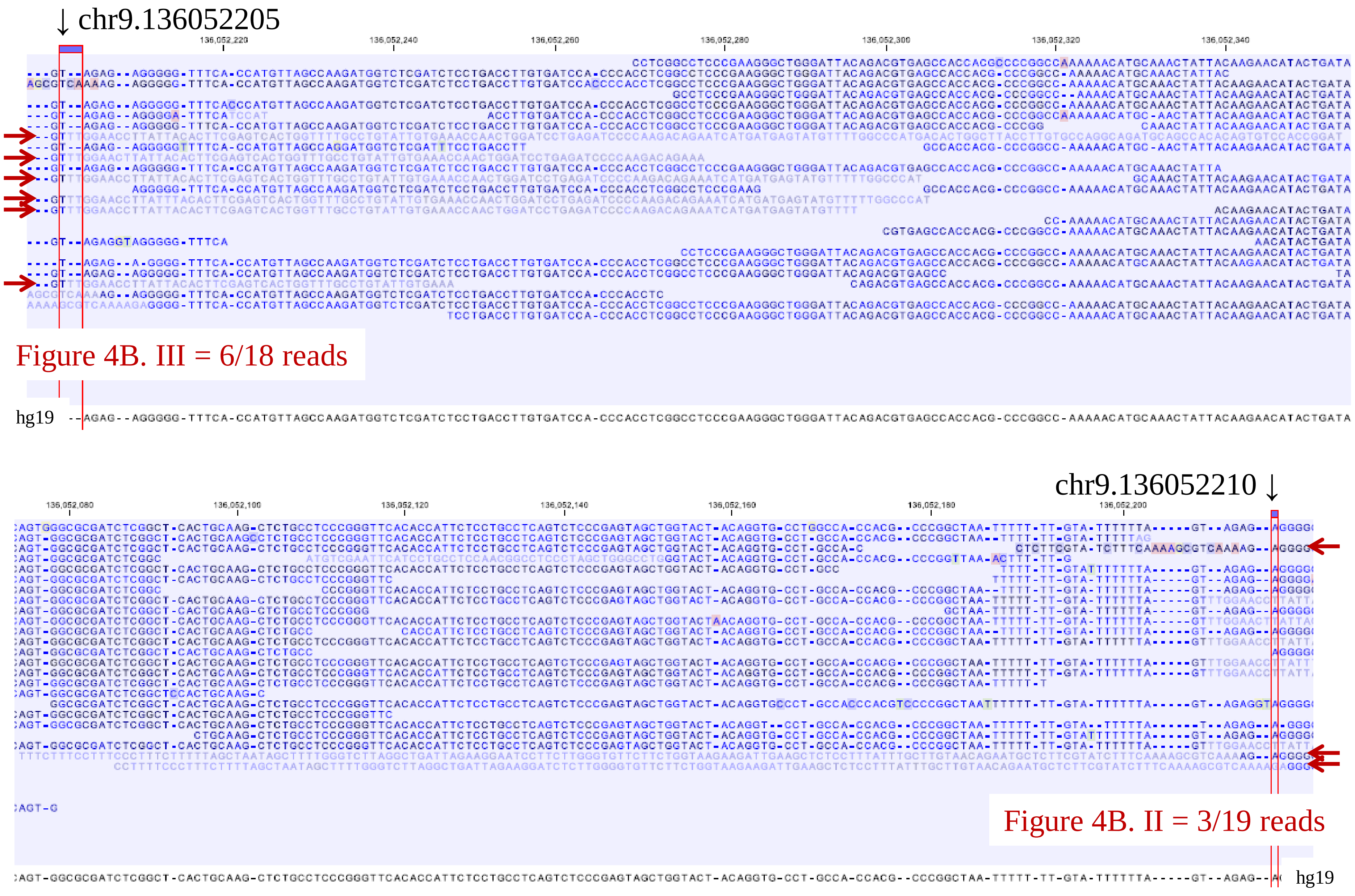

↓ chr9.136052205
Figure 4B. III = 6/18 reads
hg19
chr9.136052210 ↓
Figure 4B. II = 3/19 reads
hg19


Figure S3 C.

D

Region of chr9.136052205～136052210

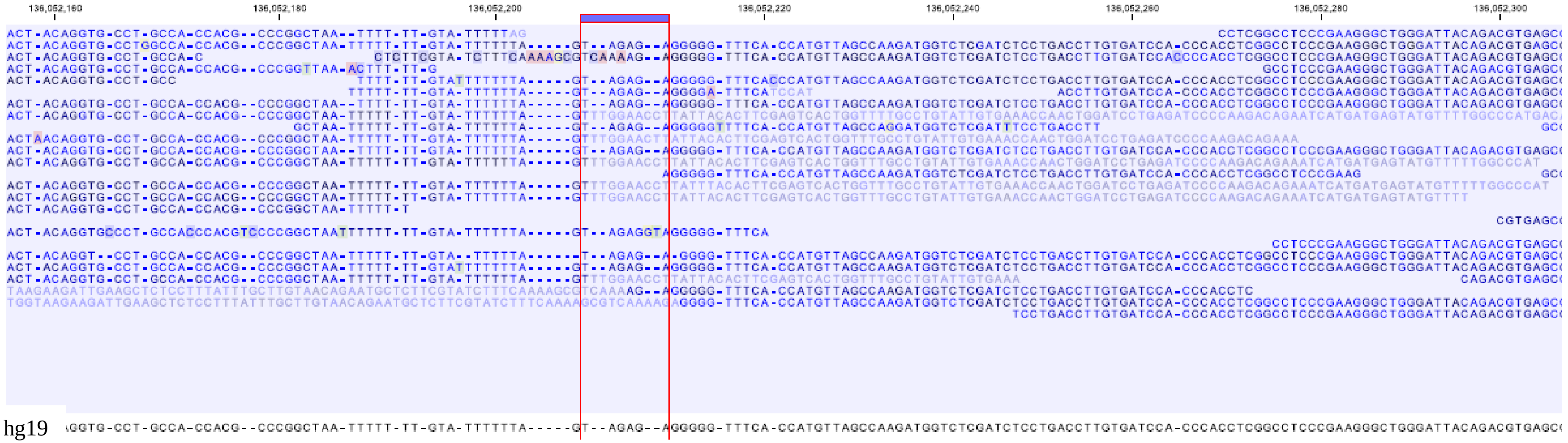


Figure S3 D.

E

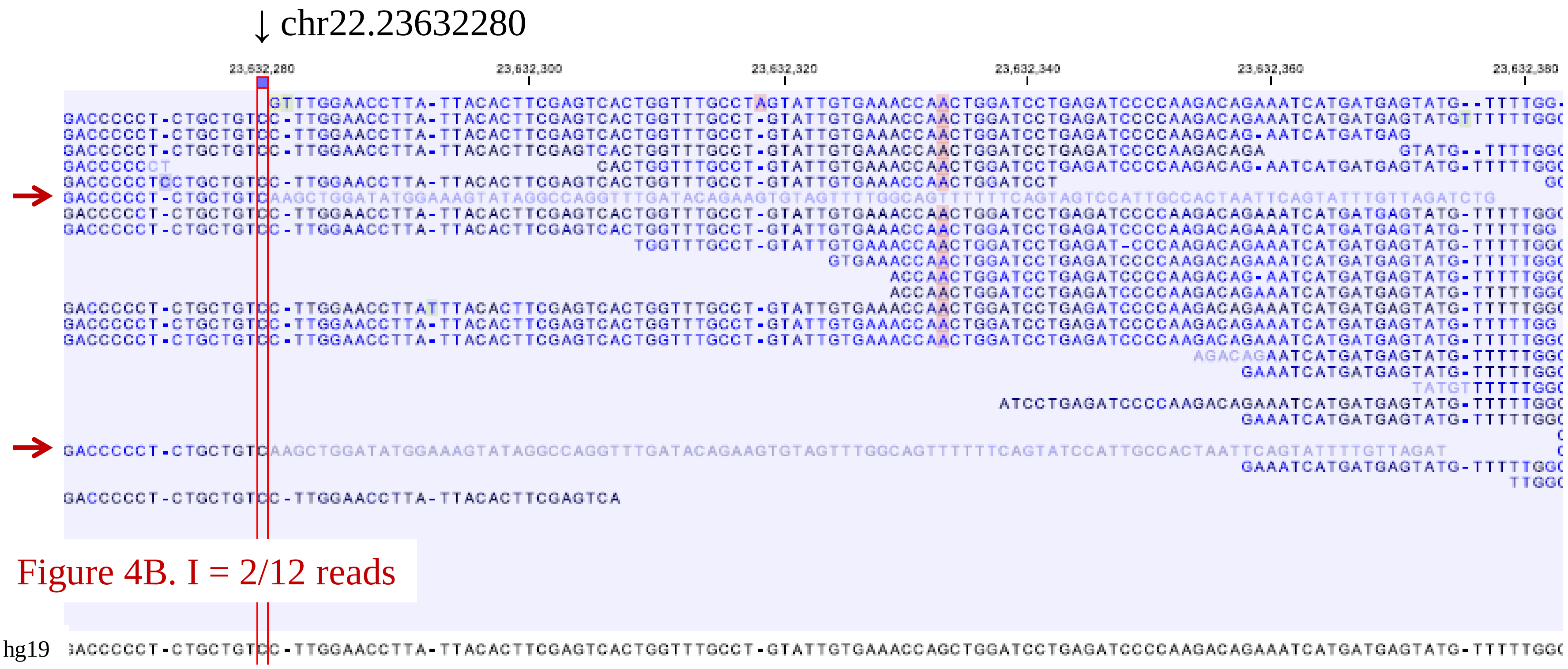
↓ chr22.23632280
23,632,280
23,632,300
23,632,320
23,632,340
23,632,360
23,632,380
Figure 4B. I = 2/12 reads
hg19

Figure S3 E.

F

Mean : 192 bases    Median : 214 bases    Mode : 222 bases

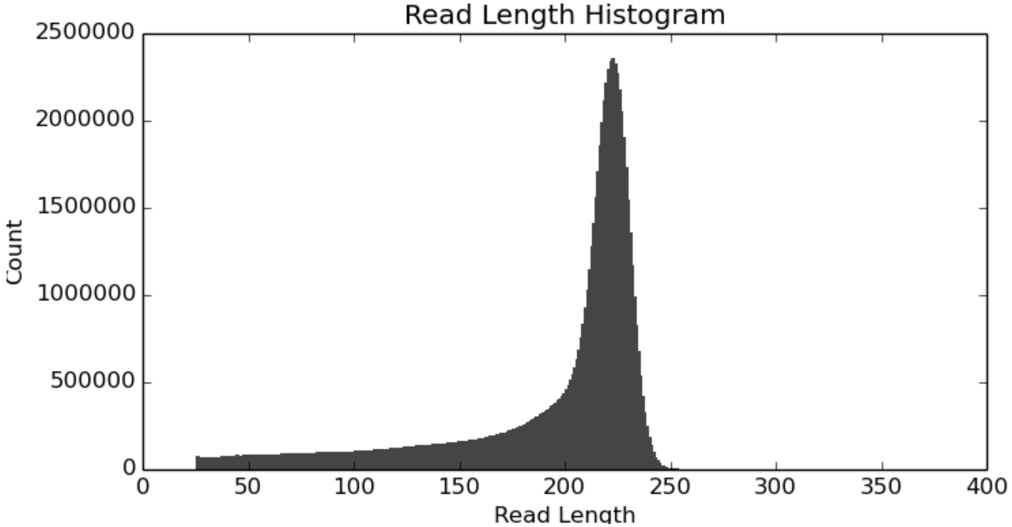


Using with:
Ion S5™ XL System
Ion Chef™ System
Ion 540™ Kit-Chef
(about 200 bases reads)
Ion 540™ Chip Kit

Library – about 230 bases insert

Mean : 293 bases    Median : 319 bases    Mode : 328 bases

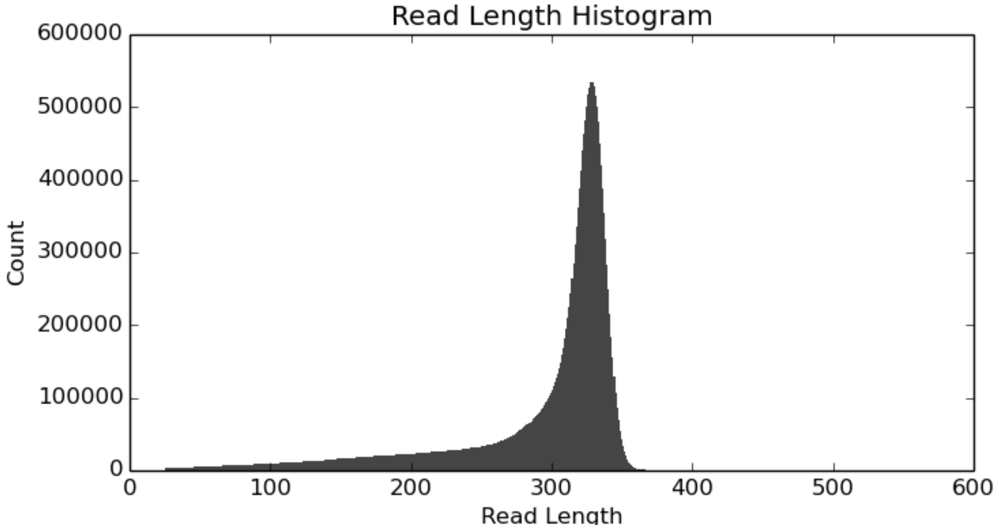


Using with:
Ion S5™ XL System
Ion Chef™ System
Ion 520™/530™ Kit-Chef
(～400 bases reads)
Ion 530™ Chip Kit

Library – about 350 bases insert

Figure S3 F.
