## Supplementary material for "ReALLEN: structural variation discovery in cancer genome by sensitive analysis of single-end reads": Figure S4

**Figure S4. Manual variation of SV calls.**
The mapped-reads of the CML sample are visualized by Integrative Genomics Viewer [31, 32] around the SVs in table S7. The description of the SV is displayed at the top of each panel. Soft-clipped bases are rendered visible. The deletions less than 200 bp are displayed with one view, and the other SVs are displayed with two views. The top of the view shows the position. The middle of the view shows the coverage (upper side) and the mapped reads (lower side). The bottom of the view shows the reference genome (hg19). Our comments are written in red text in panels where the calls have been judged FP by manual validation.

### #1, chrX: 150203866-150203894, DEL (27 bp), validation: TP

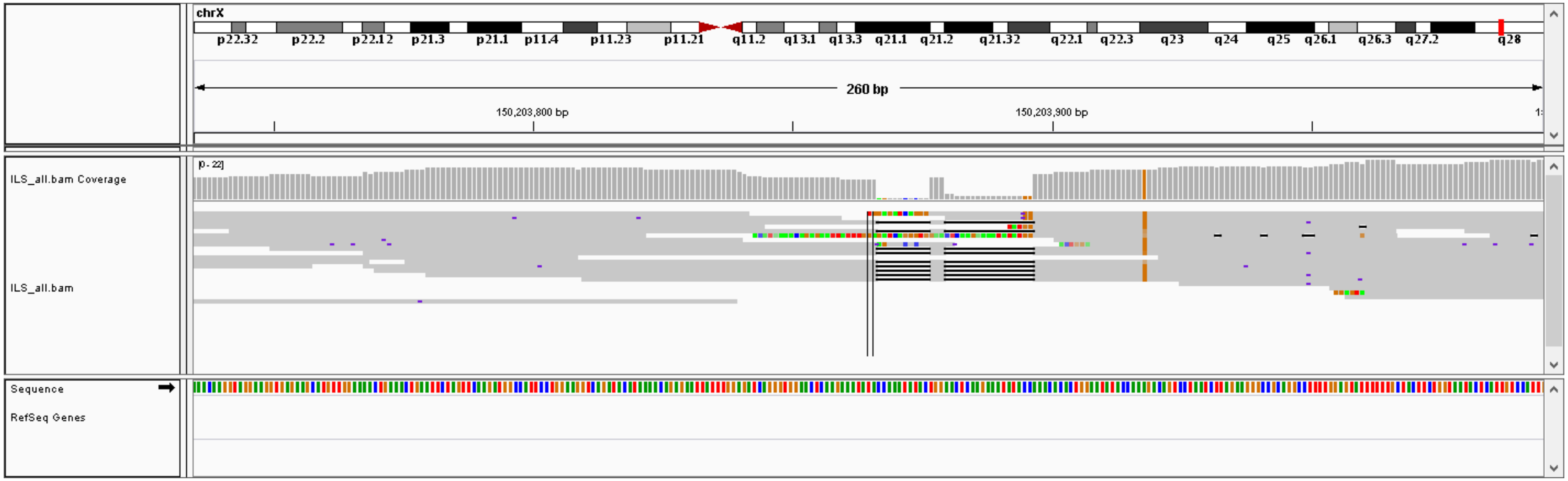

### #2, chr2: 81604891-81604928, DEL (36 bp), validation: TP

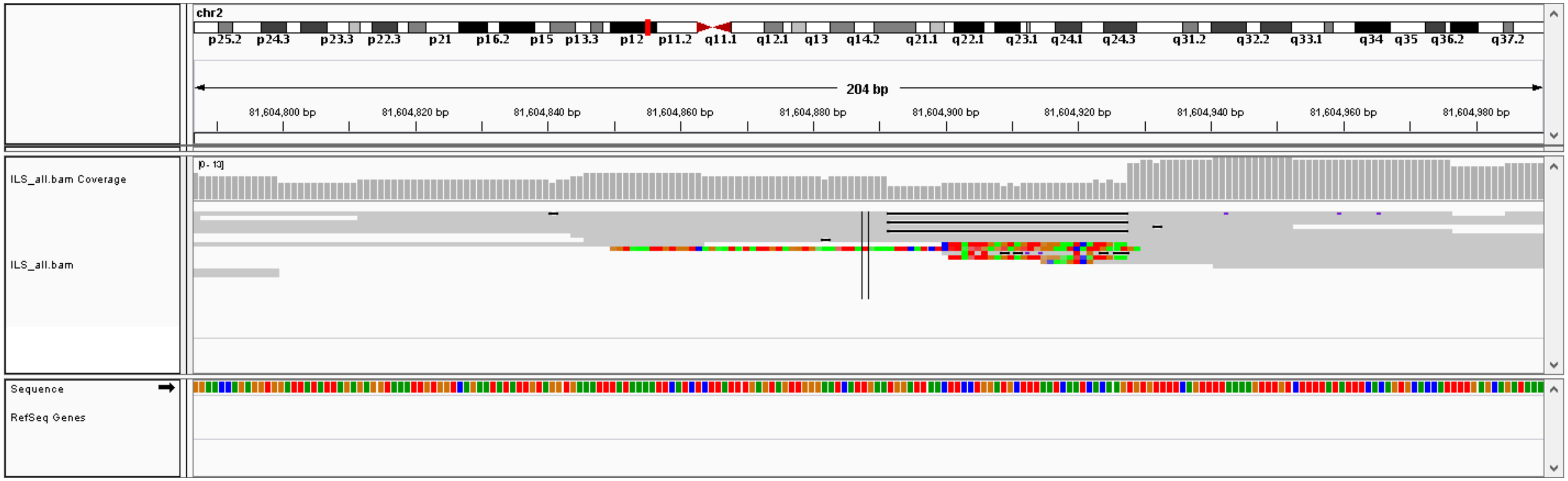

### #3, chrX: 138384894-138385221, DEL (326 bp), validation: TP

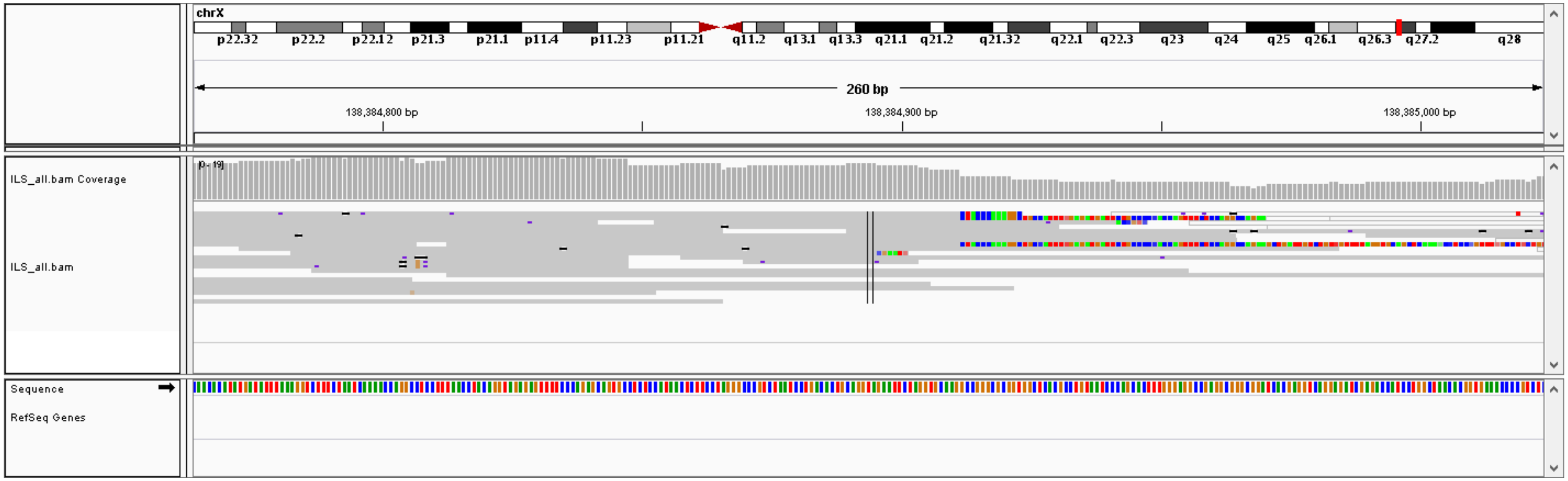


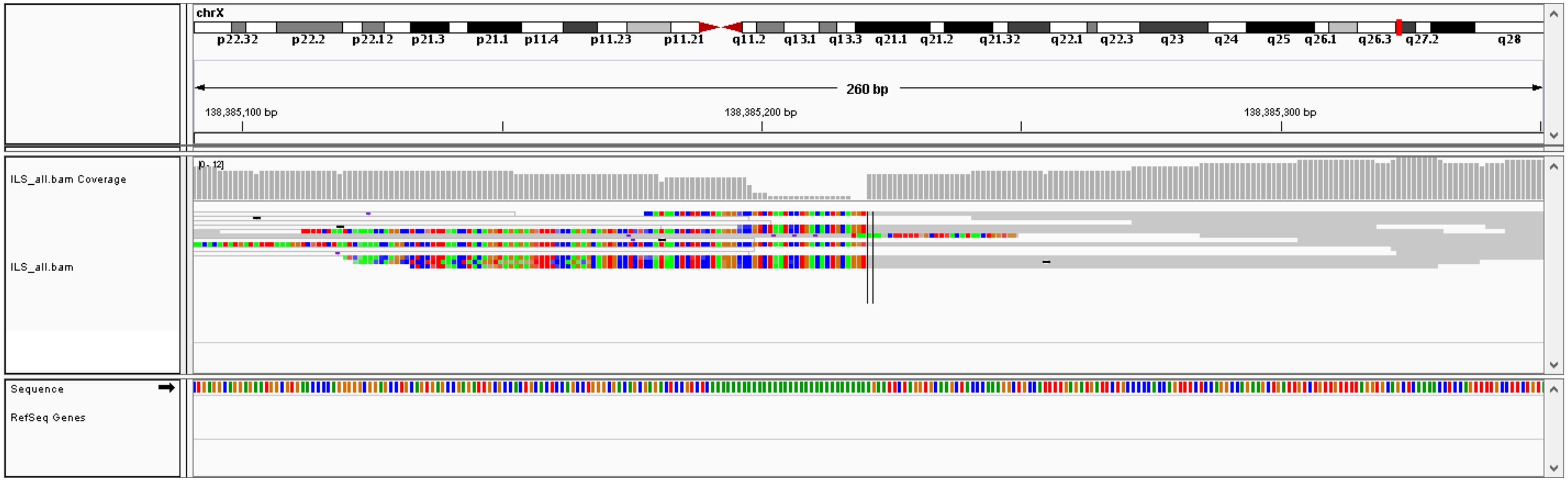

#4, chr7: 51822314-51822355,  DEL (40 bp), validation: TP

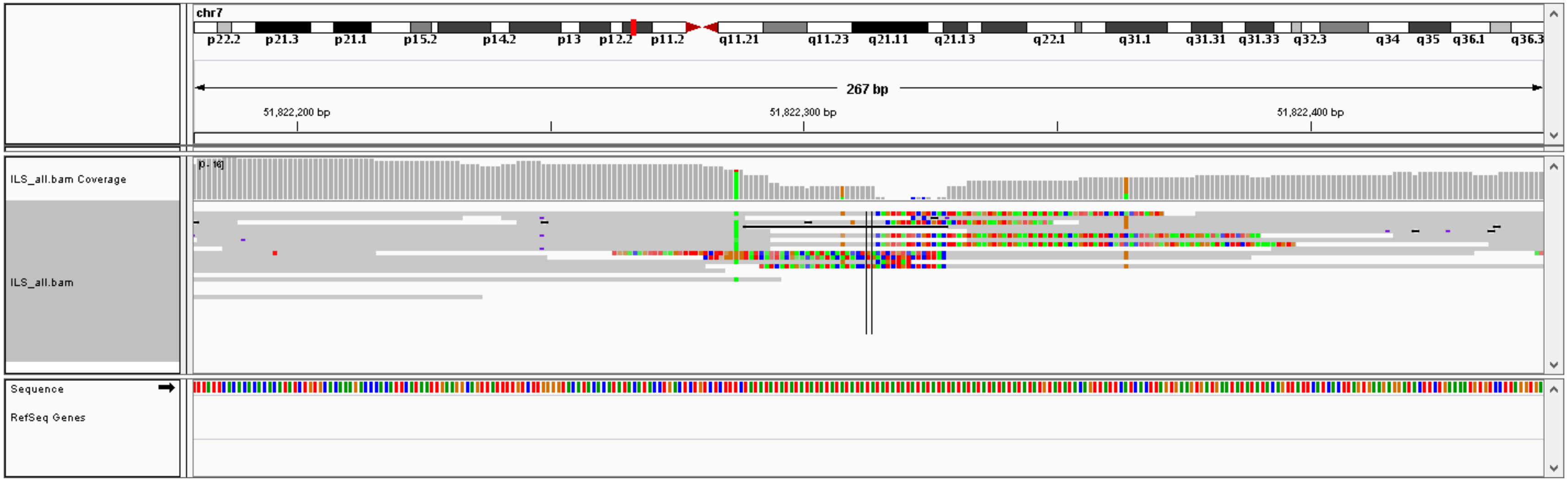

### #5, chr13: 30578140-30578179, DEL (38 bp), validation: TP

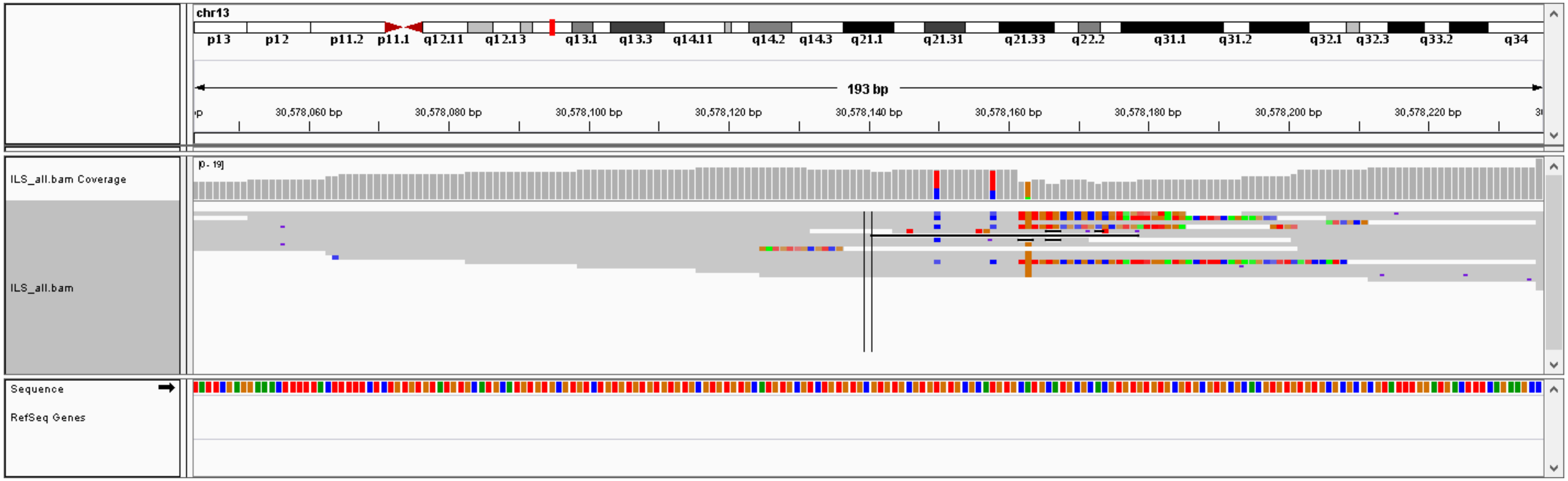

### #6, chr22: 32887583-32887612,  DEL (28 bp), validation: TP

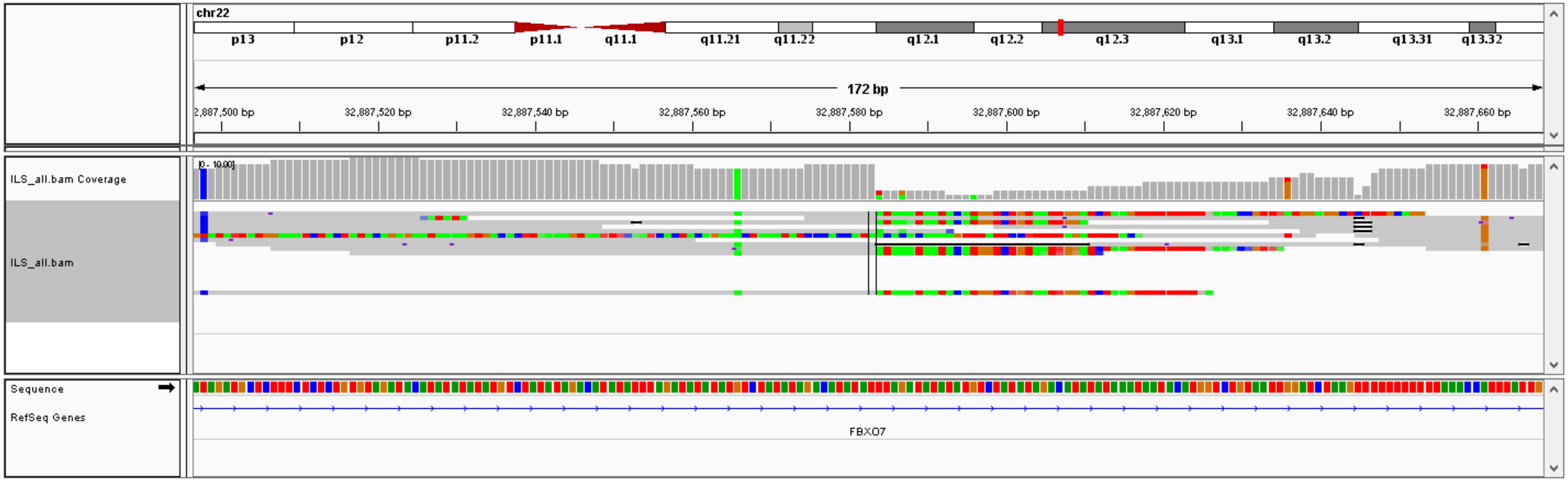

### #7, chr12: 29280251-29280568,  DEL (316 bp), validation: TP

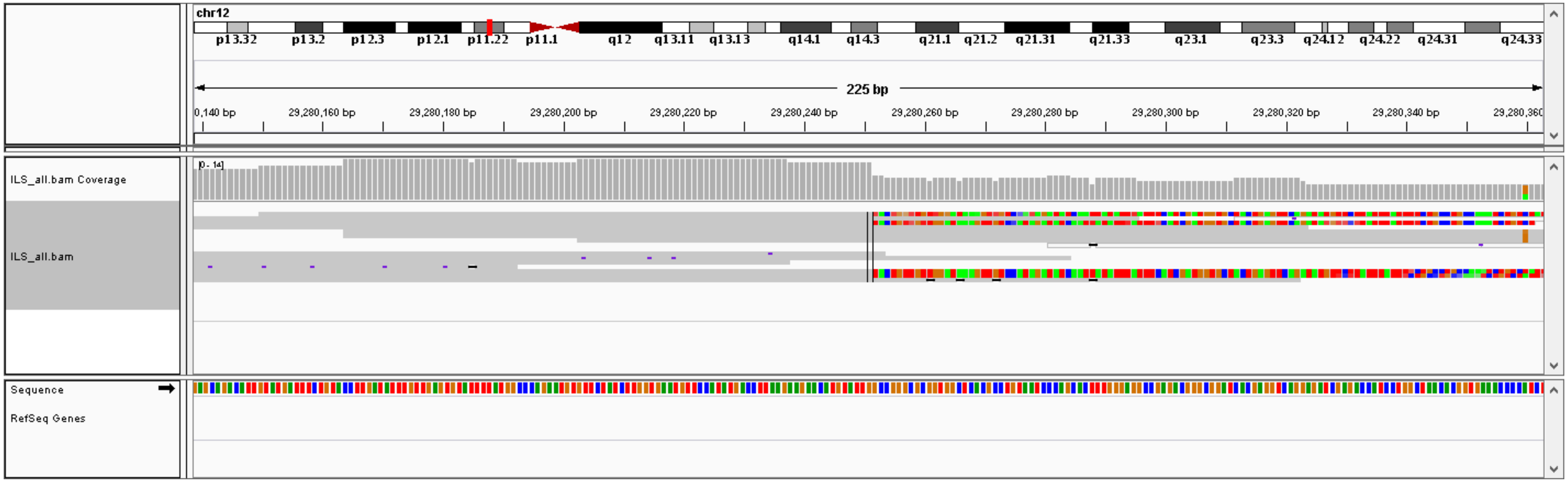


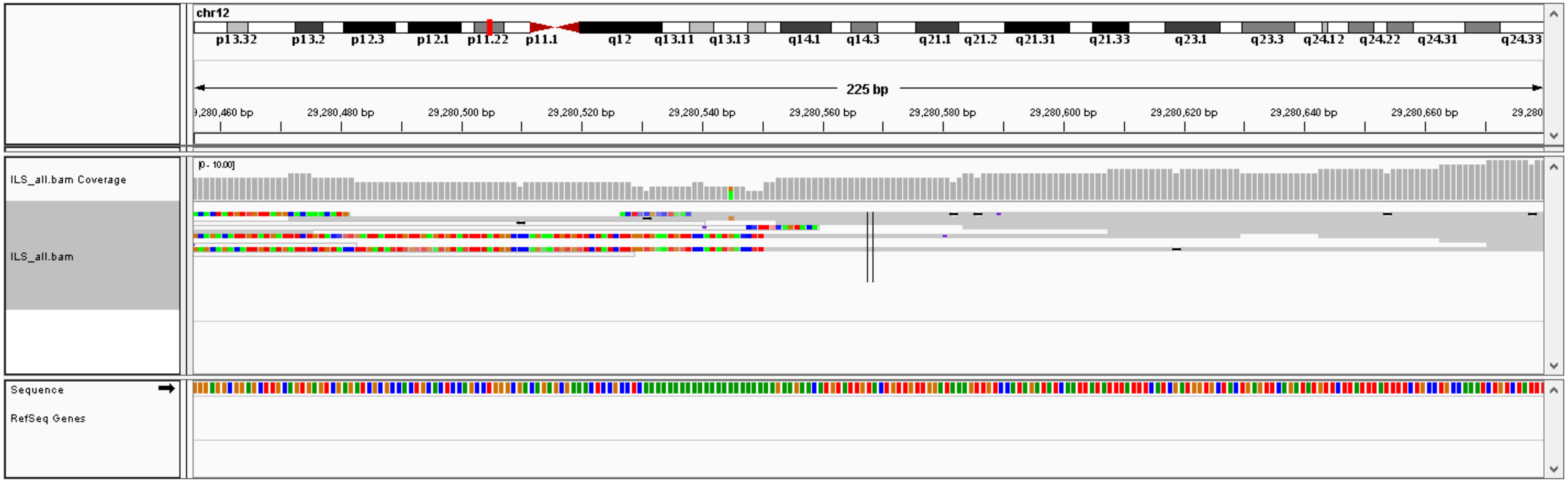

### #8, chr5: 167334525-167334551,  DEL (25 bp), validation: TP

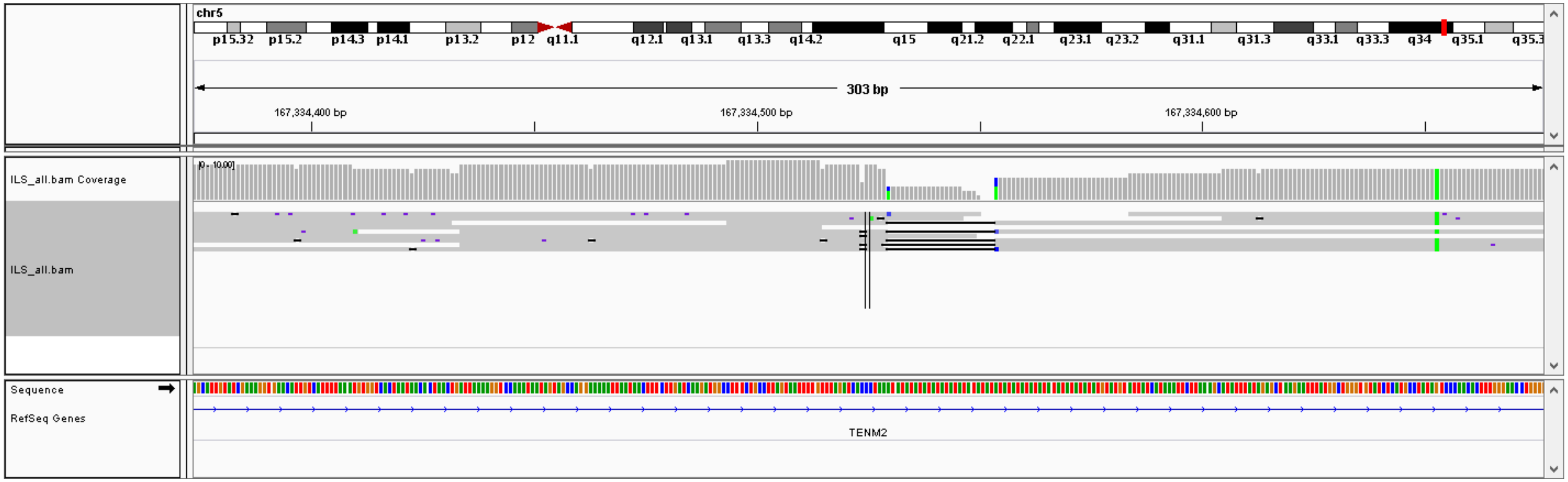

### #9, chr3: 136021013-136026199,  DEL (5185 bp), validation: TP

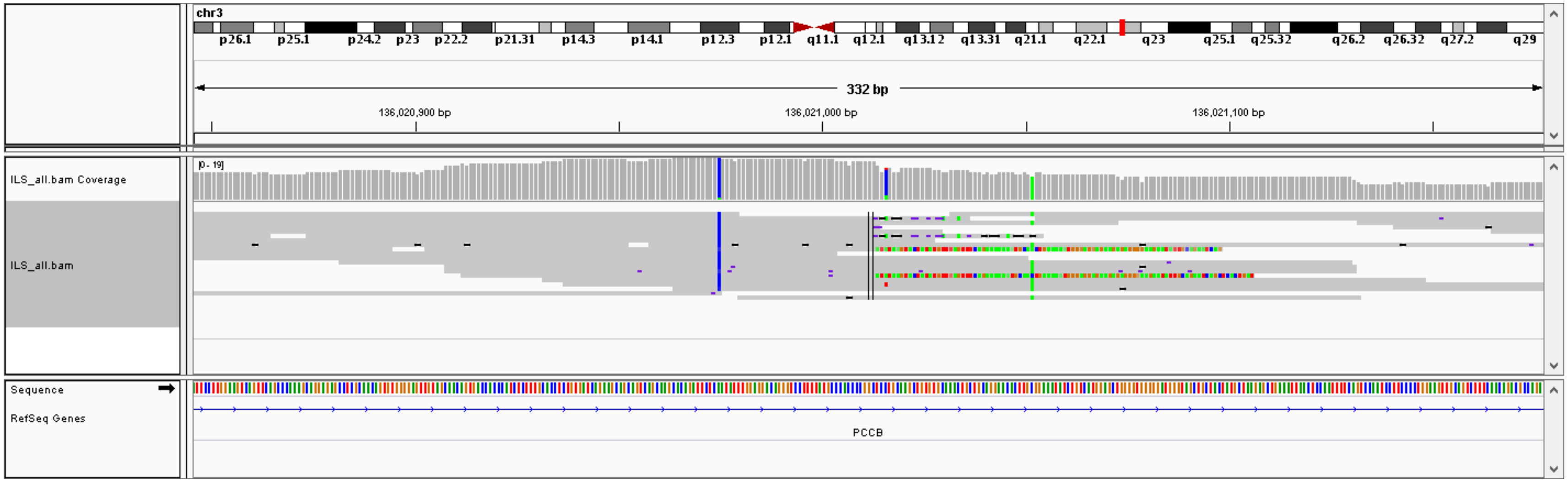


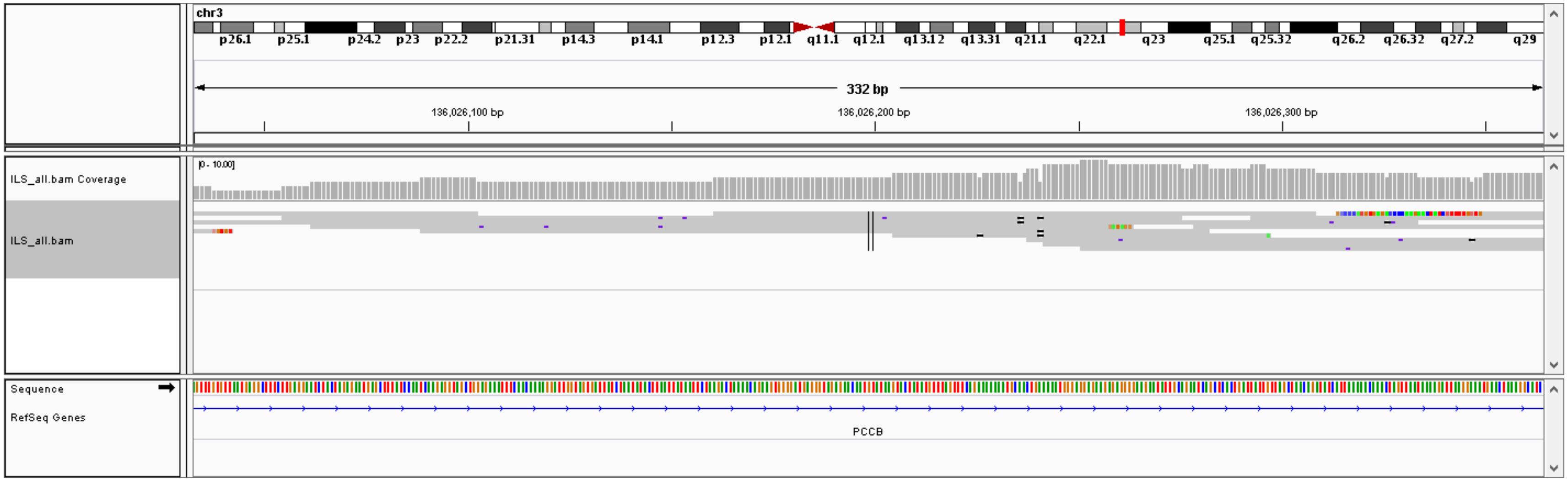

### #10, chr1: 189032968-189033030,  DEL (61 bp), validation: TP

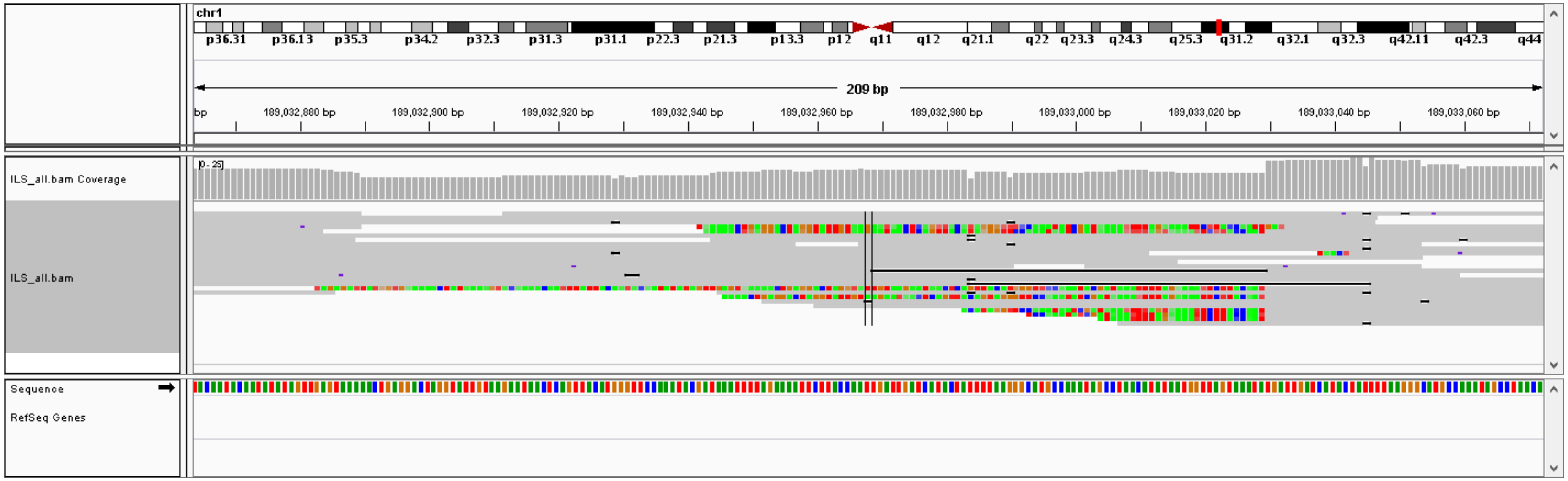

#11, chrX: 12842519- chr19: 28570792,  CTX, validation: TP

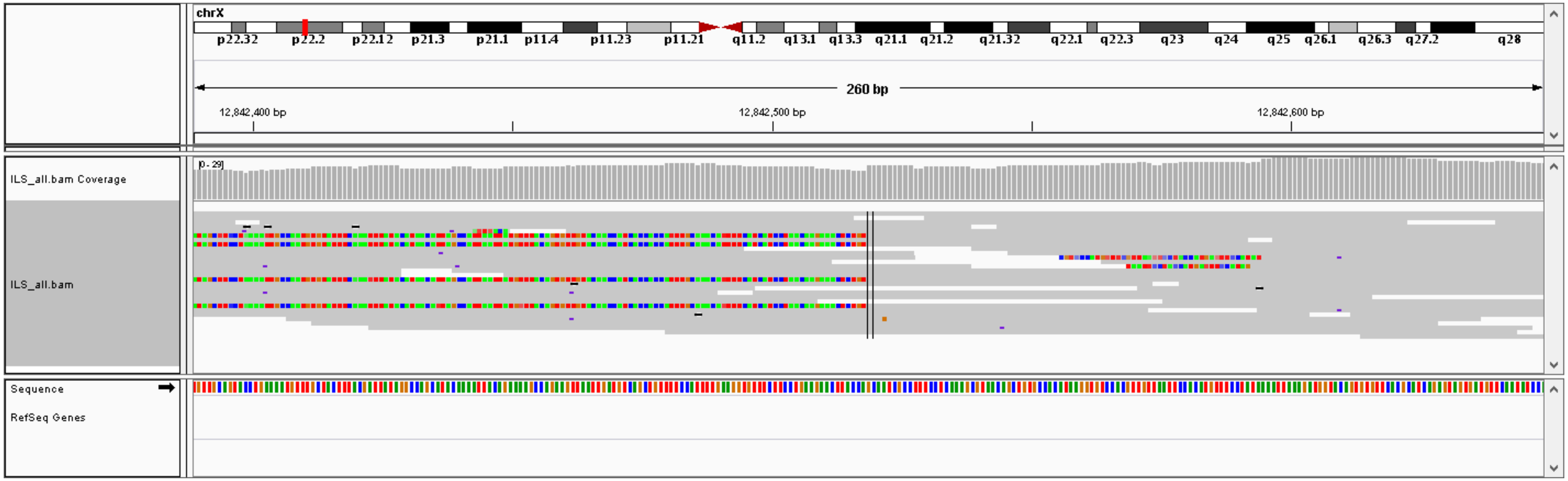


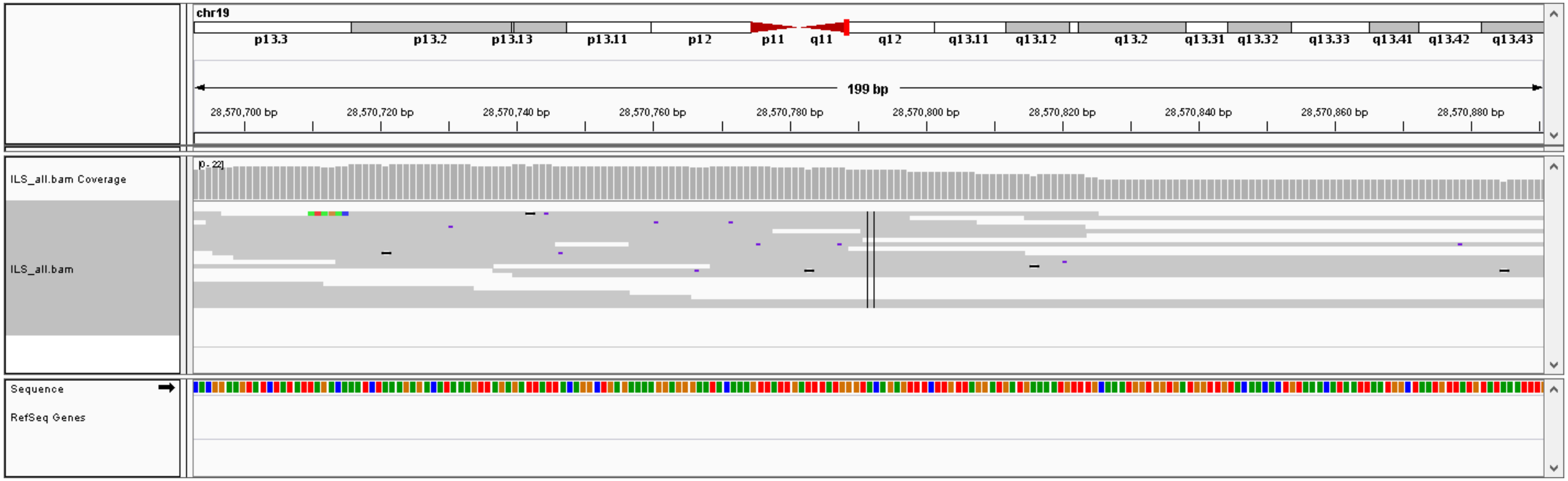

### #12, chr9: 9557895- chr18: 34206962,  CTX, validation: TP

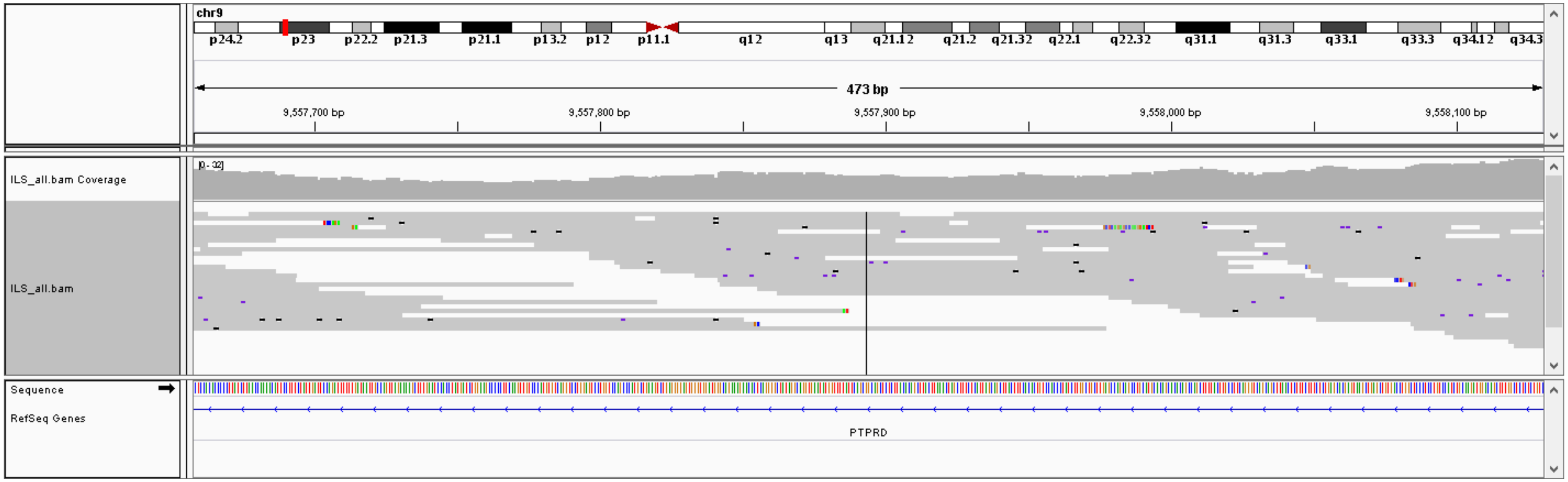


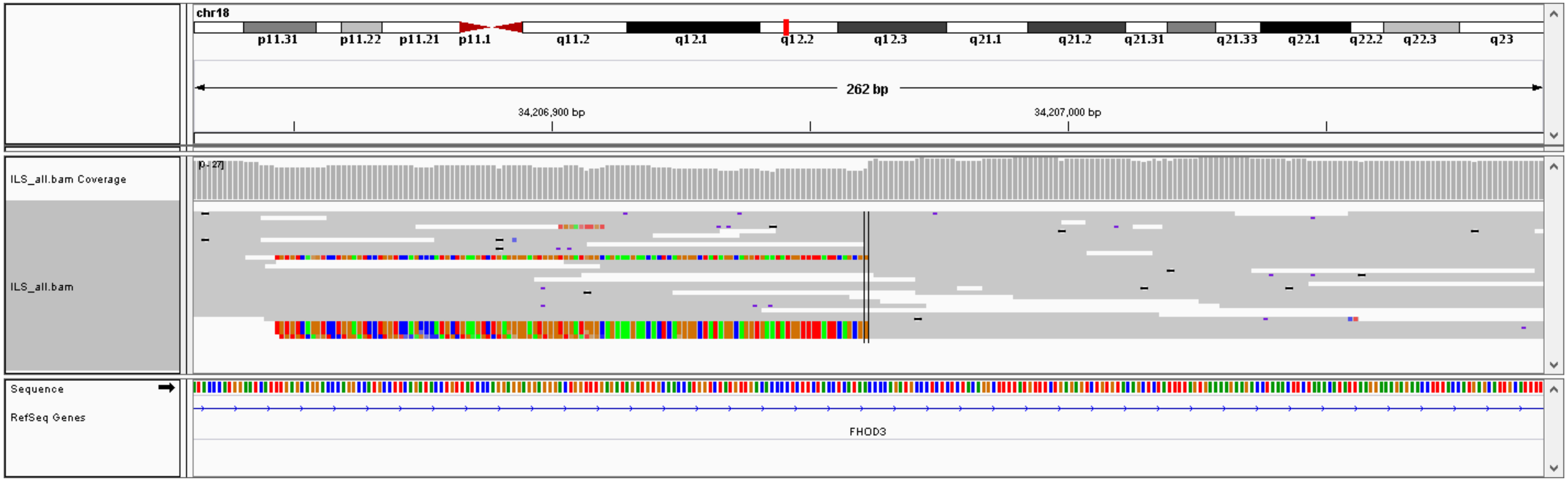

### #13, chr6: 83650368- chr2: 2617192,  CTX, validation: TP

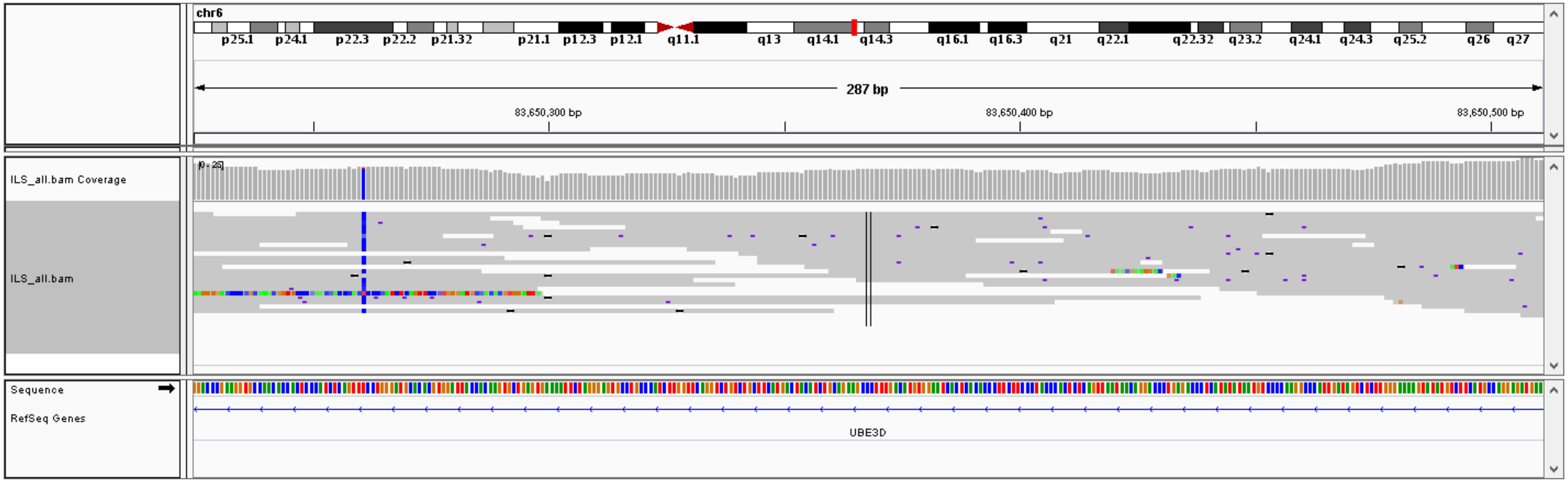


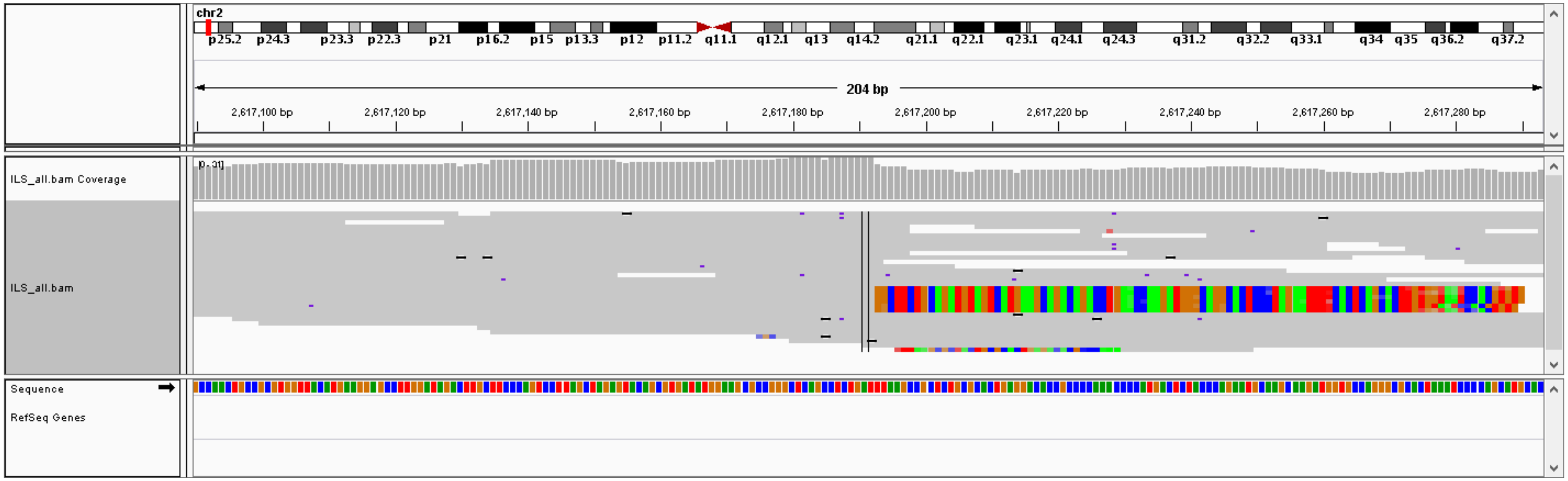

### #14, chr7: 128900405- chr10: 2990717, CTX, validation: TP

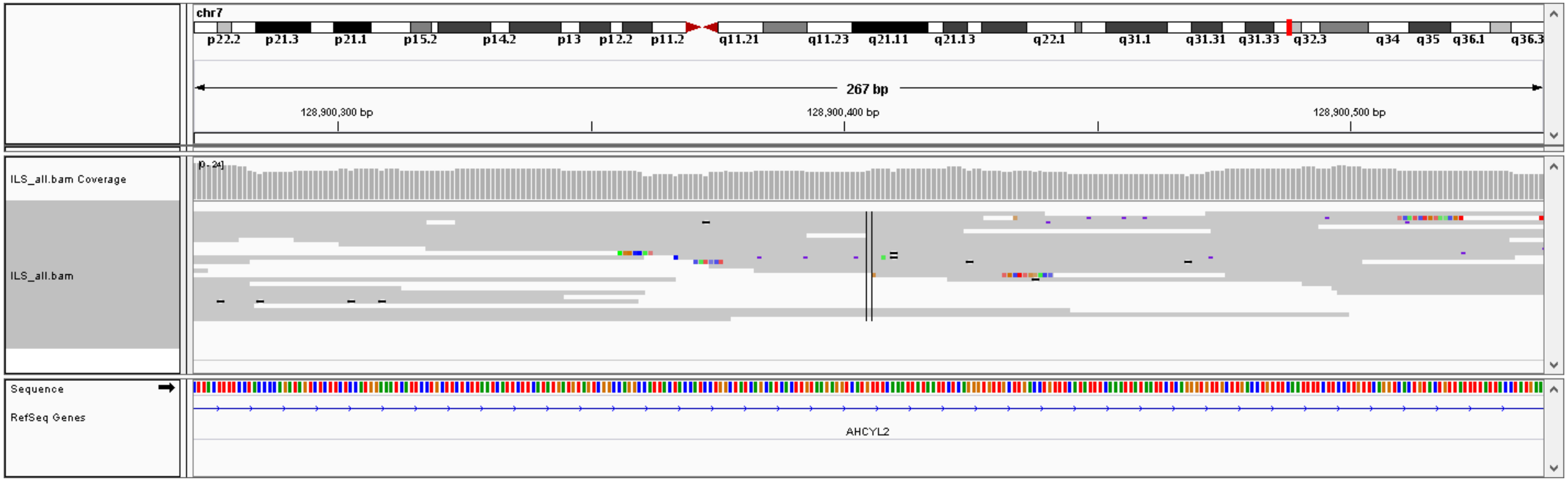


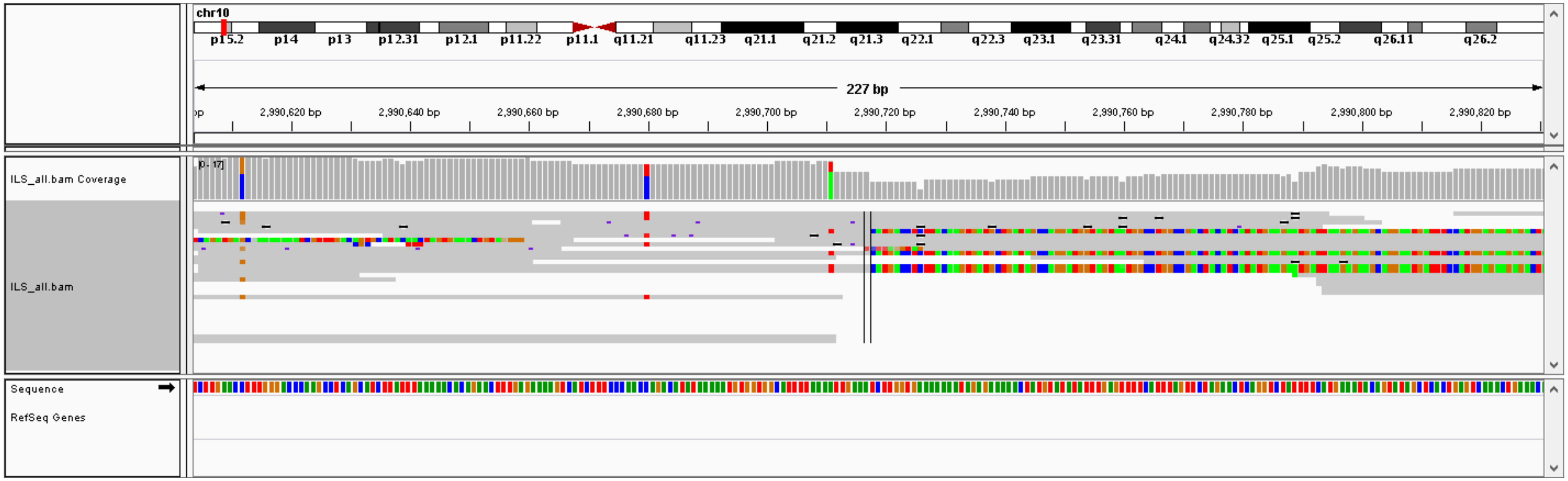

### #15, chr8: 104822514- chr1: 16431681,  CTX, validation: TP

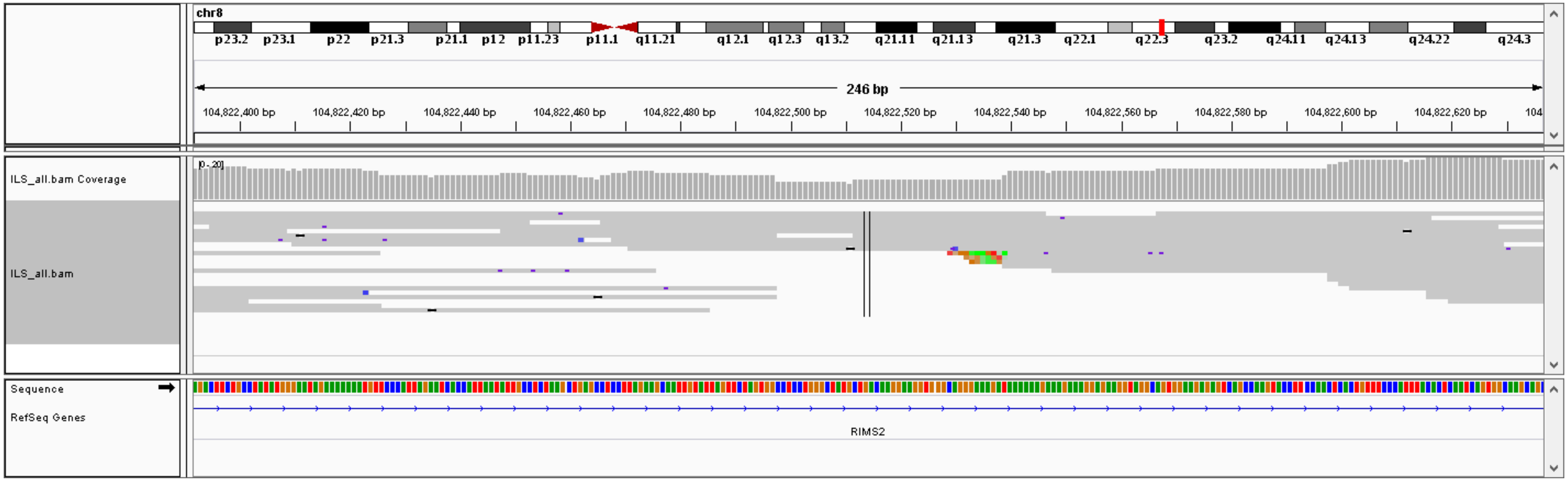


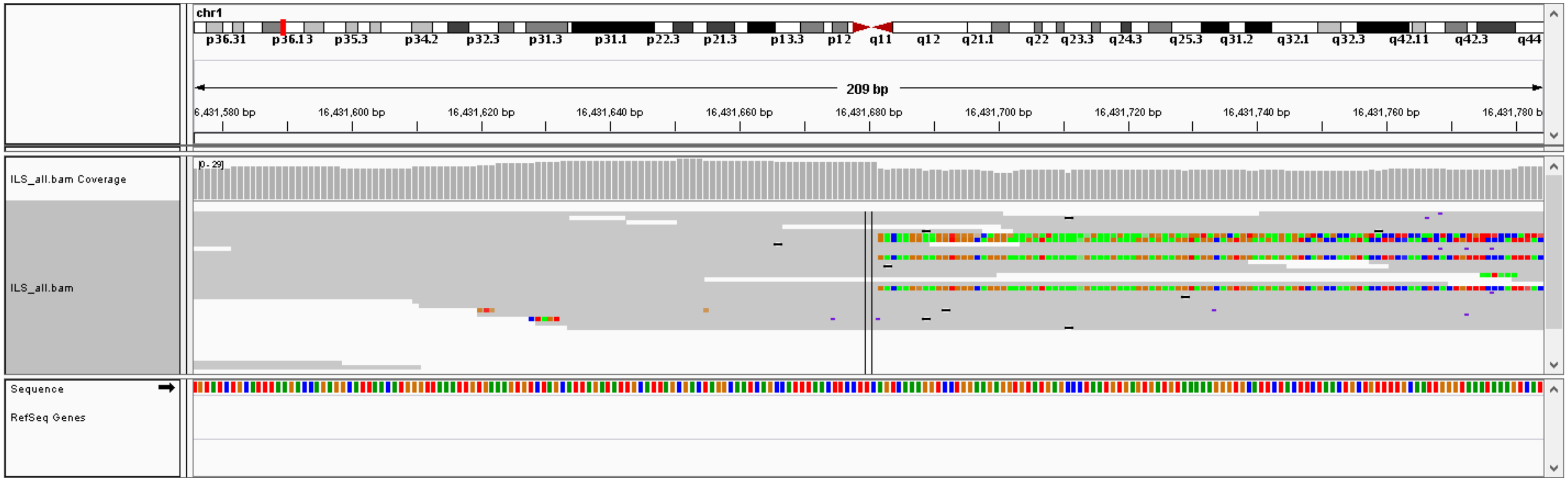

### #16, chr5: 77227895- chr14: 54446184,  CTX, validation: TP

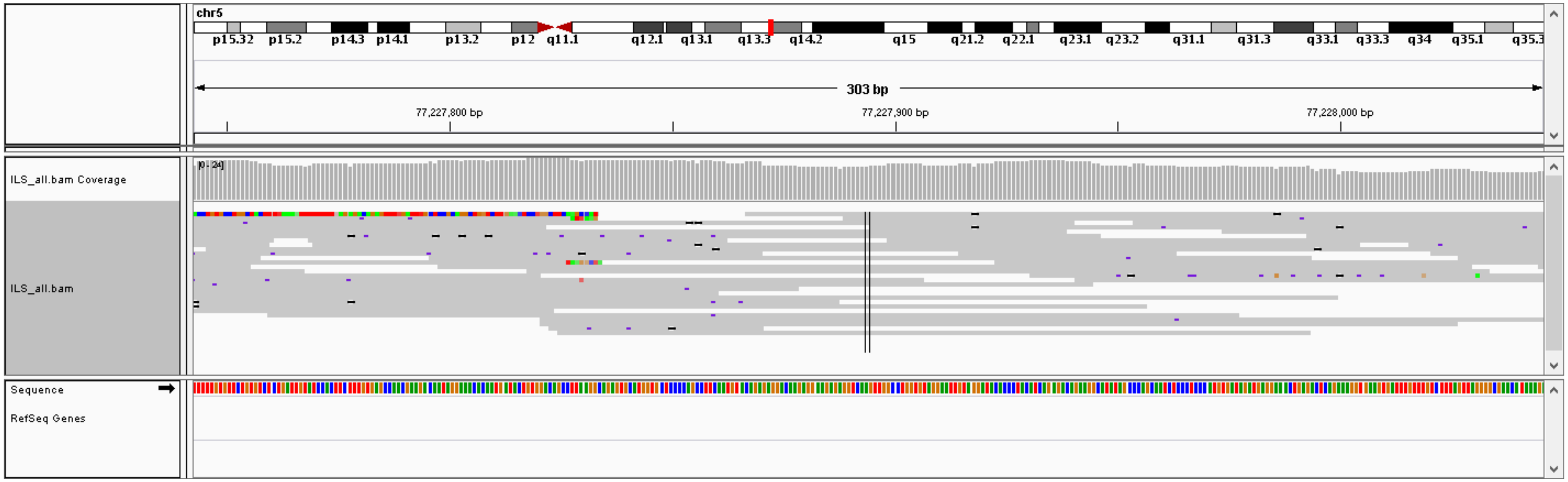


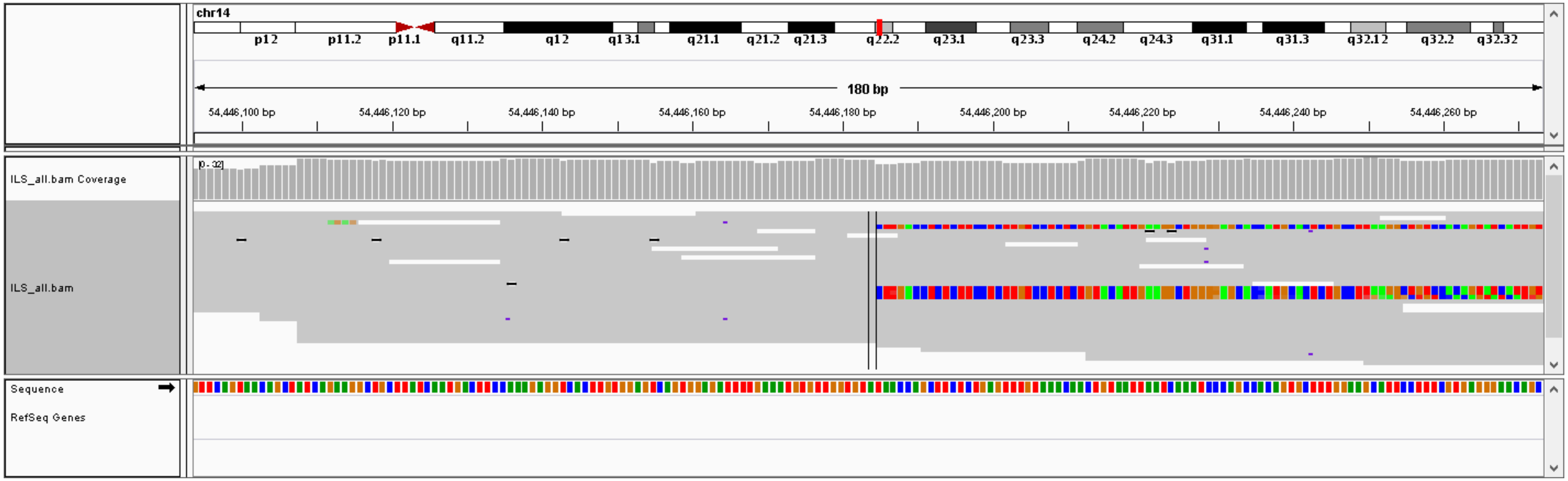

### #17, chr6: 57575919- chr5: 21573435,  CTX, validation: TP

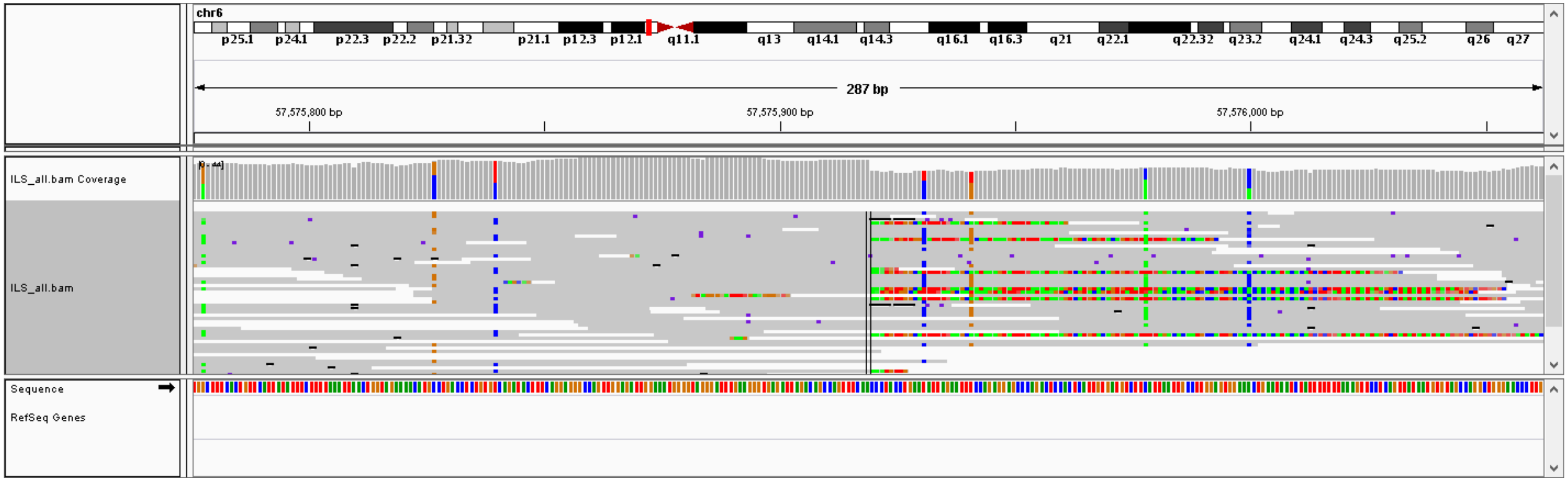


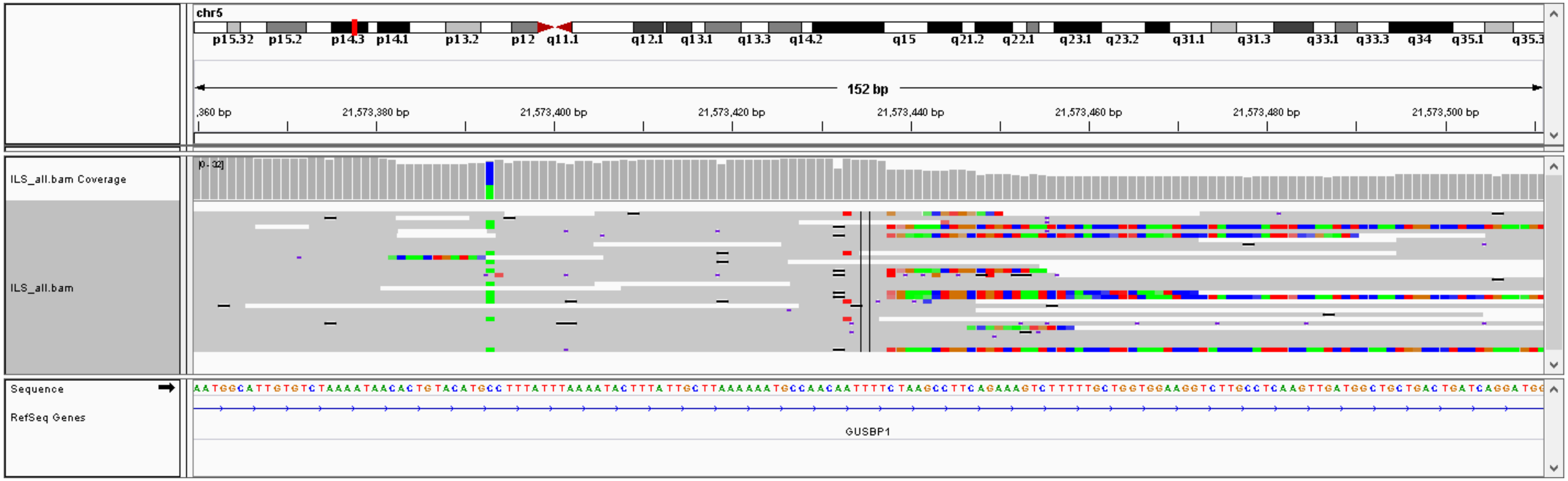

#18, chr9: 68418844- chr1: 80168979,  CTX, validation: FP

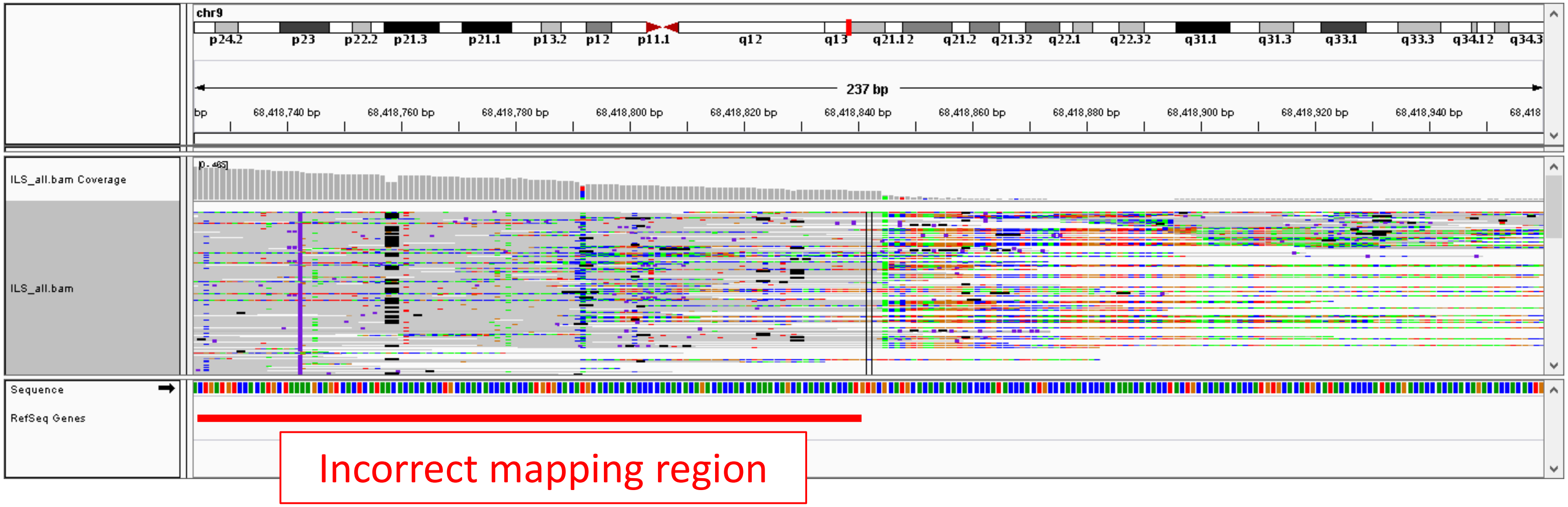


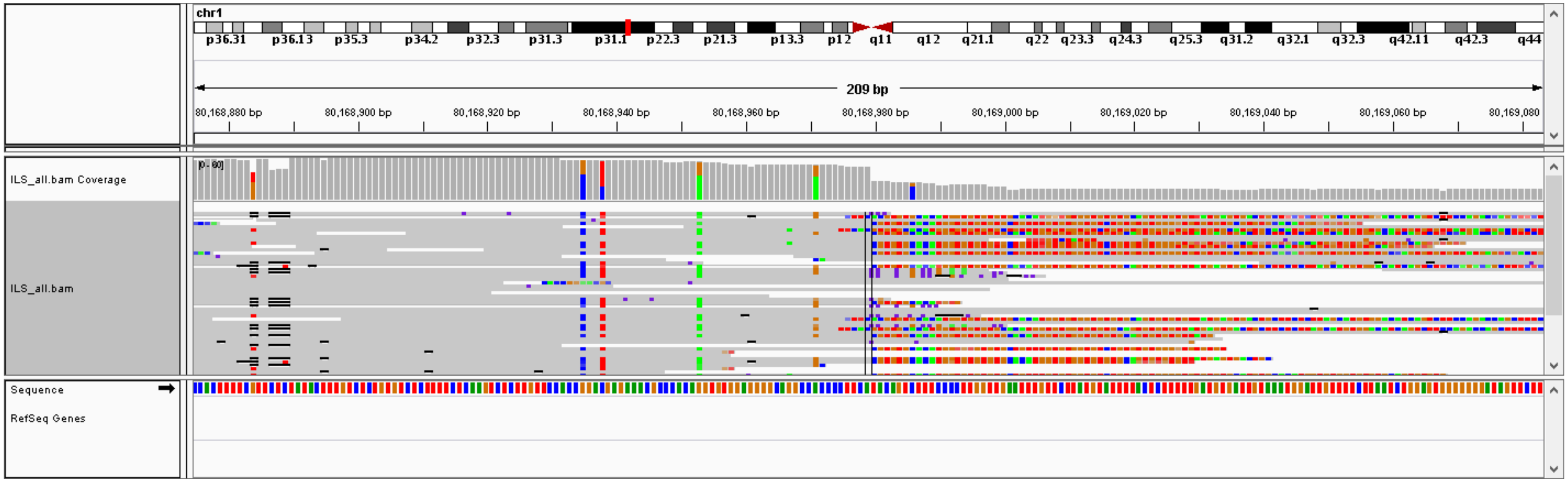

### #19, chr11: 113077144- chr1: 226814662,  CTX, validation: TP

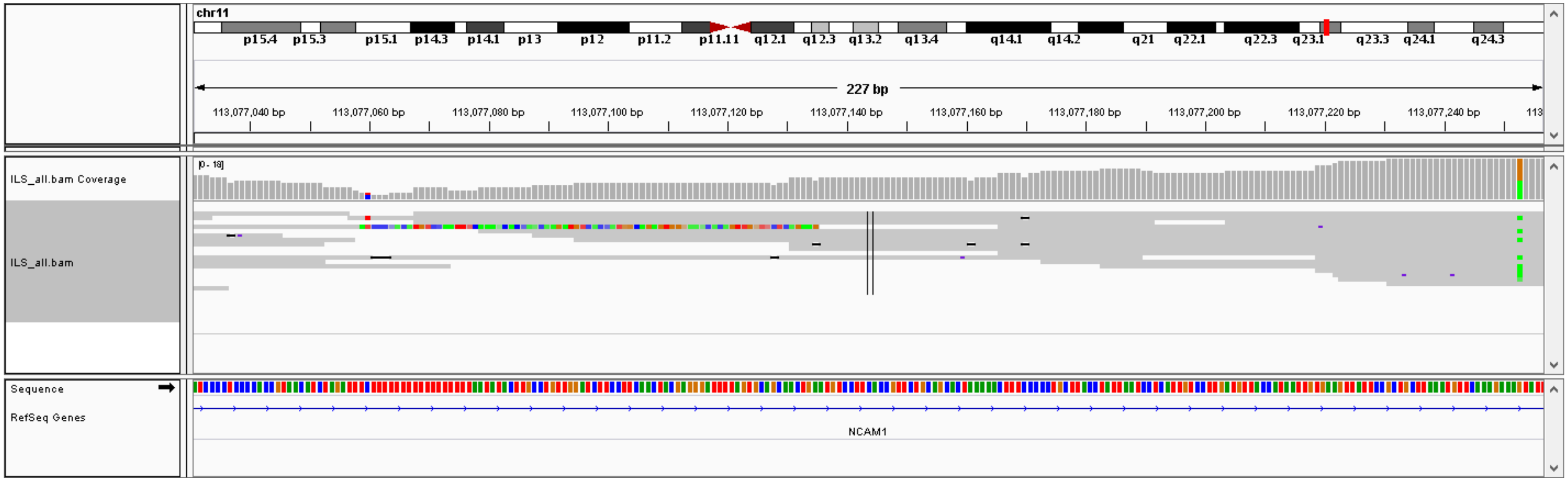


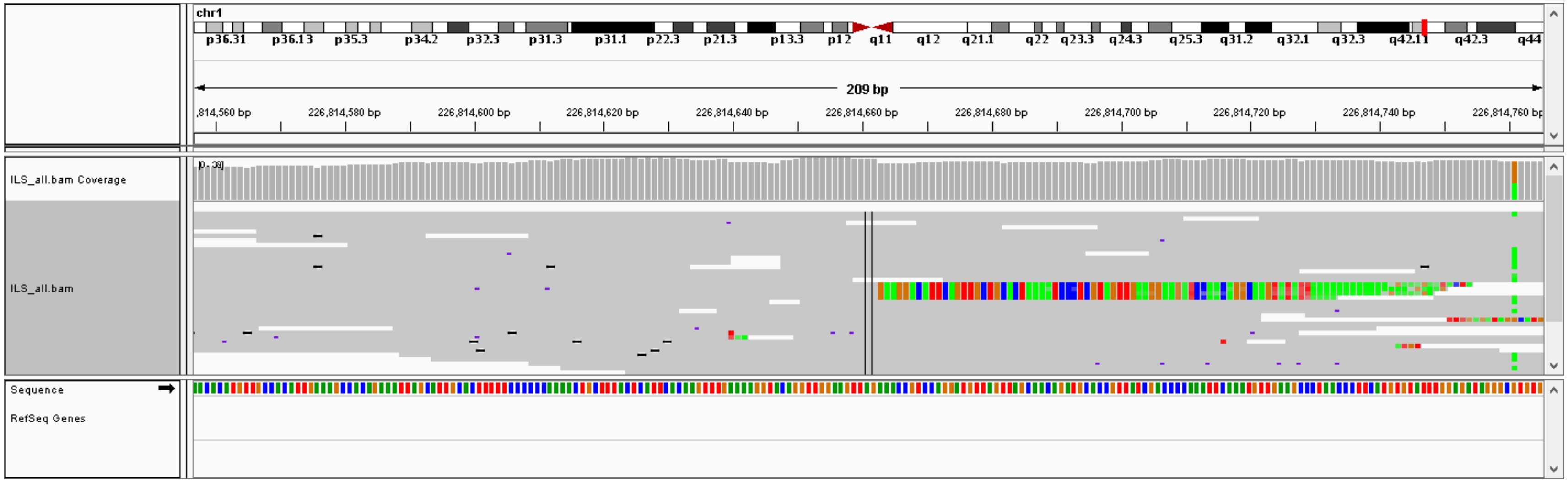

#20, chrM: 16536- chr17: 22020694,  CTX, validation: FP

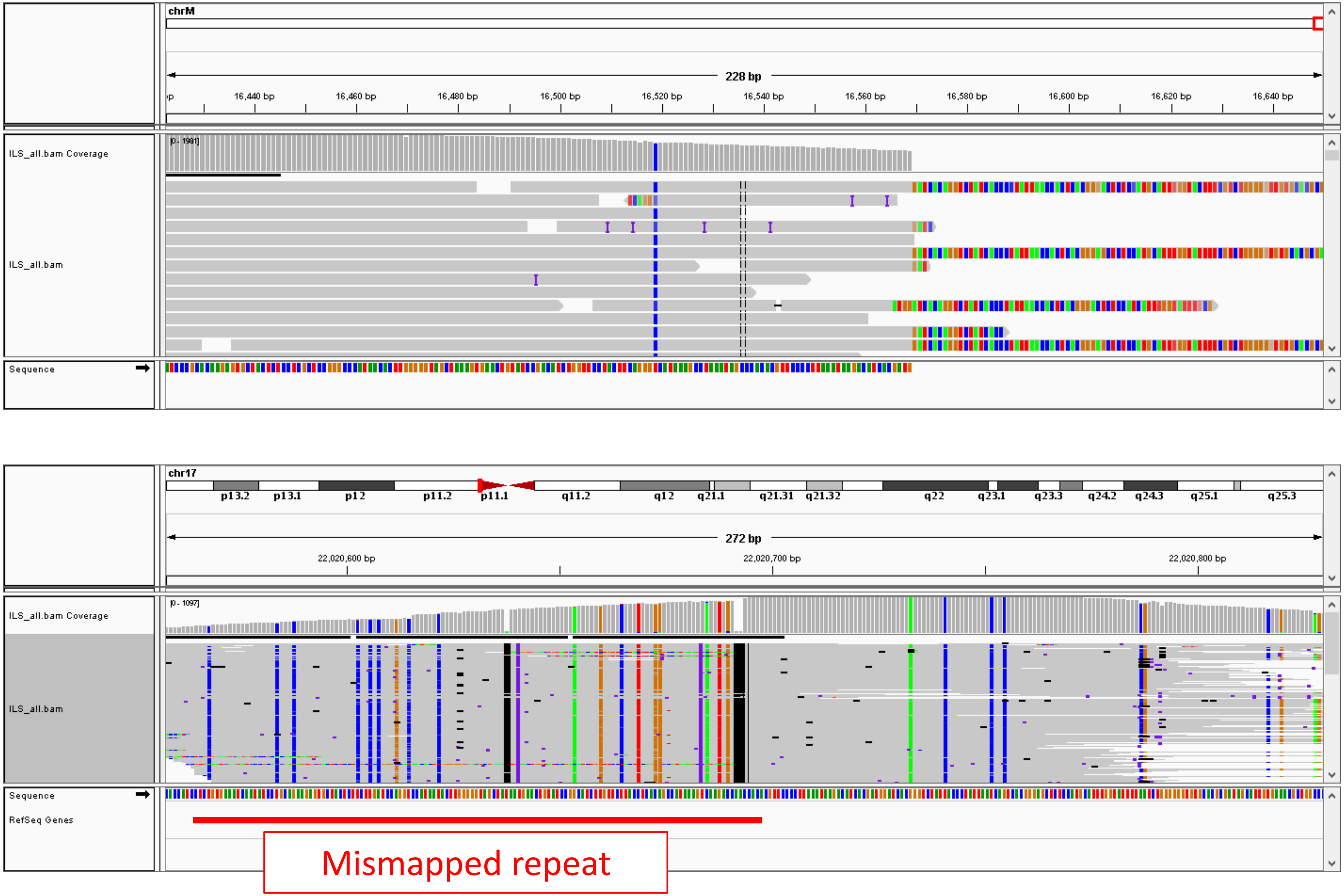

### #21, chr1: 60048636-60049662,  DEL (1025 bp), validation: TP

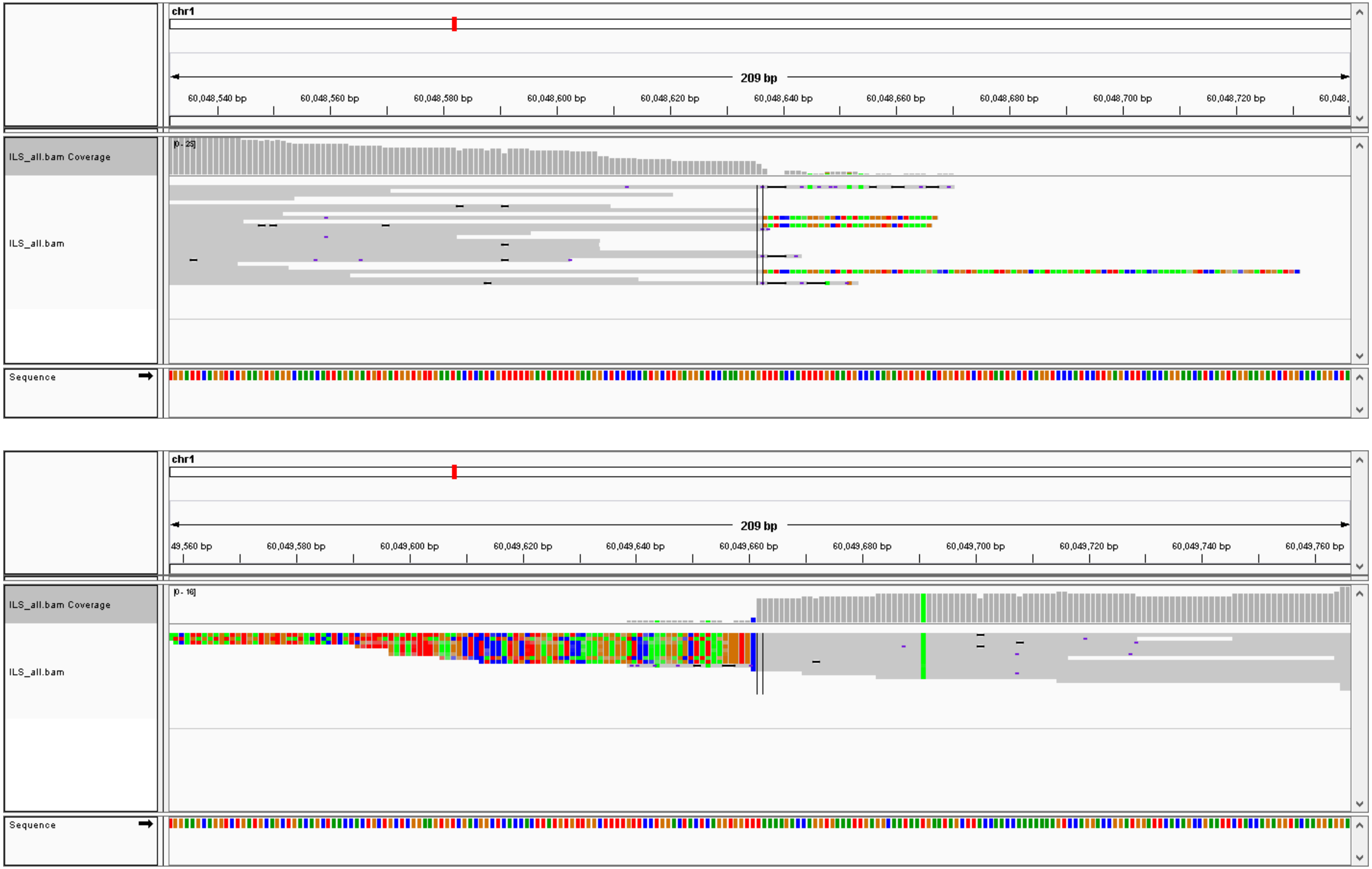

### #22, chr4: 9576977-9577449, DEL (471 bp), validation: TP

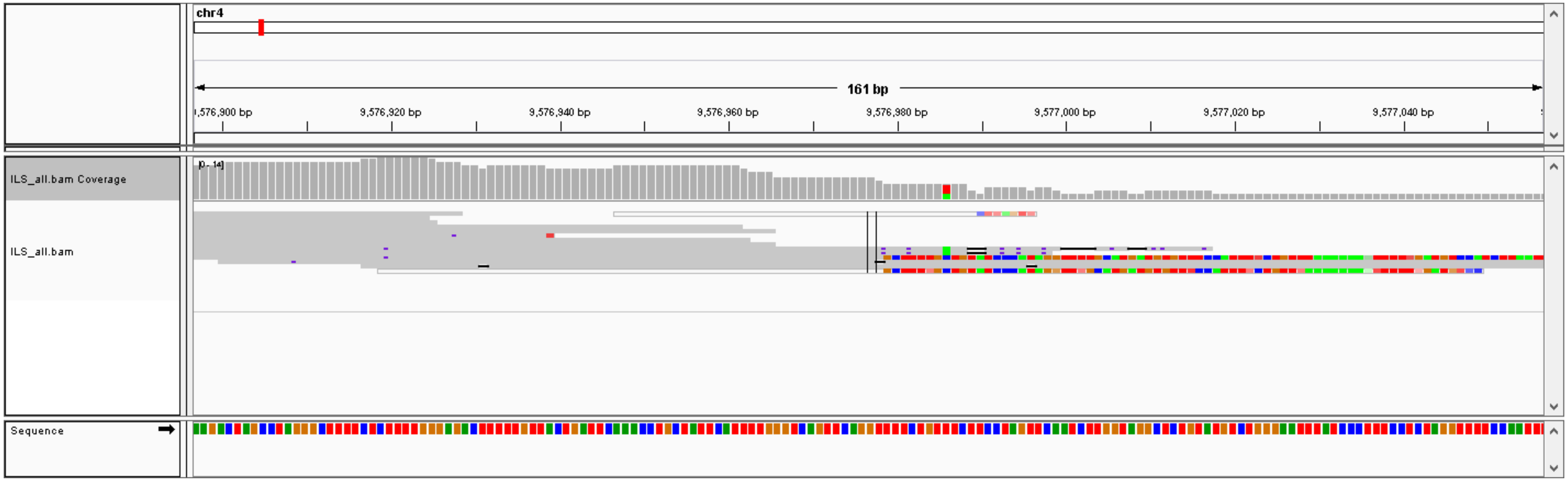


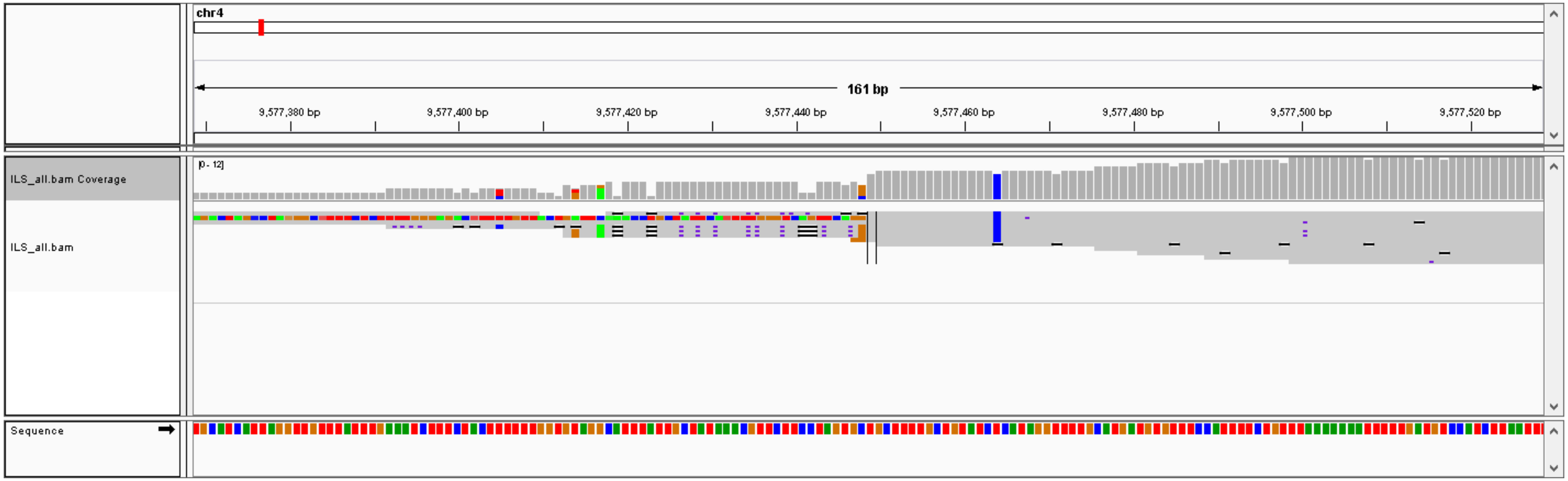

### #23, chr3: 158275628-158275730,  DEL (101 bp), validation: TP

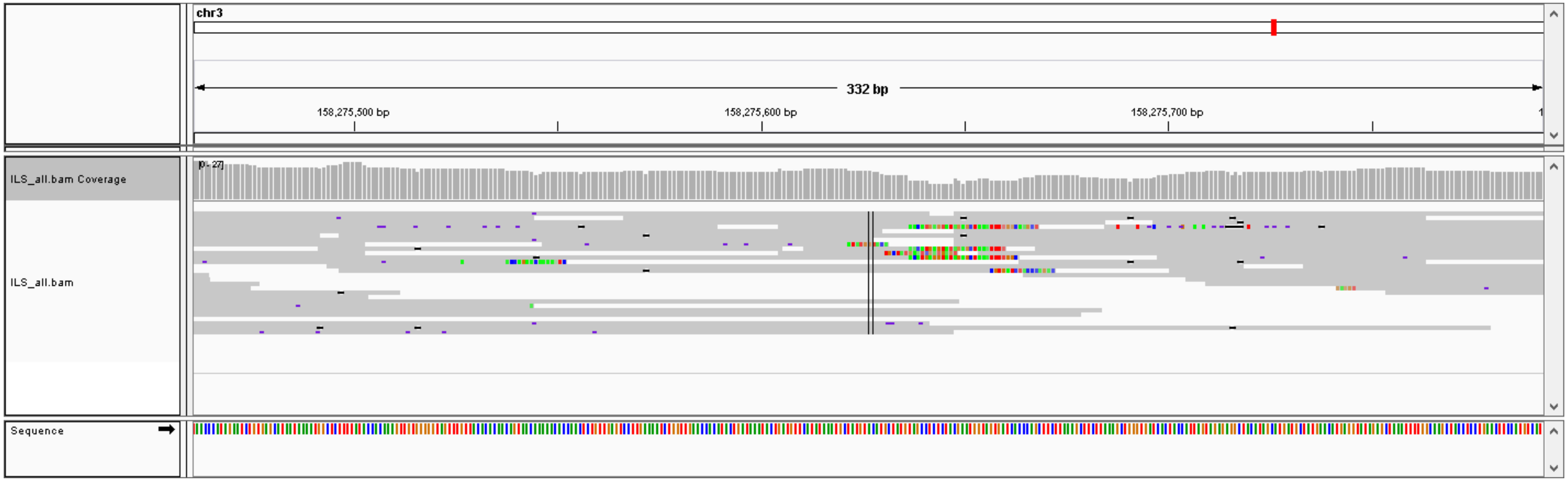

### #24, chr6: 57738609-57738722,  DEL (112 bp), validation: TP

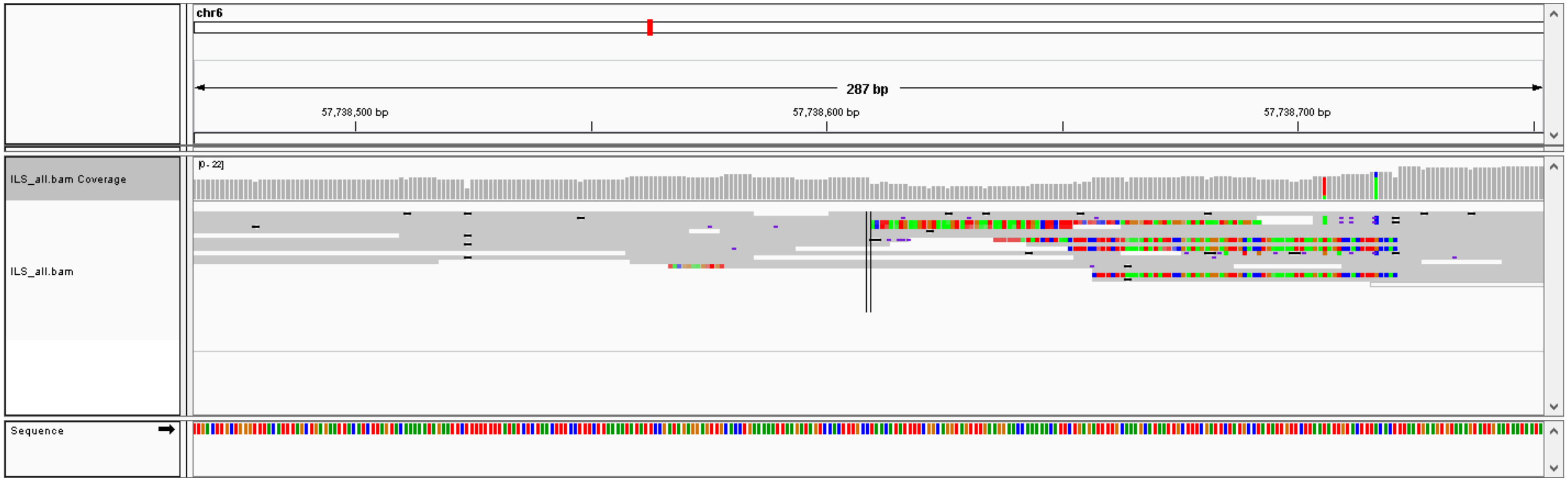

### #25, chr5: 173035295-173036241,  DEL (945 bp), validation: TP

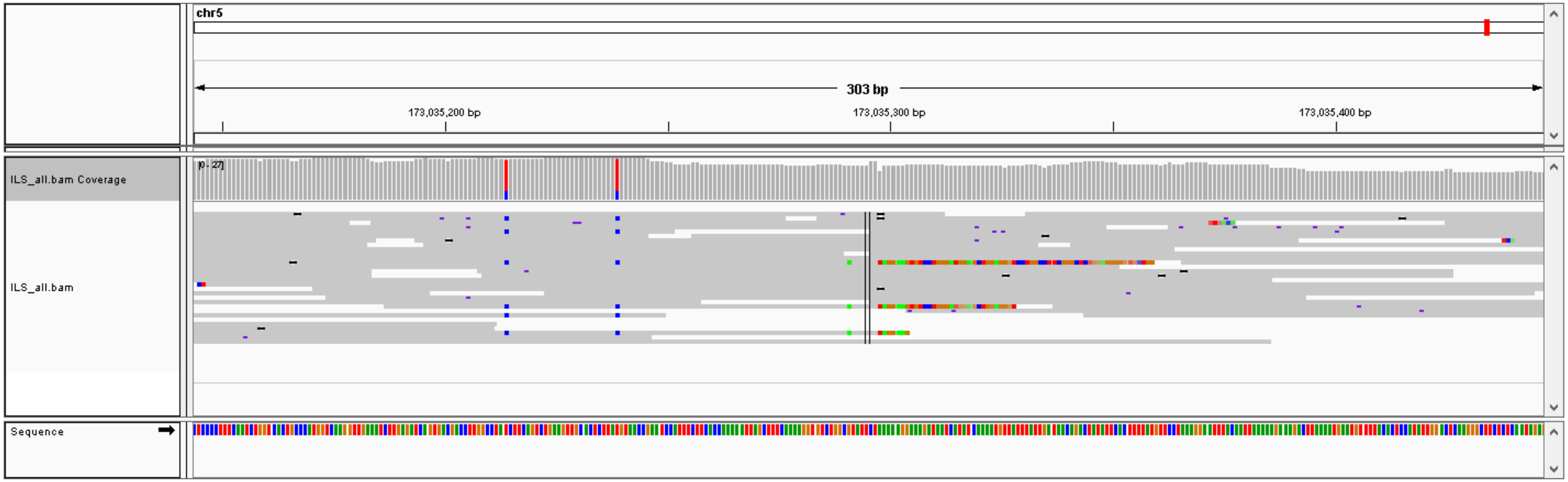


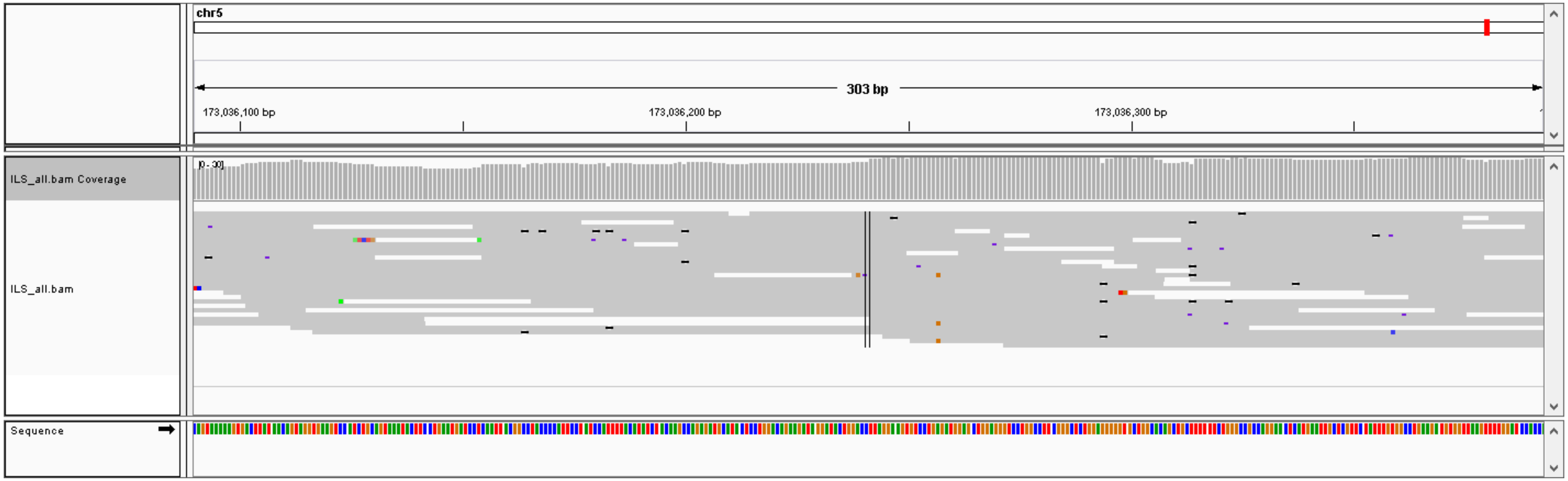

### #26, chr5: 176387579-176390181,  DEL (2601 bp), validation: TP

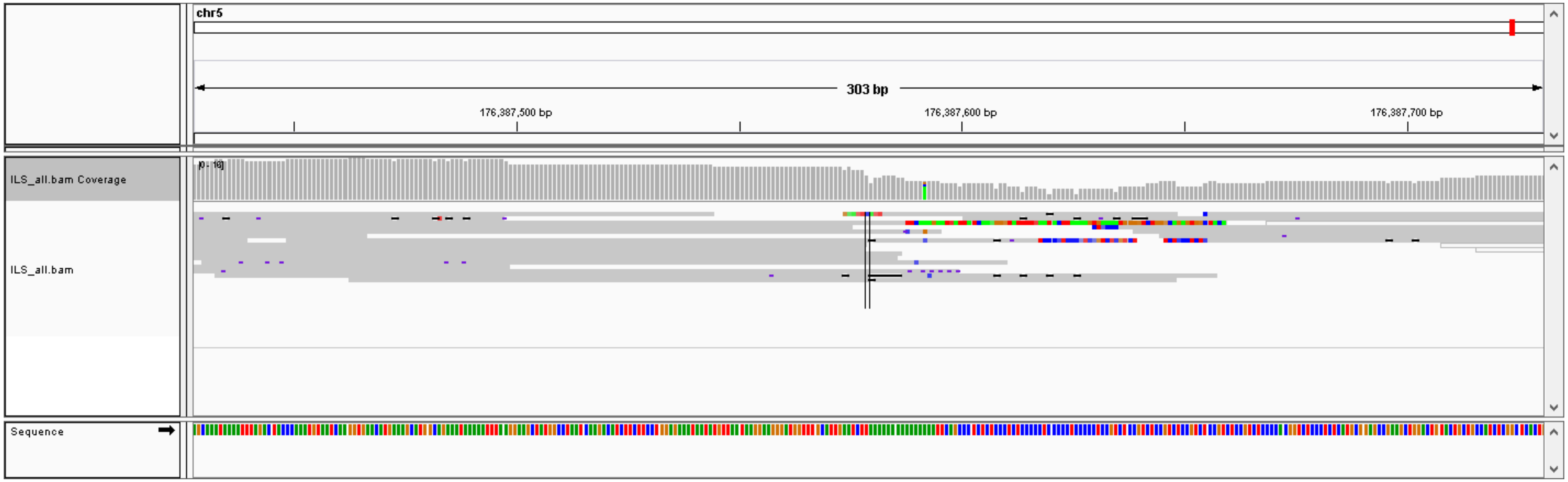


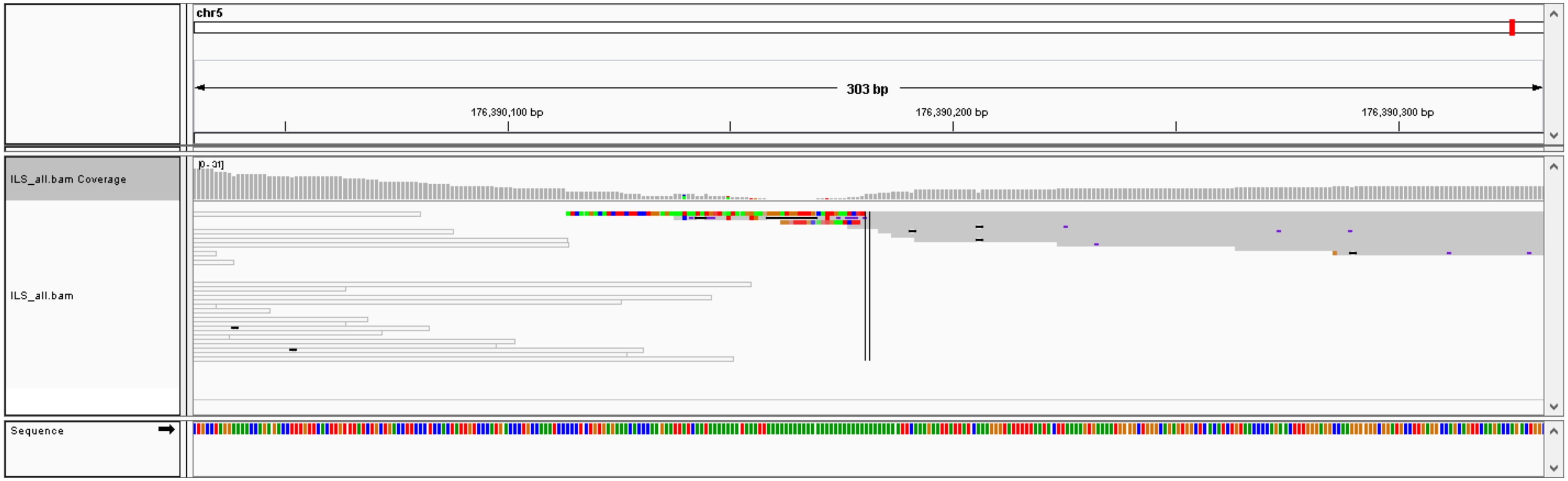

### #27, chr21: 11096782-11108481,  DEL (11698 bp), validation: FP

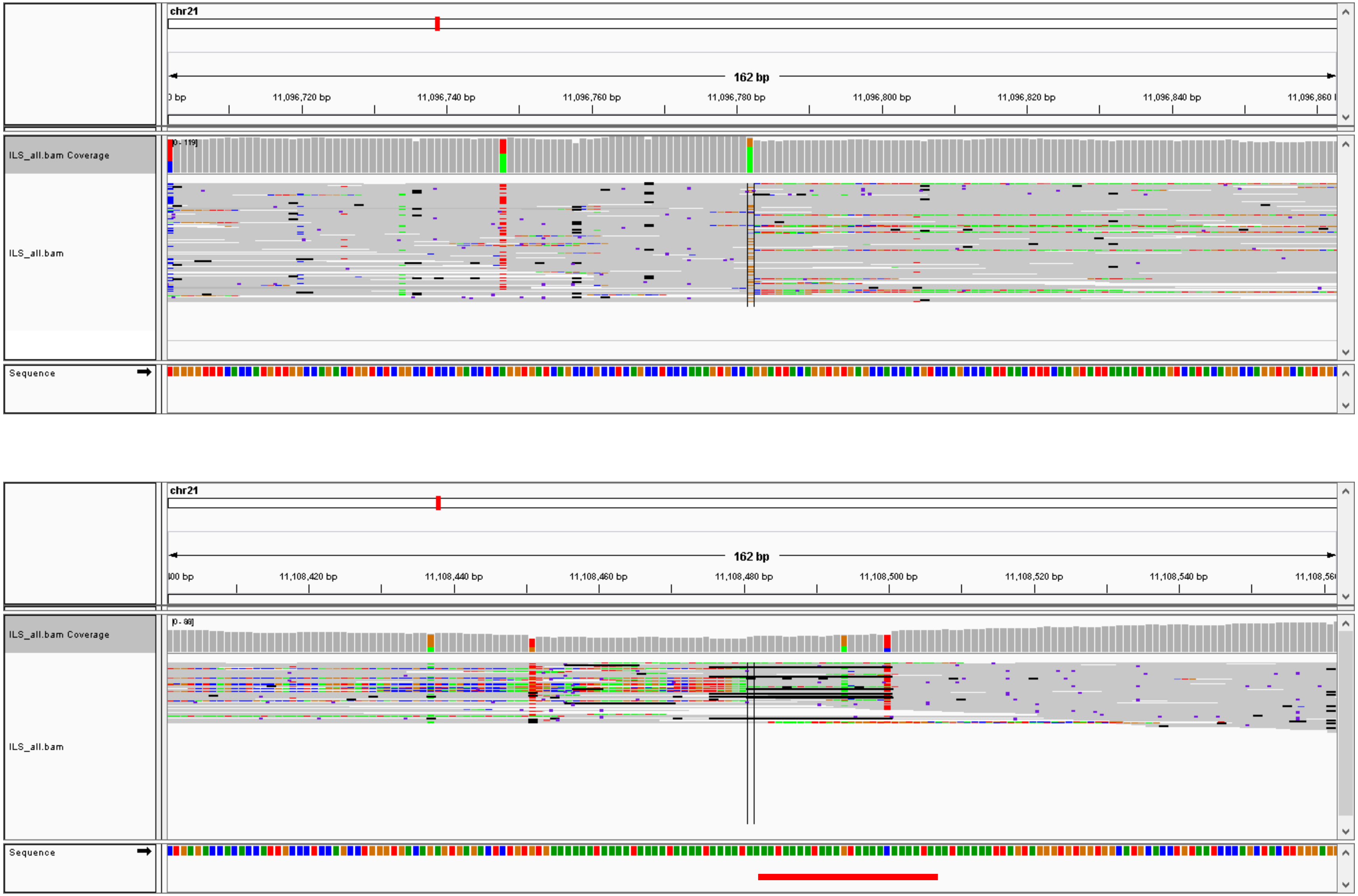


Incorrect mapping region

#28, chr2: 91679897-91685489,  DEL (5591 bp), validation: TP

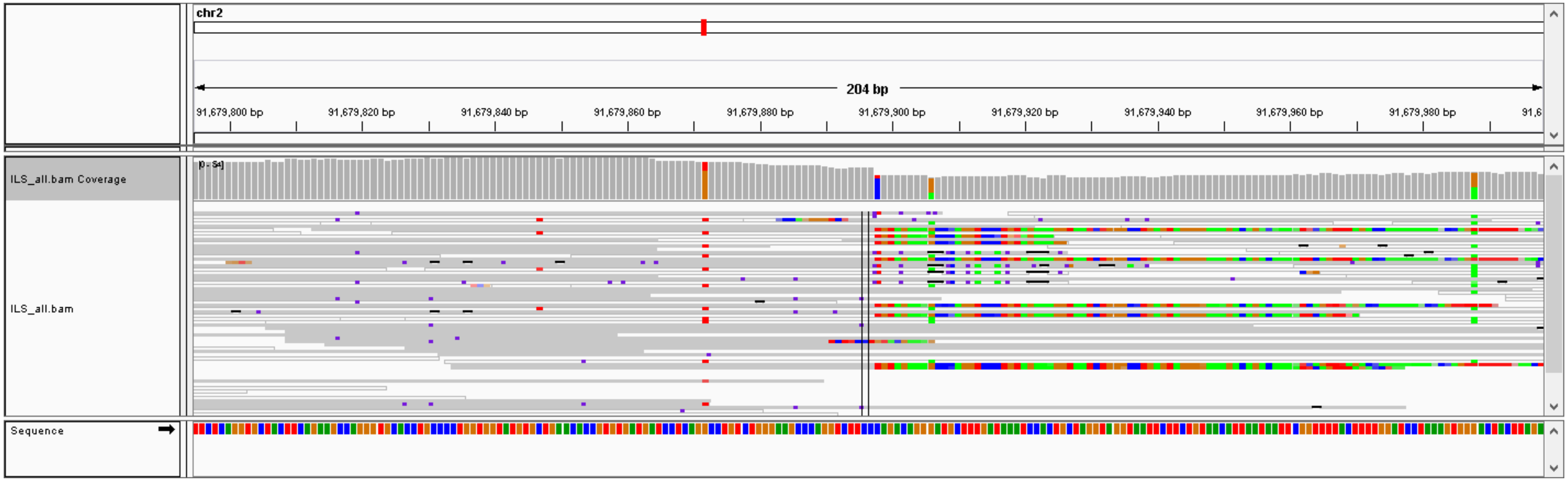


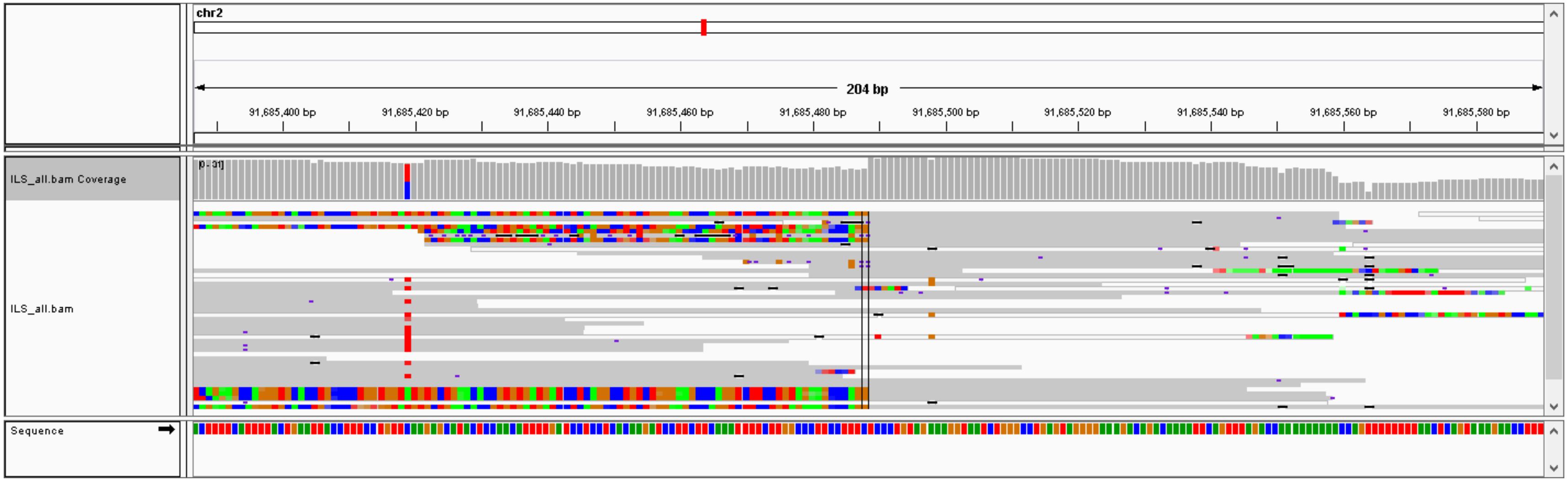

### #29, chr1: 144954613-144955362,  DEL (748 bp), validation: TP

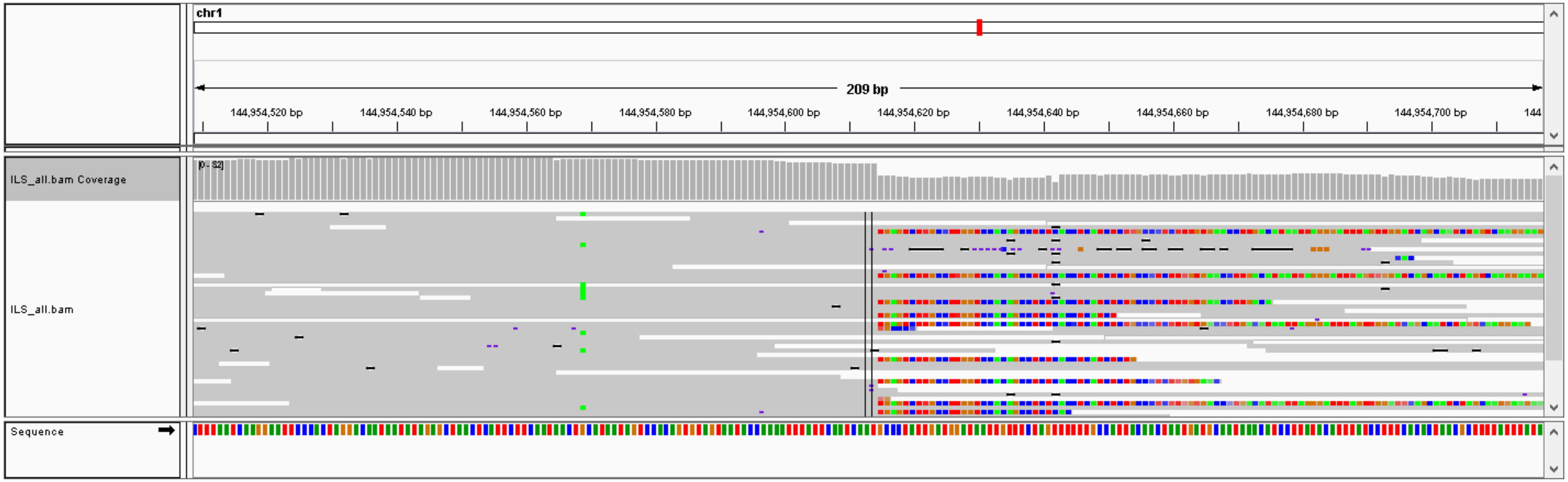


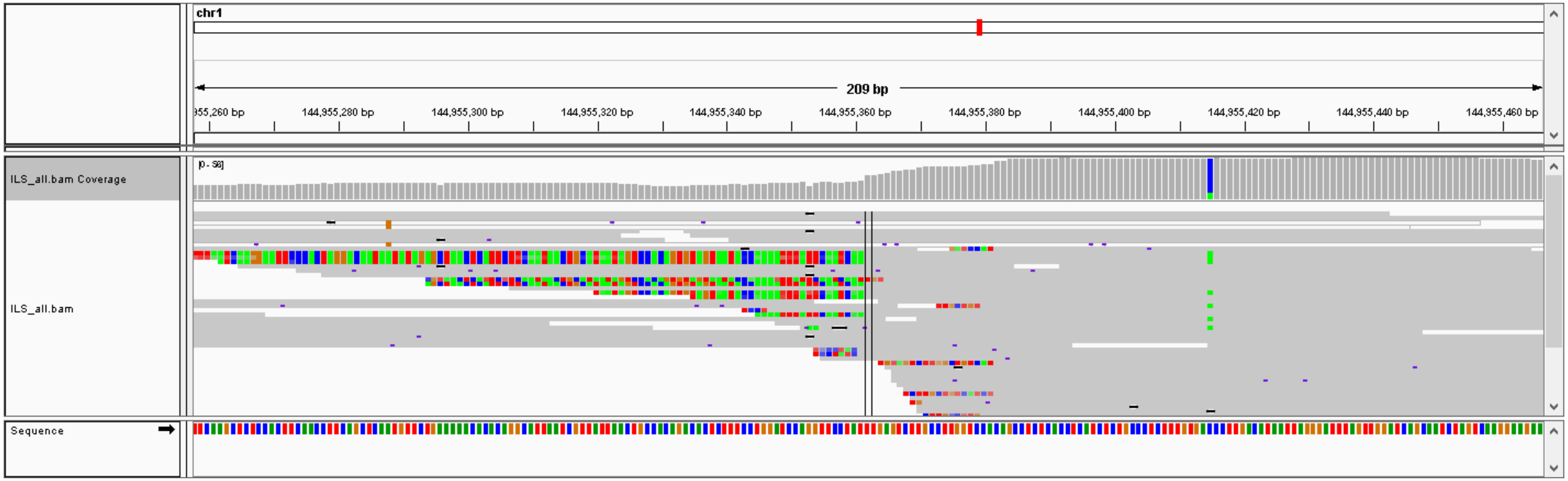

### #30, chrX: 138586559-138586630,  DEL (70 bp), validation: TP

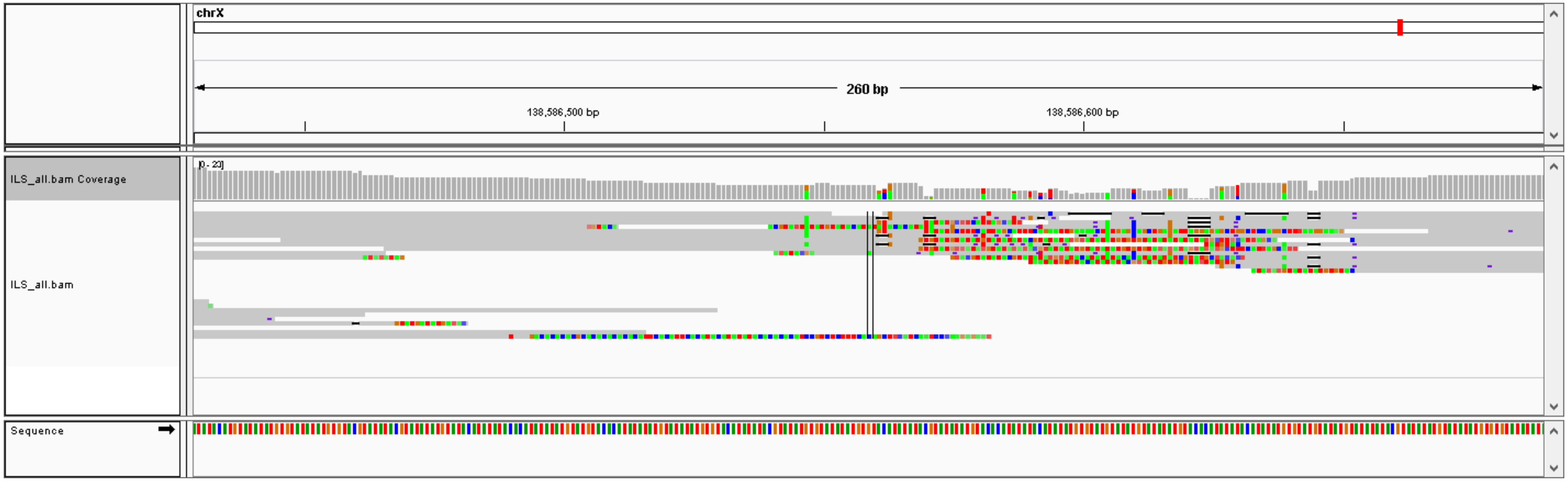

### #31, chr6: 149081506- chr22: 33414615,  CTX, validation: TP

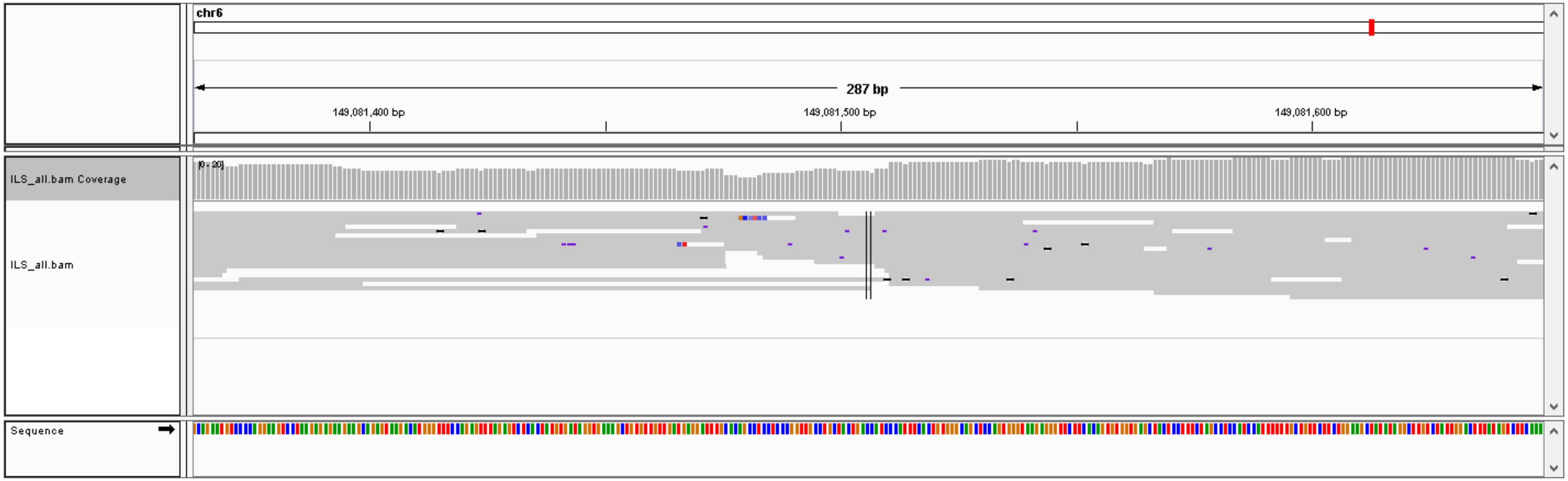


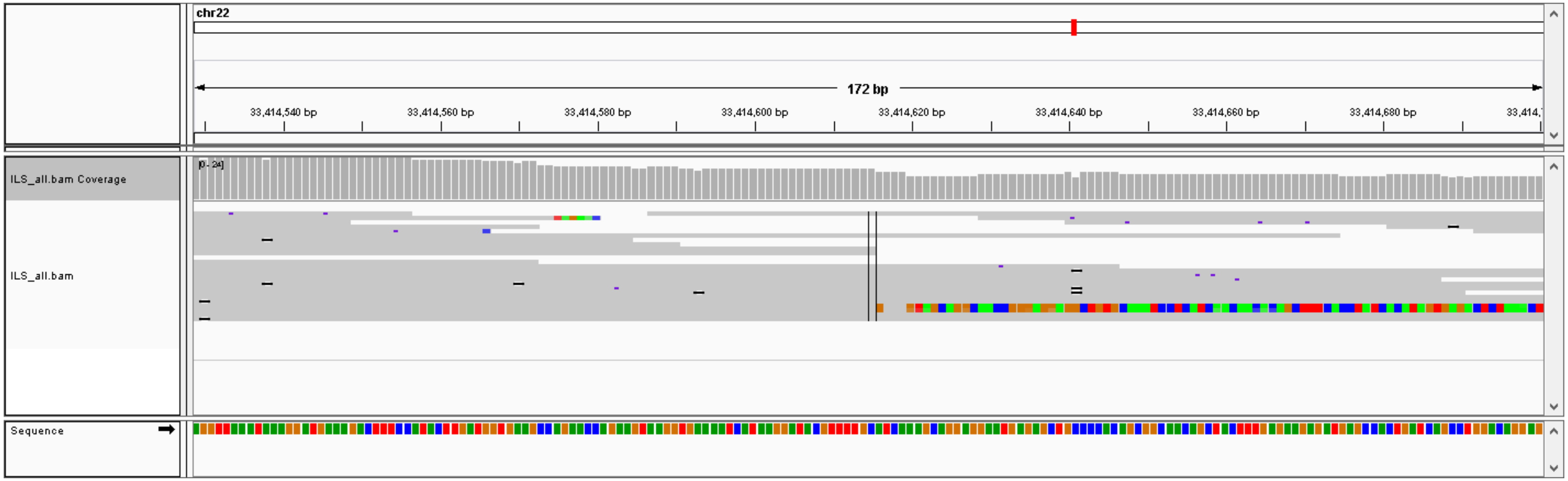

### #32, chr11: 28946243- chr19: 40865236,  CTX, validation: TP

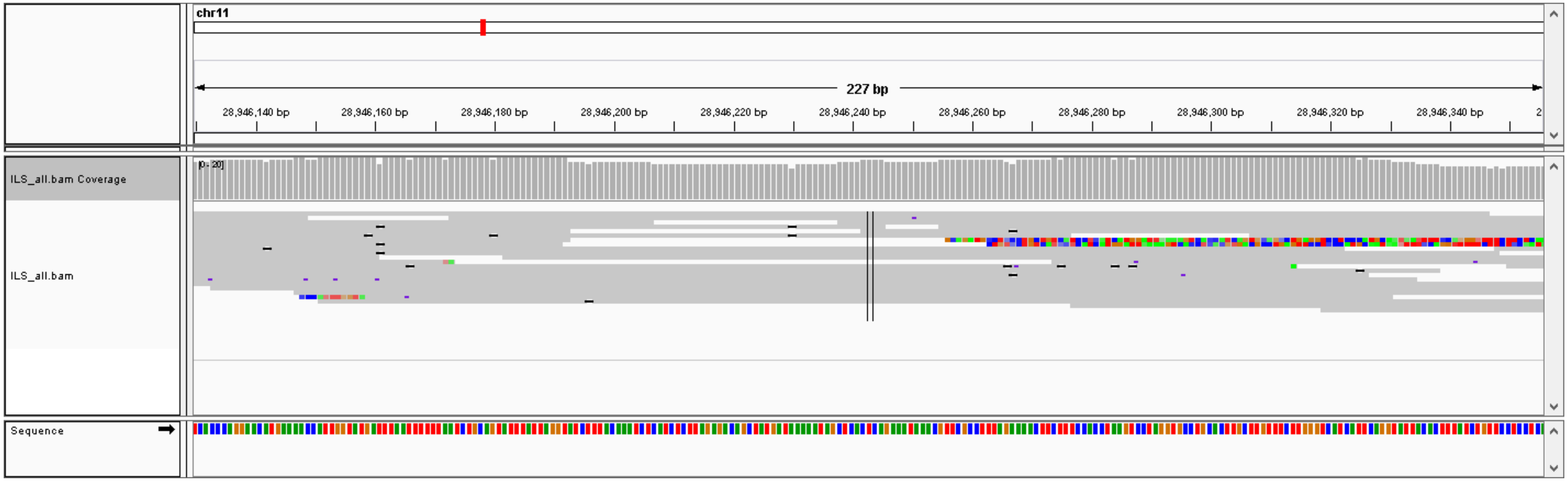


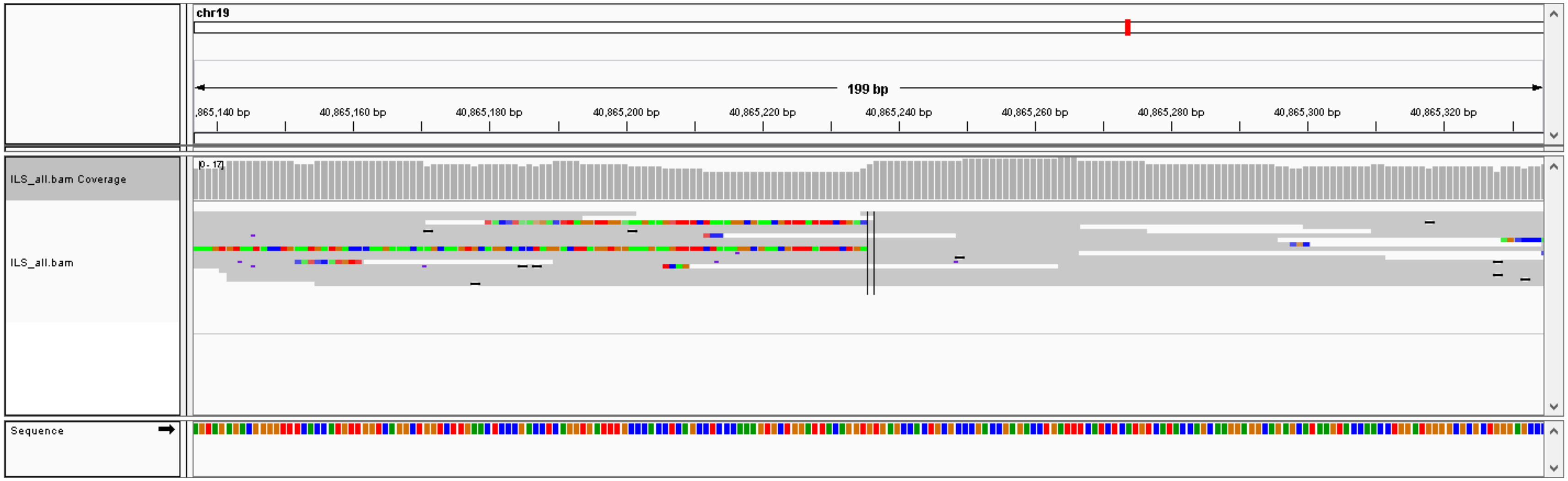

### #33, chr4: 31034908- chr5: 29574787,  CTX, validation: TP

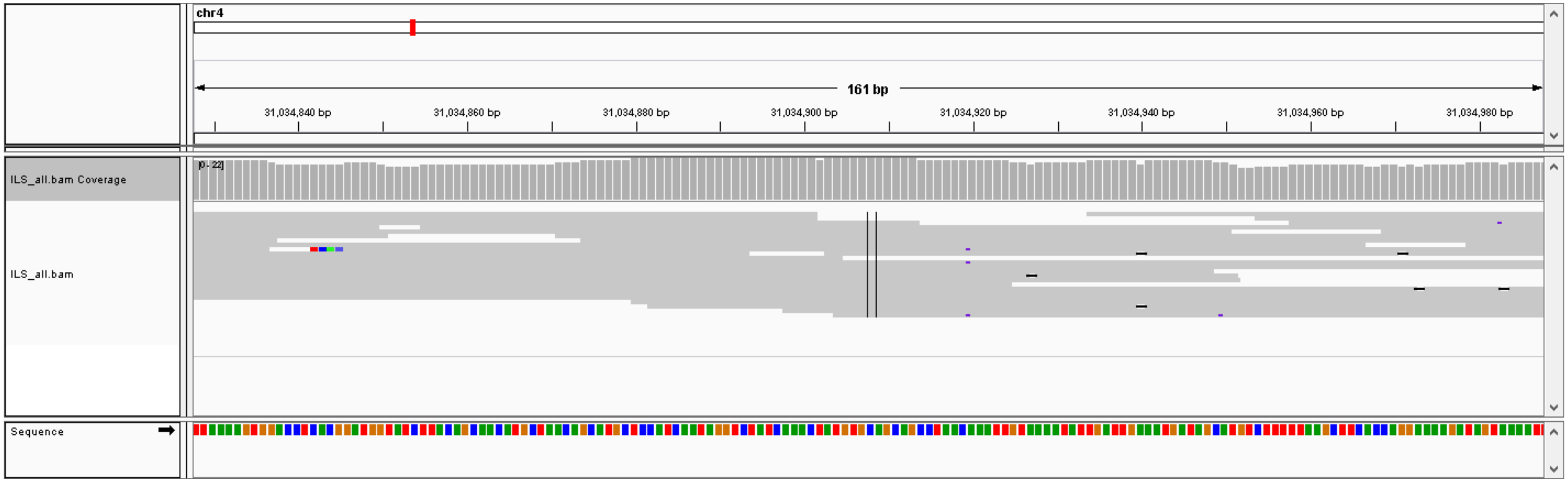


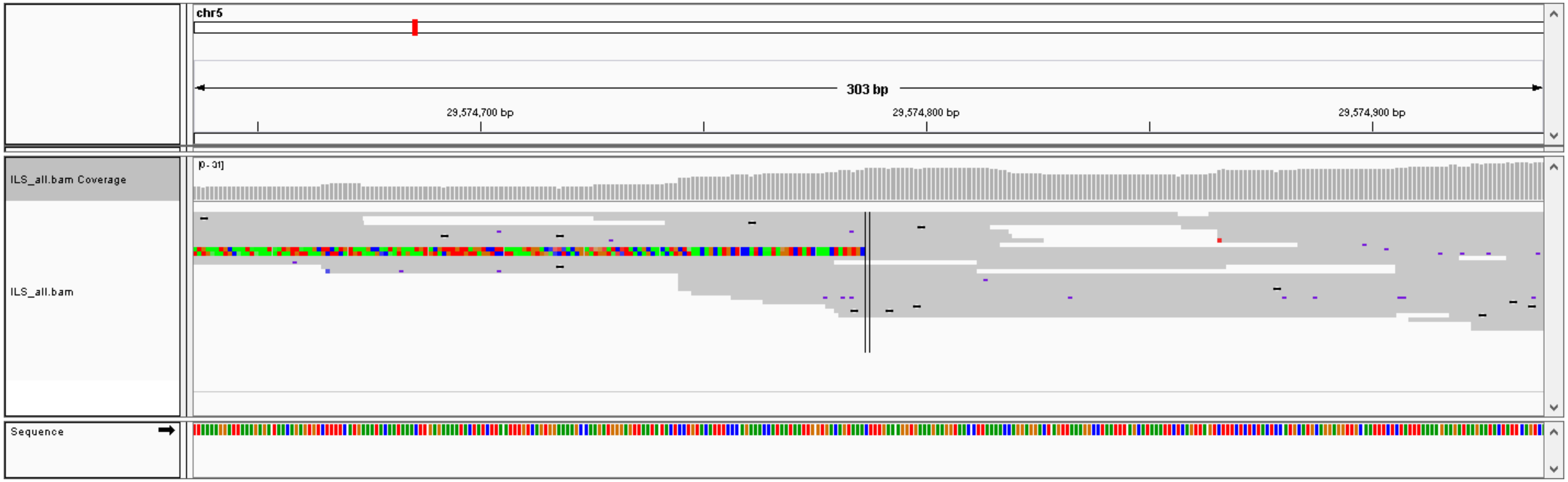

### #34, chr2: 228915510- chrX: 40536242,  CTX, validation: TP

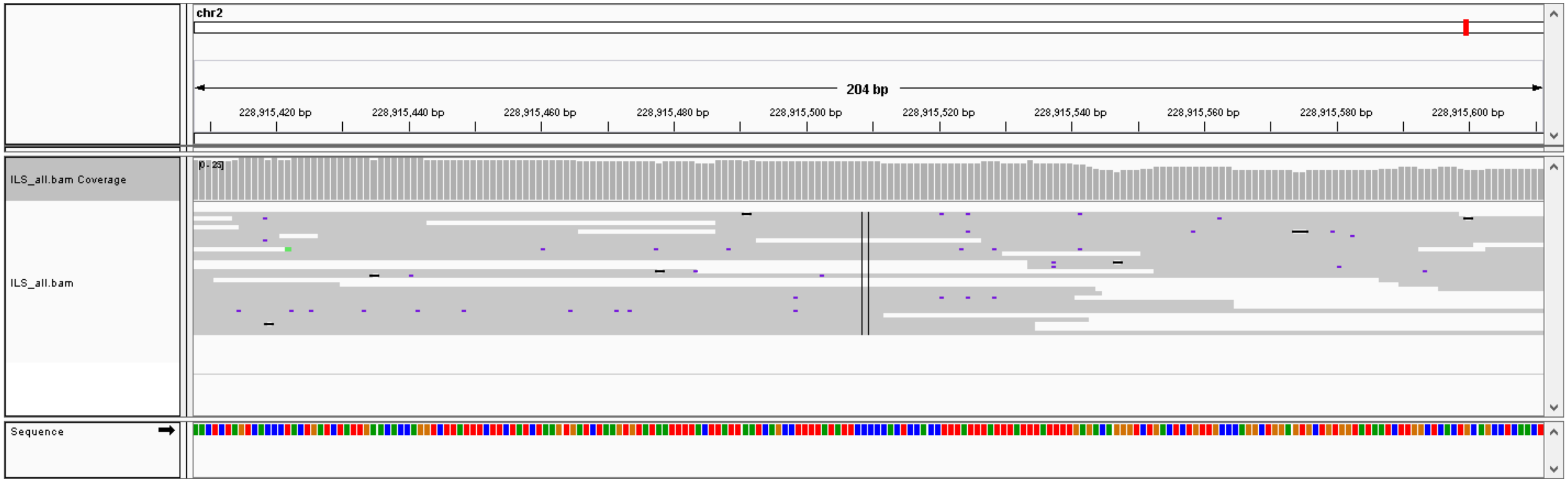


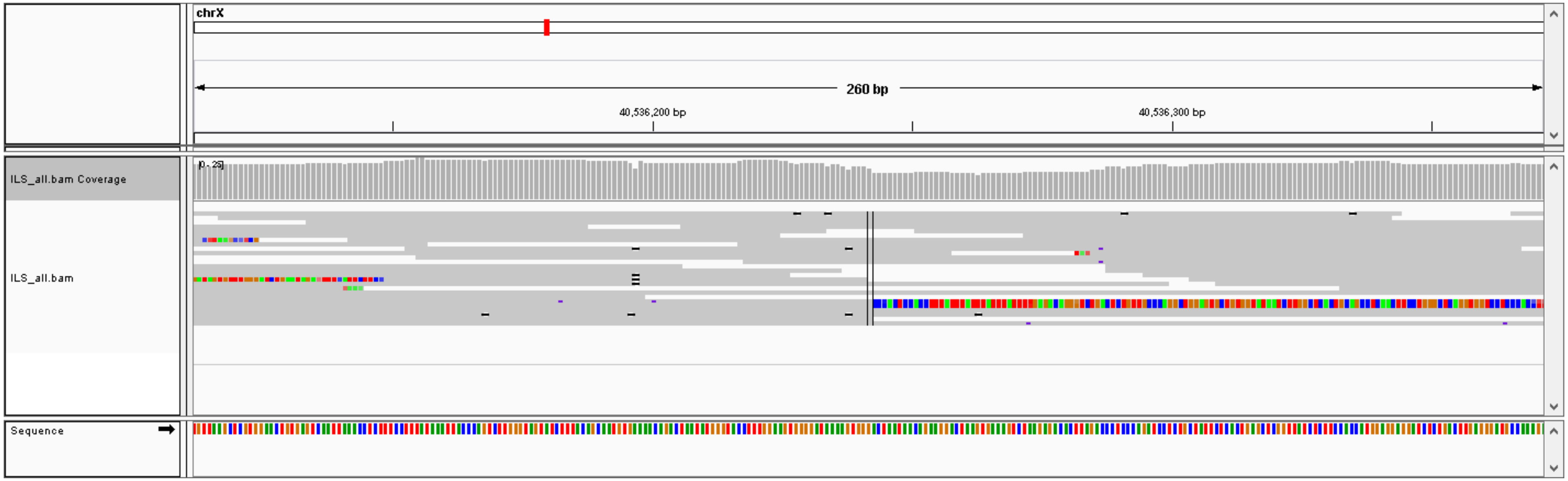

### #35, chr8: 18617473- chr1: 236594936,  CTX, validation: FP

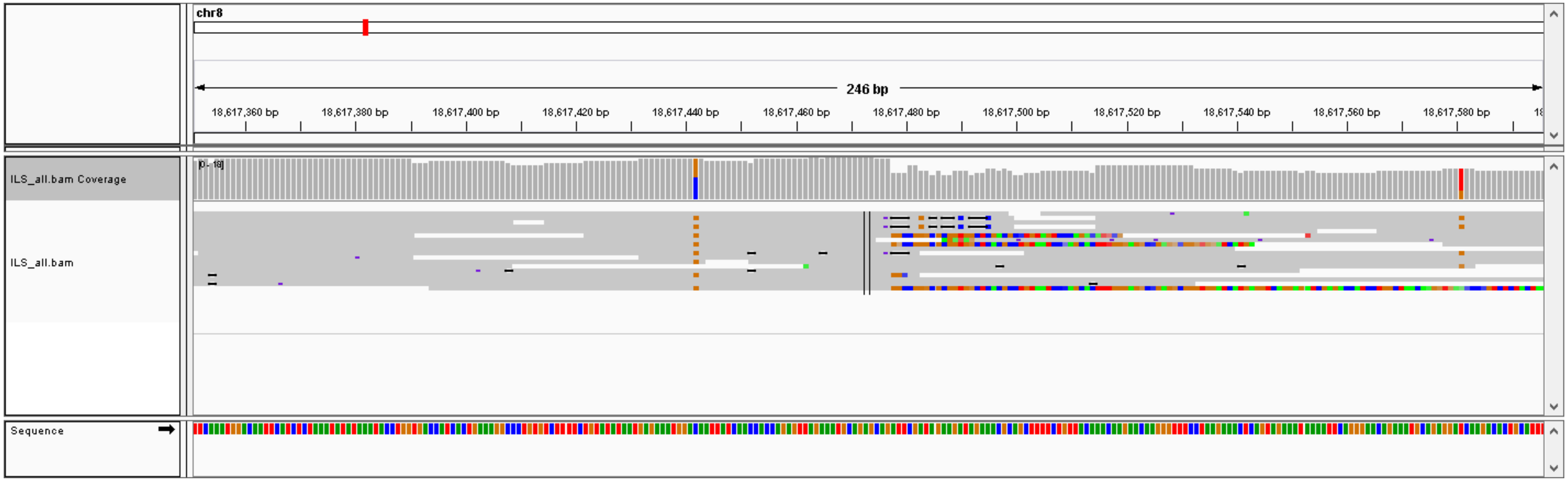


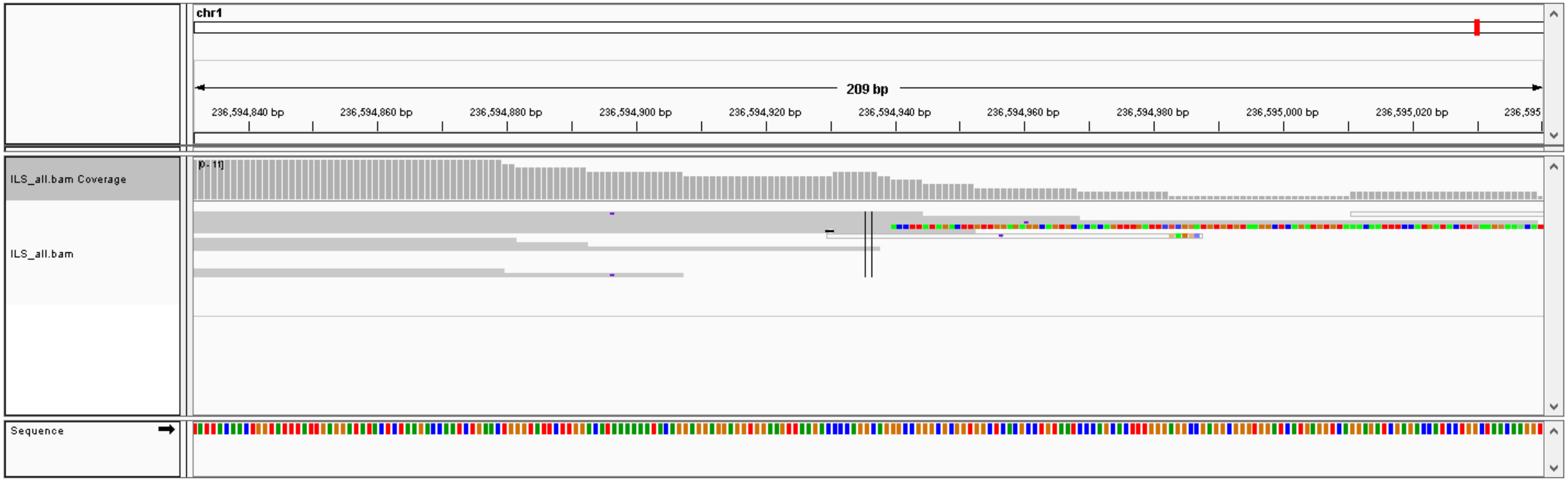


Very low coverage

### #36, chr1: 121484728- chr11: 29533224,  CTX, validation: FP

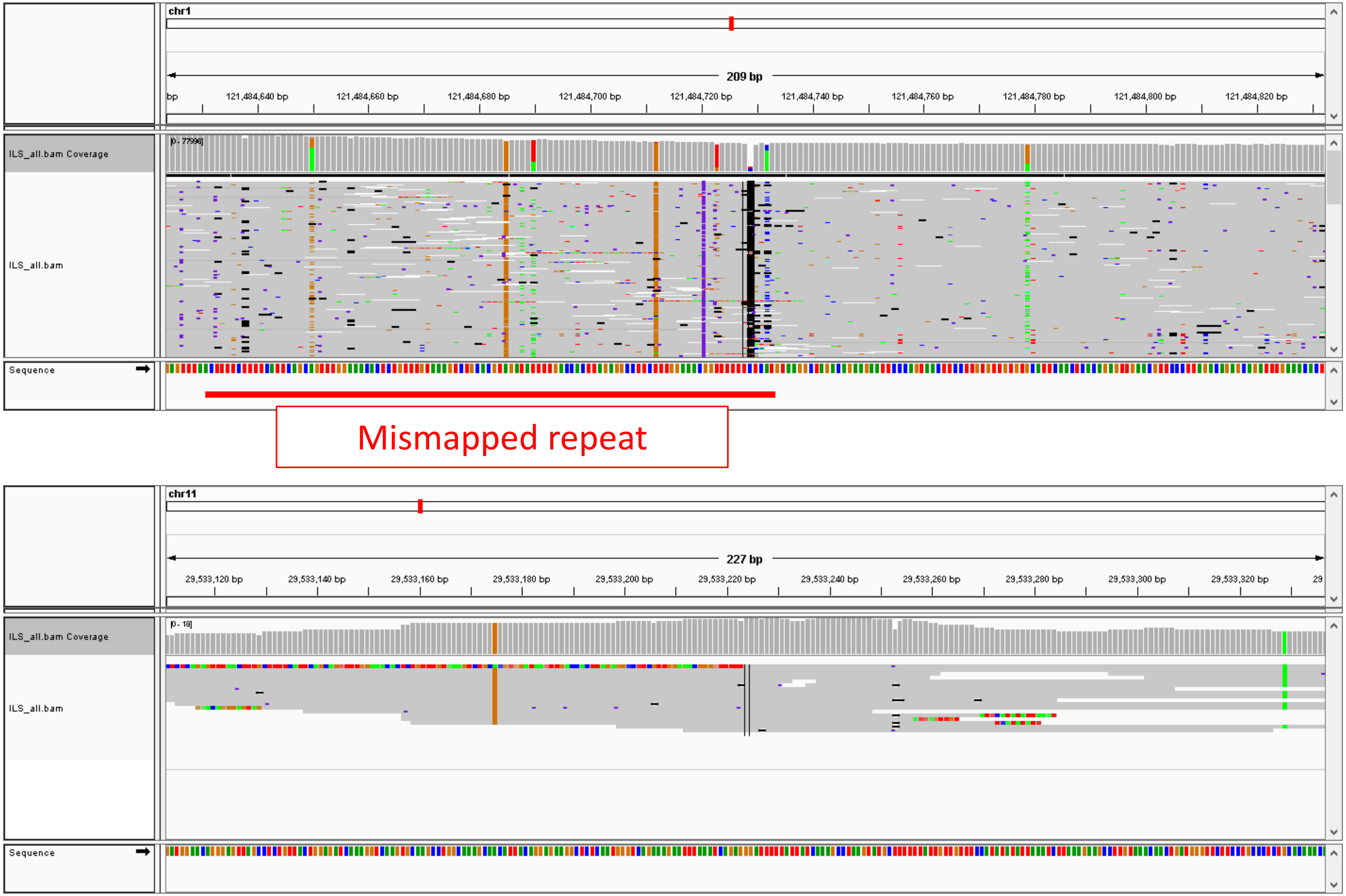

#37, chr10: 42597040- chr11: 18775637, CTX, validation: FP

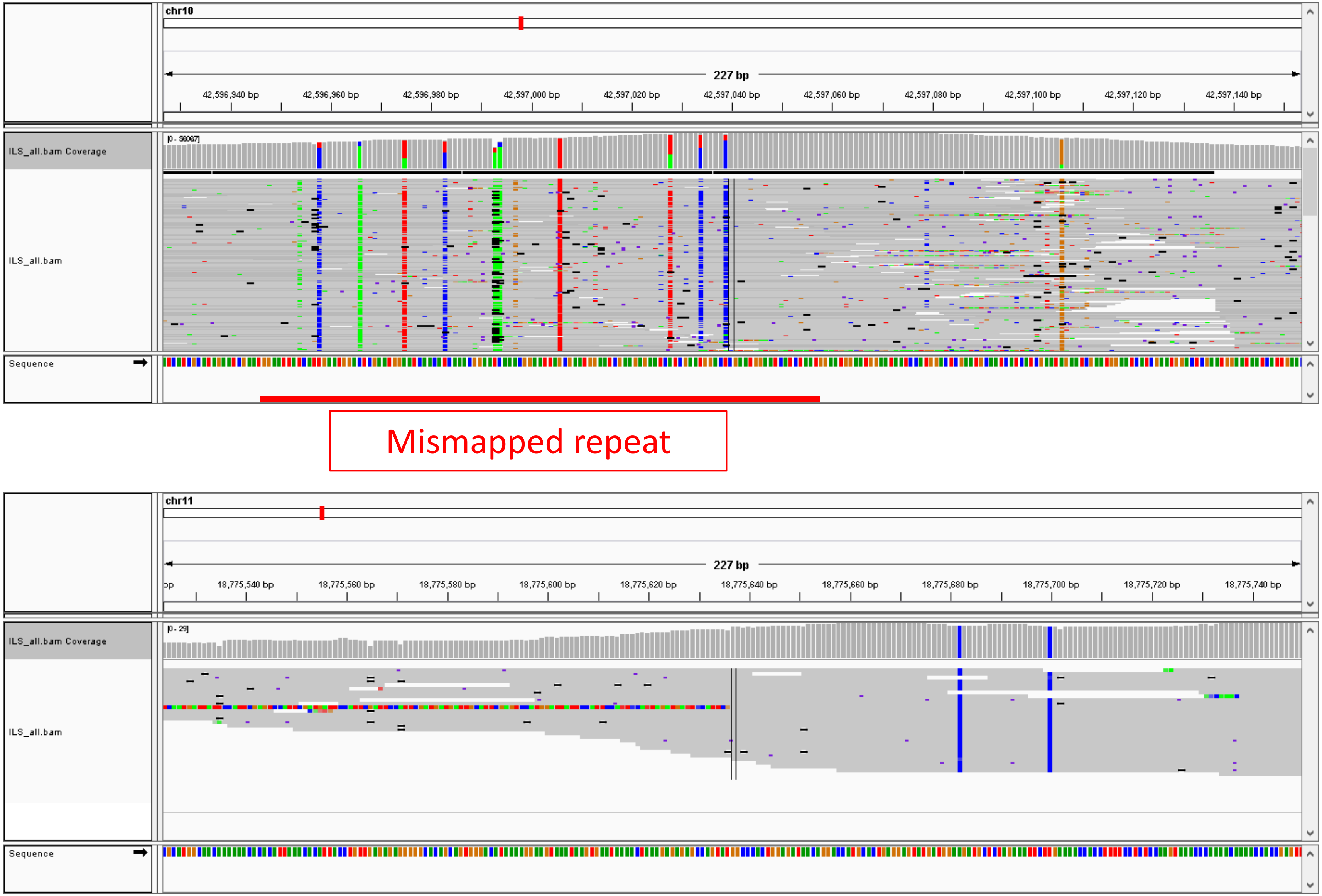

### #38, chr11: 21264489- chr12: 77881618,  CTX, validation: TP

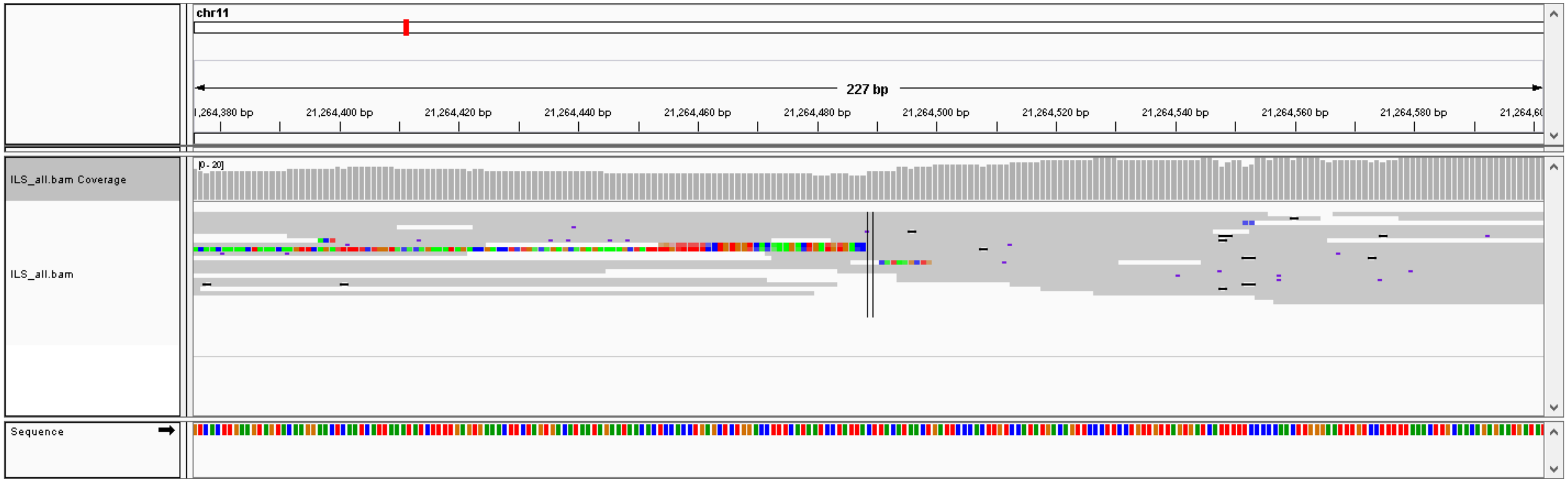


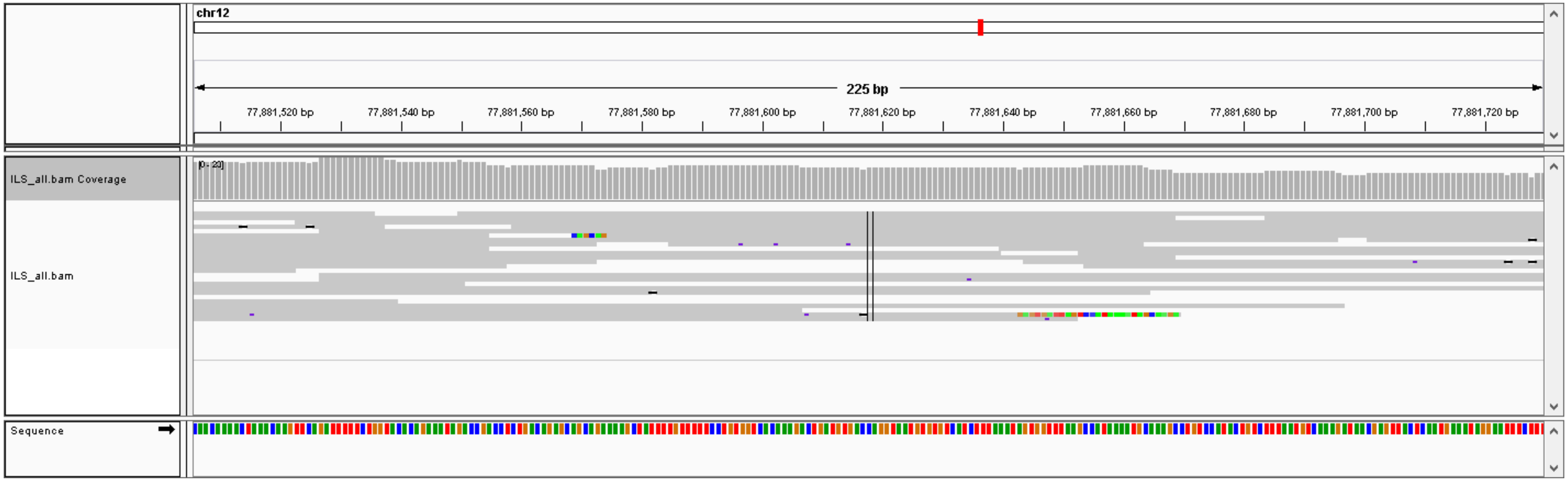

#39, chr2: 230045488- chr11: 102014349,  CTX, validation: FP

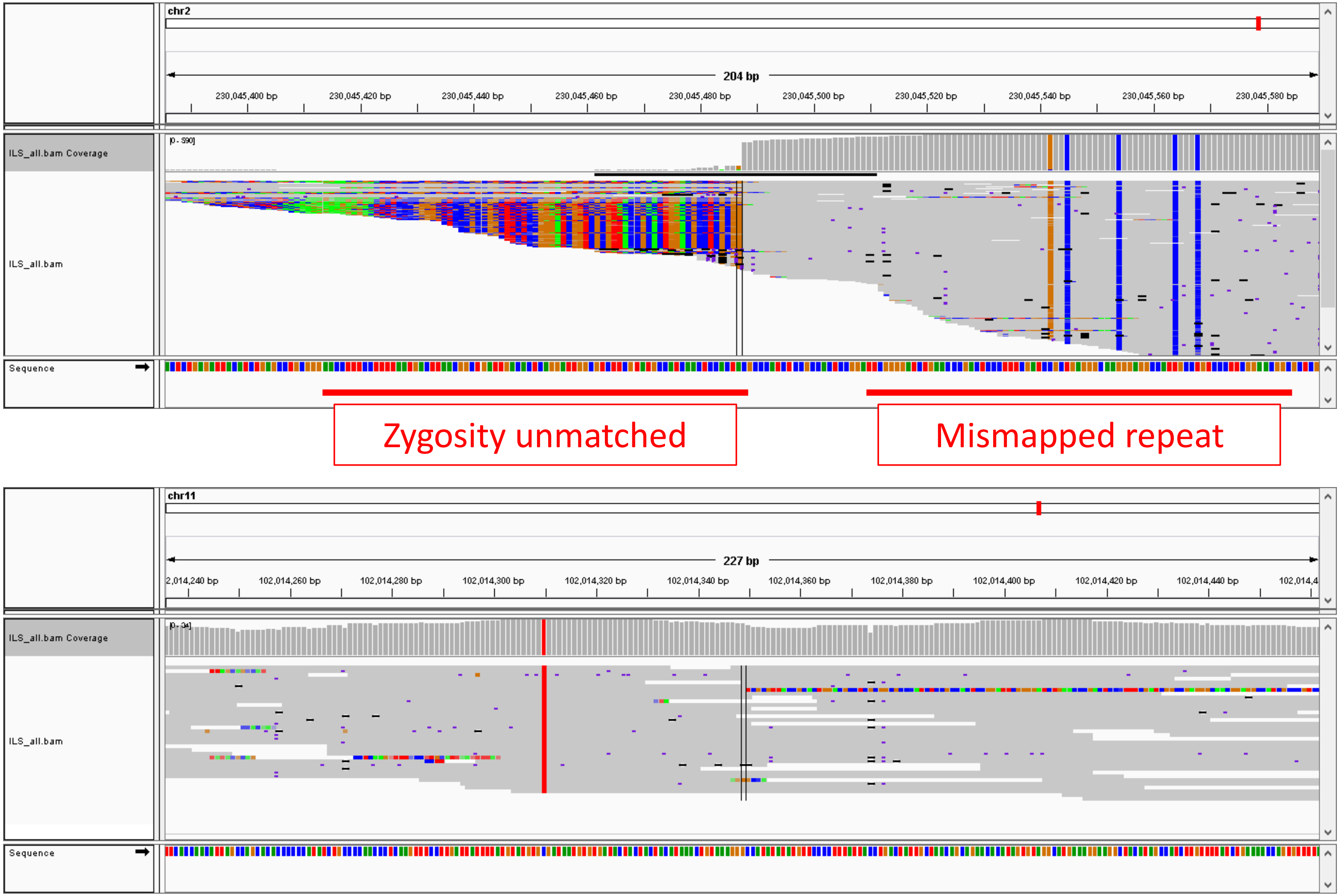

#40, chr5: 269395- chr19: 9107409,  CTX, validation: TP

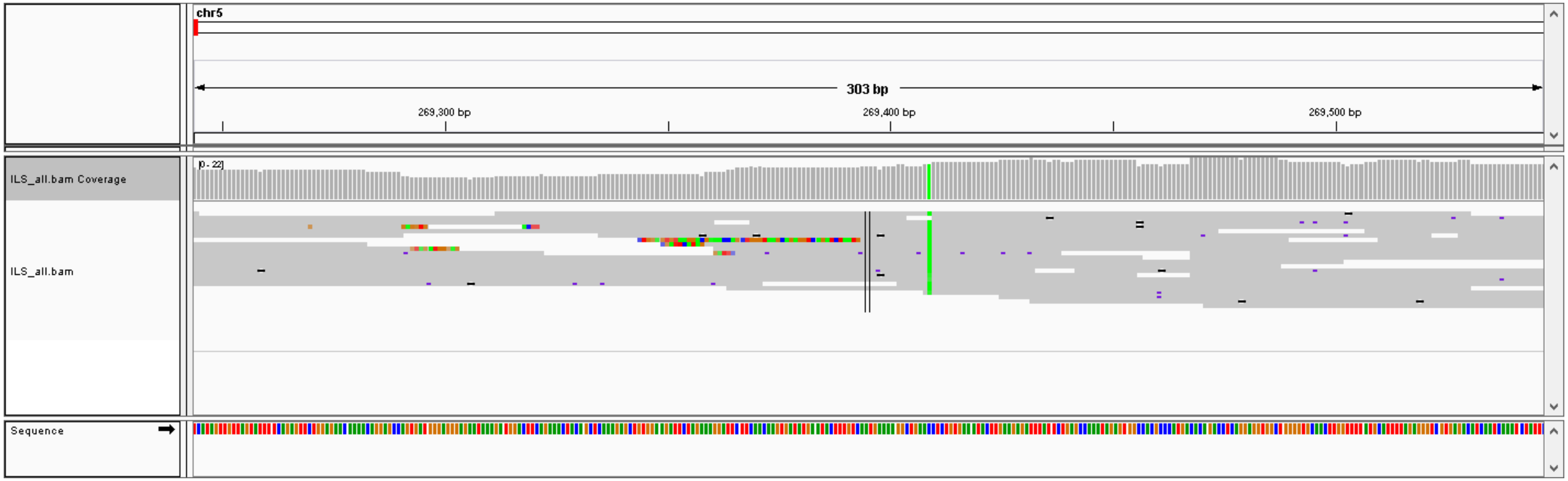


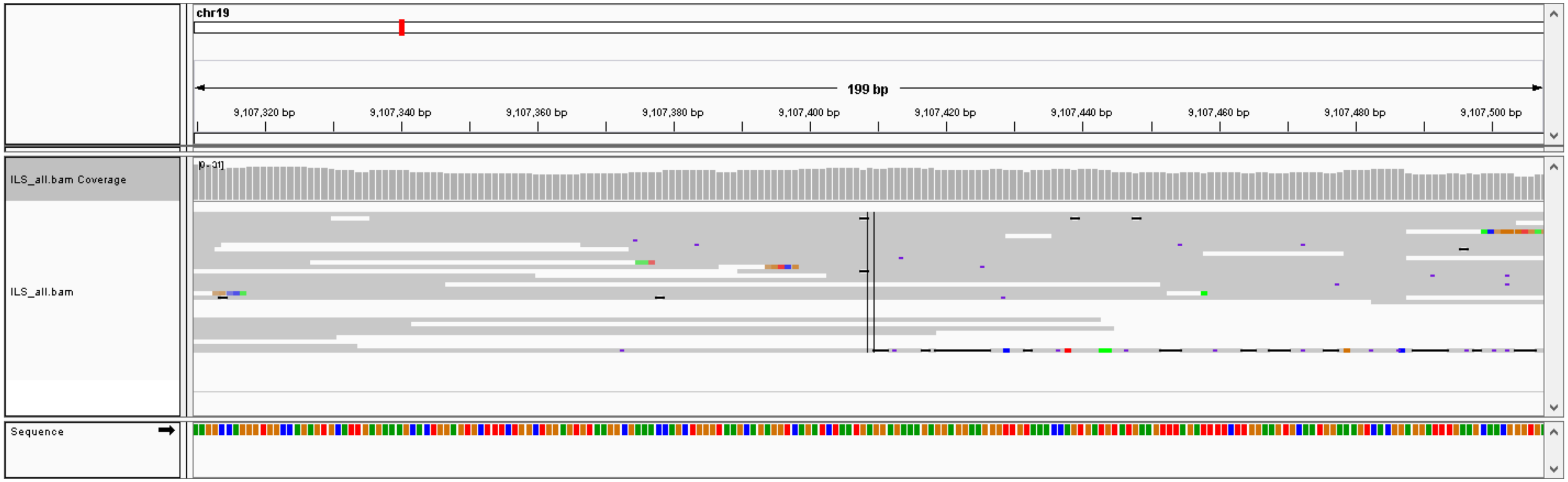

### #41, chr2: 142350998-142351324,  DEL (325 bp), validation: TP

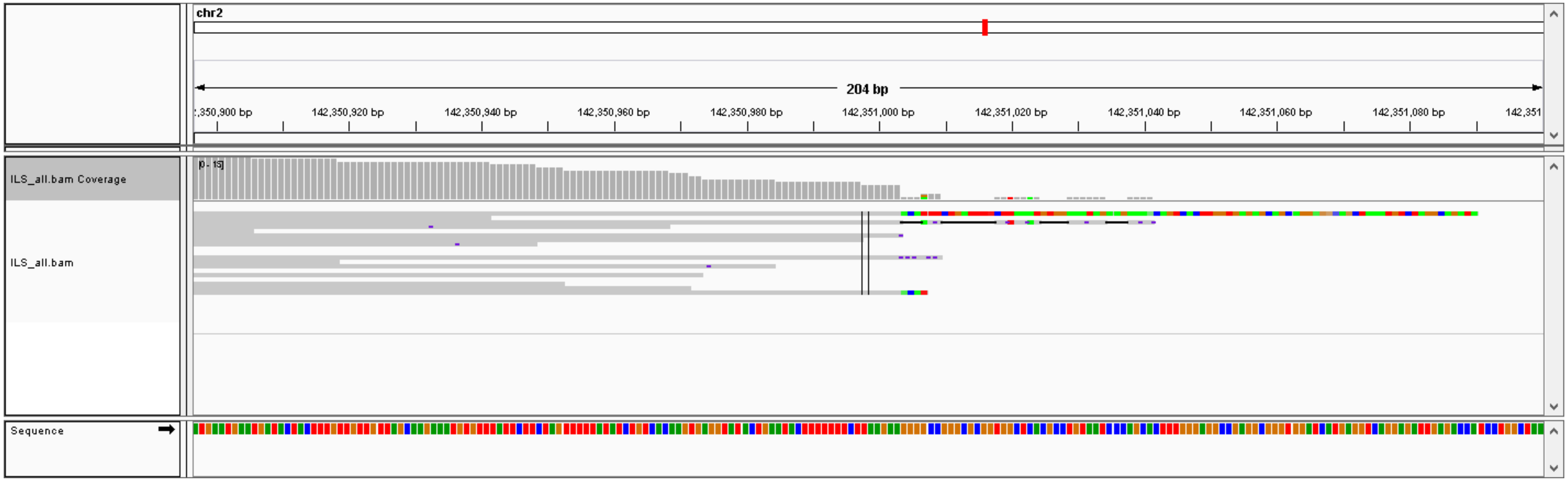


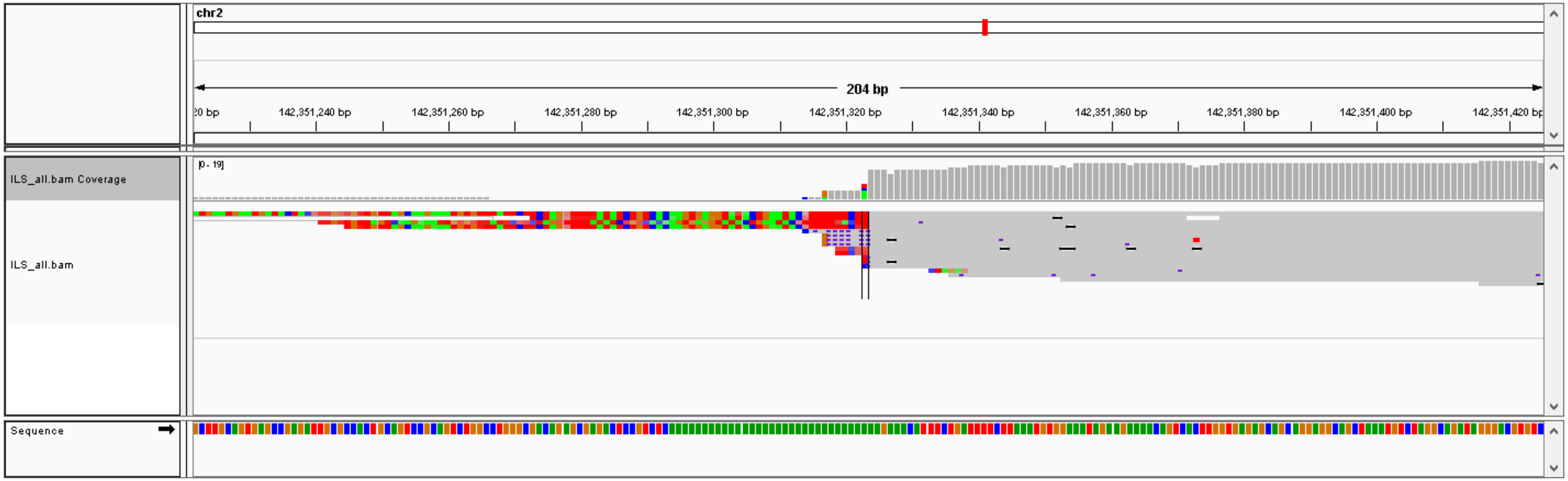

### #42, chr12: 41183195-41183502,  DEL (306 bp), validation: TP

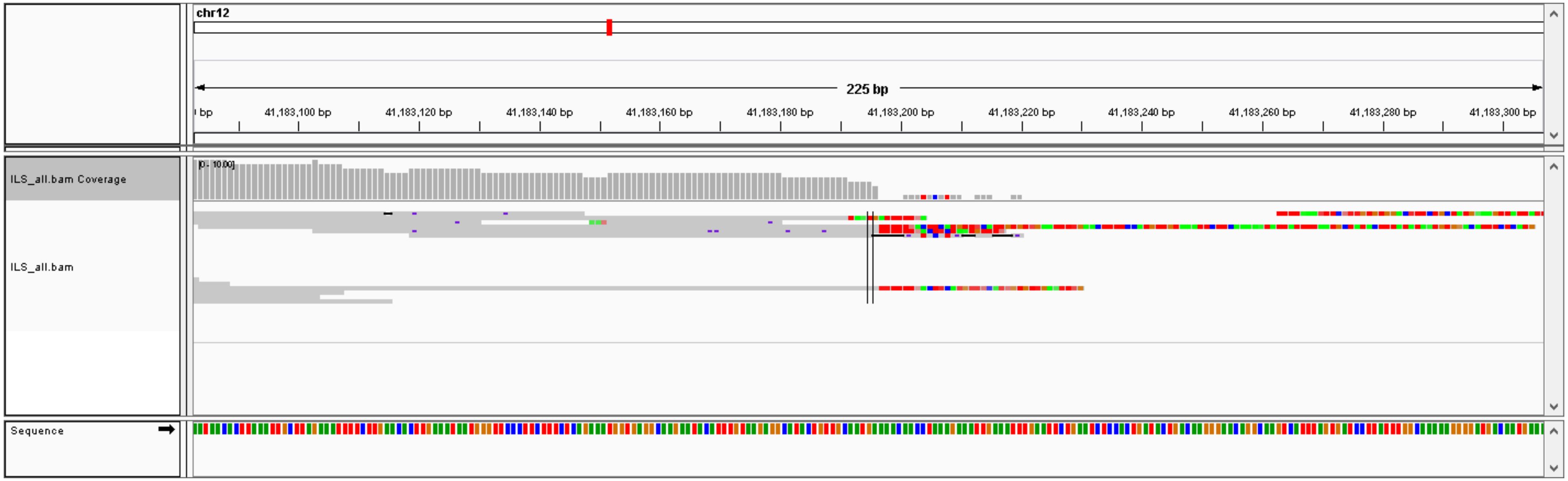


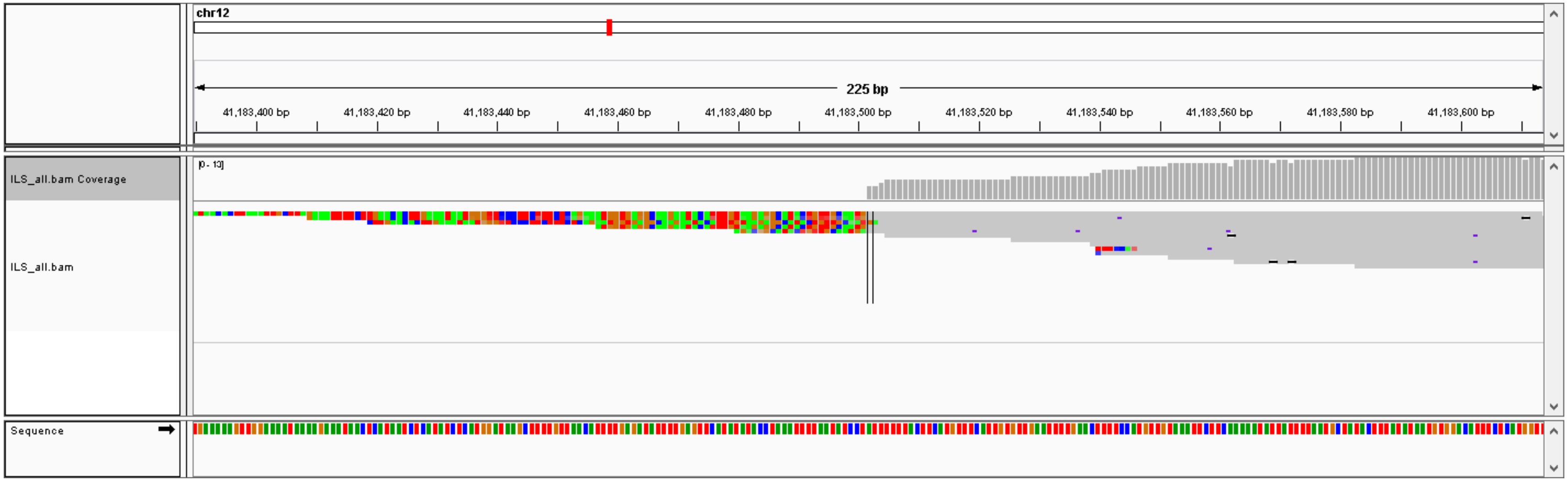

#43, chr1: 28247799-28248109, DEL (309 bp), validation: TP

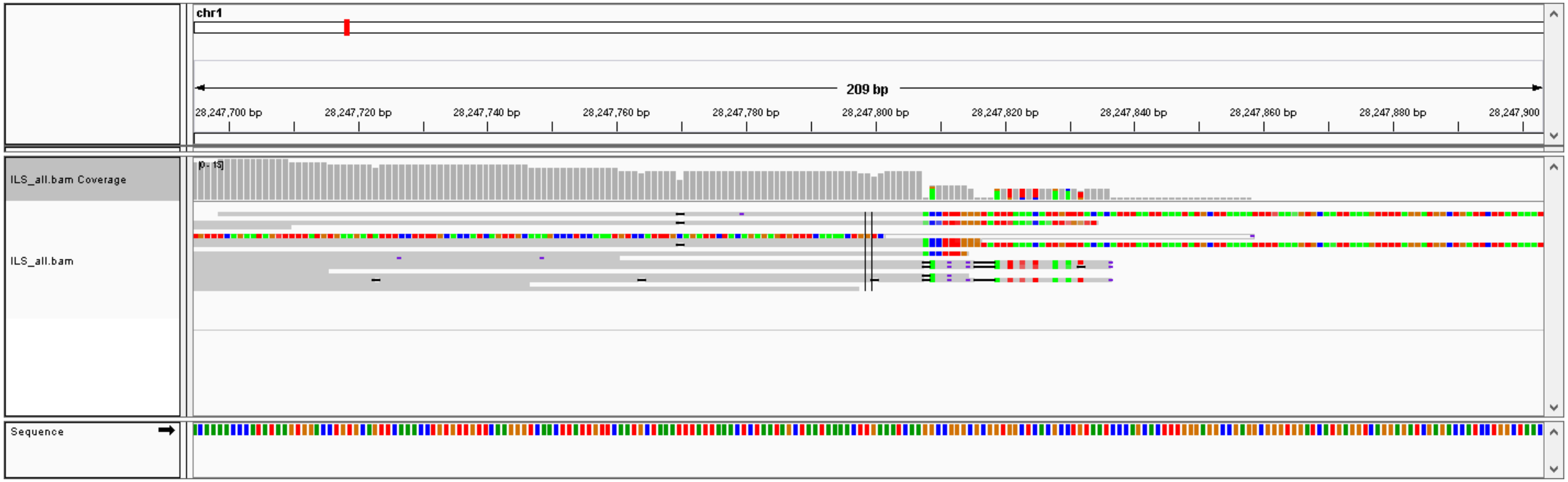


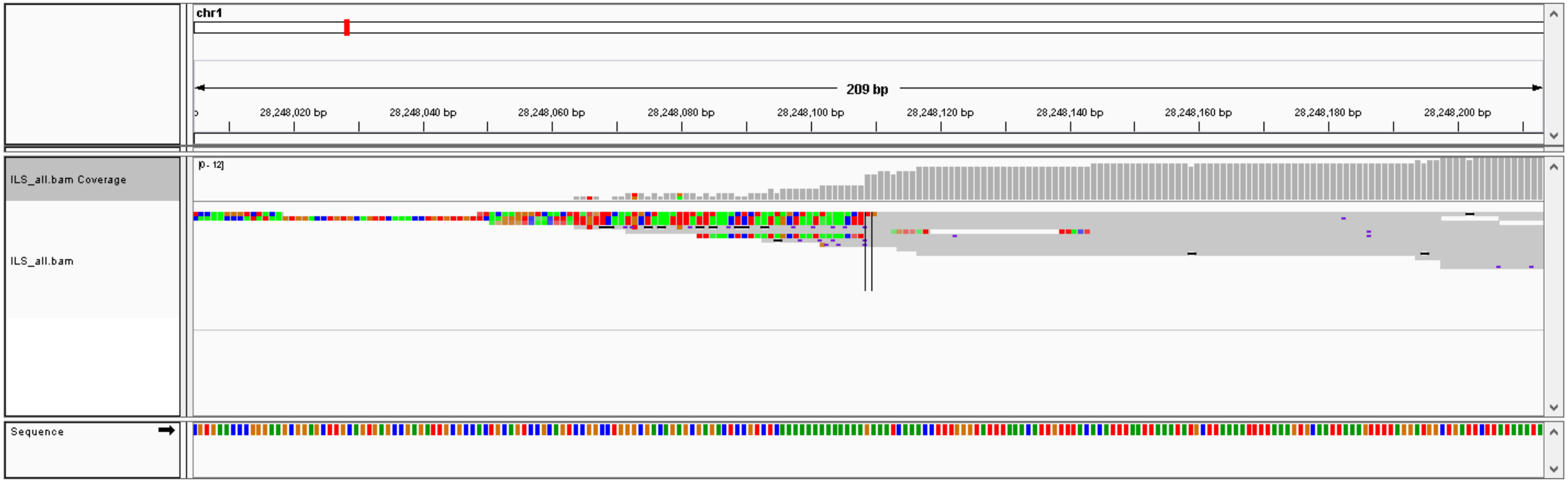

### #44, chr17: 26780387-26783494, DEL (3106 bp), validation: TP

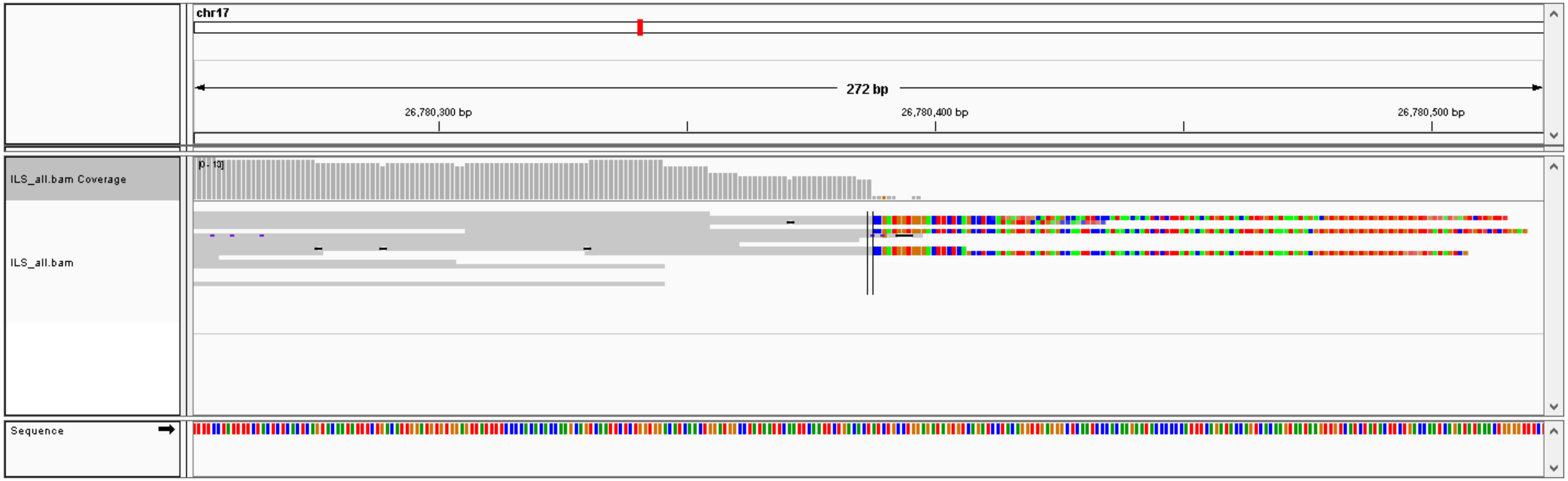


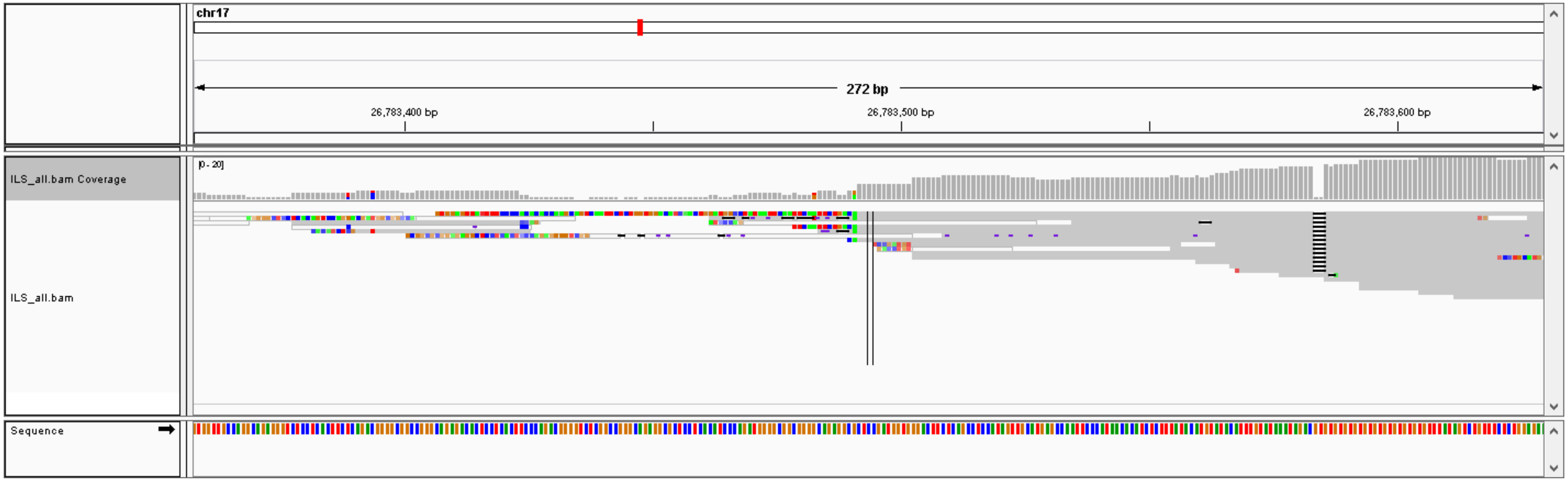

### #45, chr22: 27168066-27169793, DEL (1726 bp), validation: TP

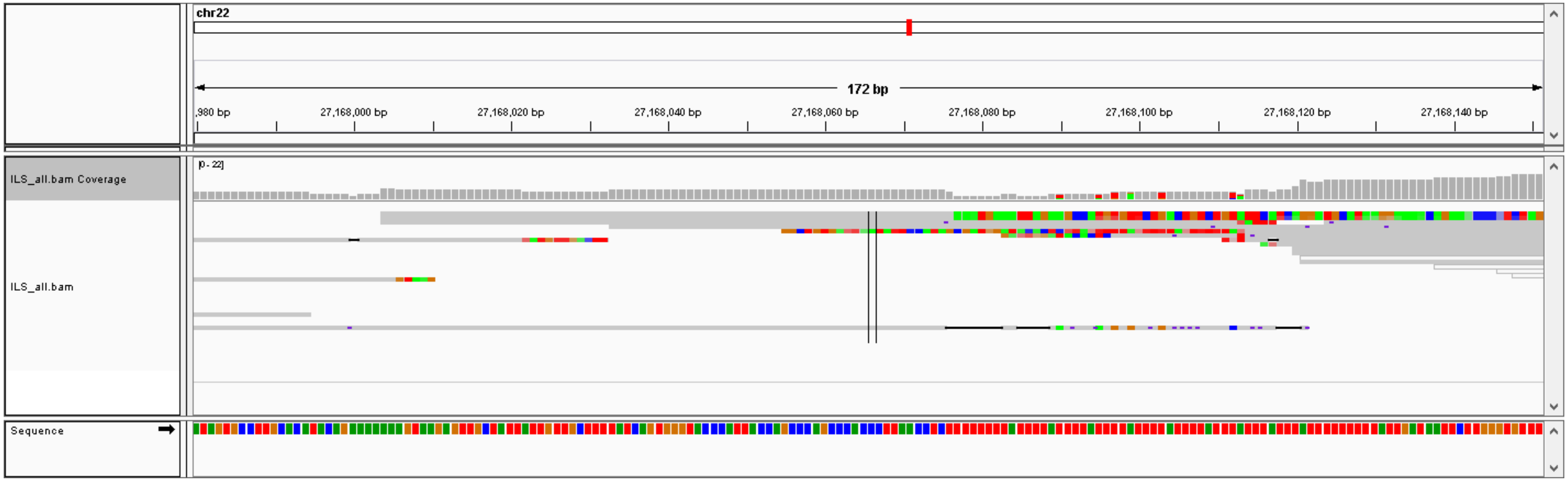


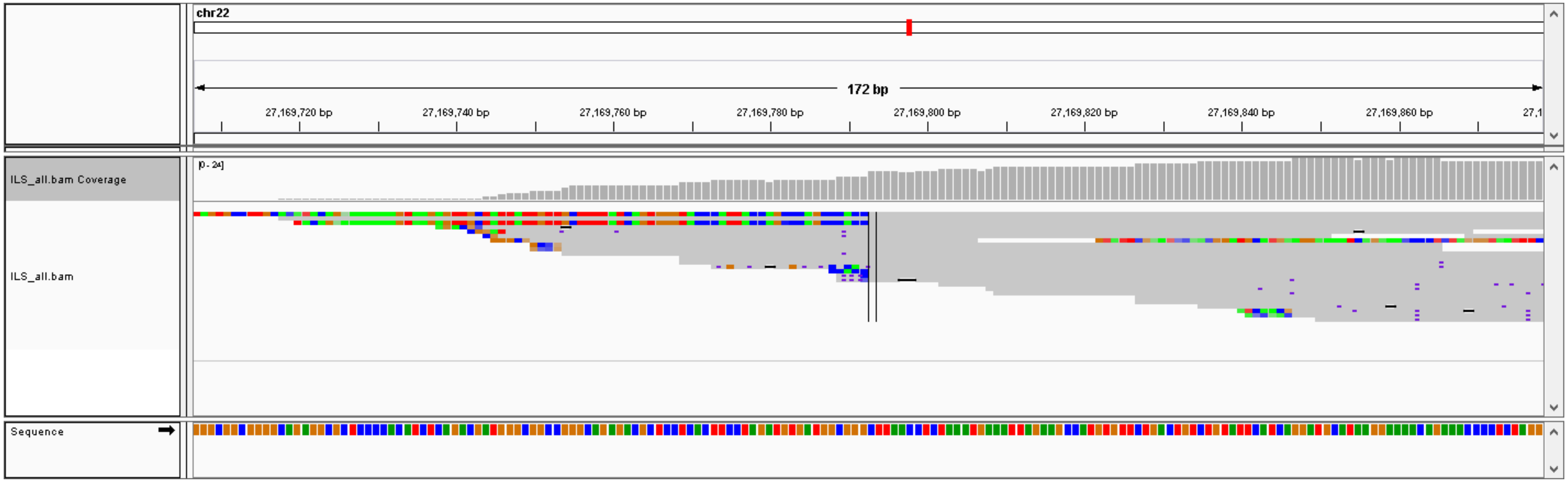

### #46, chr15: 56785305-56785365,  DEL (59 bp), validation: TP

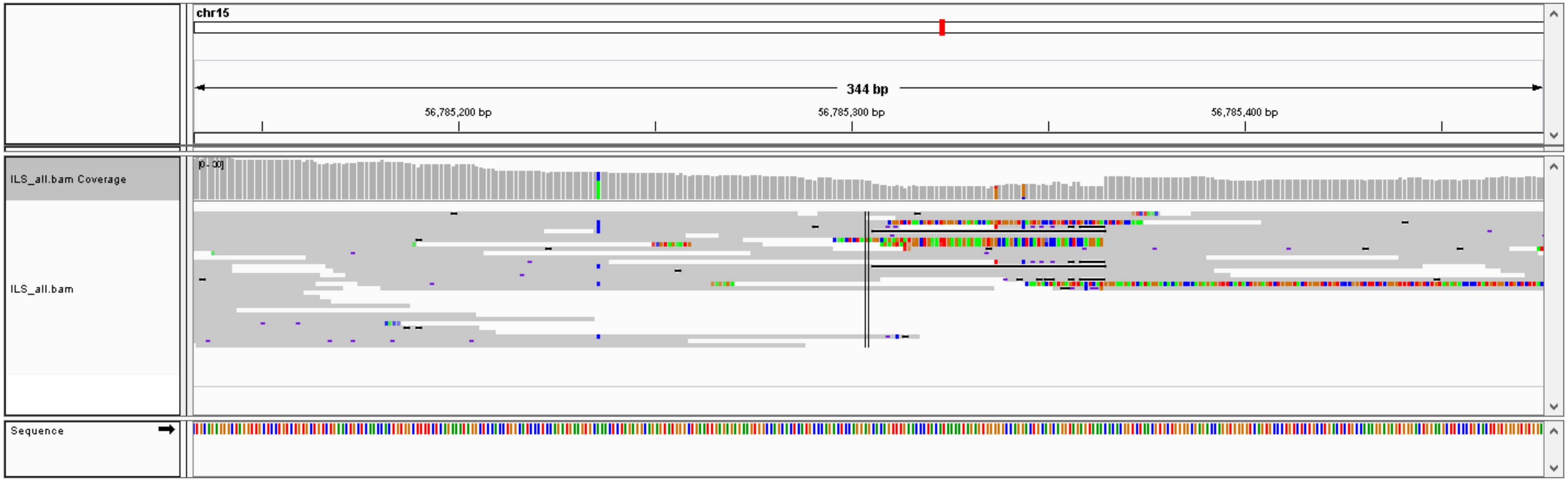

### #47, chr8: 72214745-72217809,  DEL (3063 bp), validation: TP

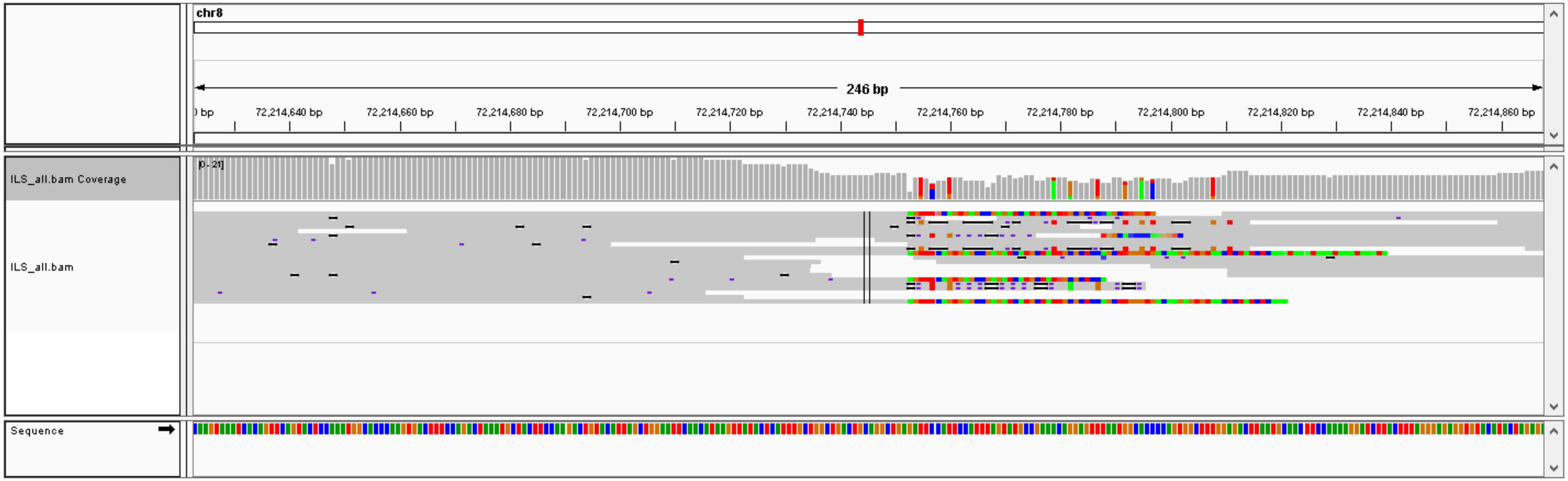


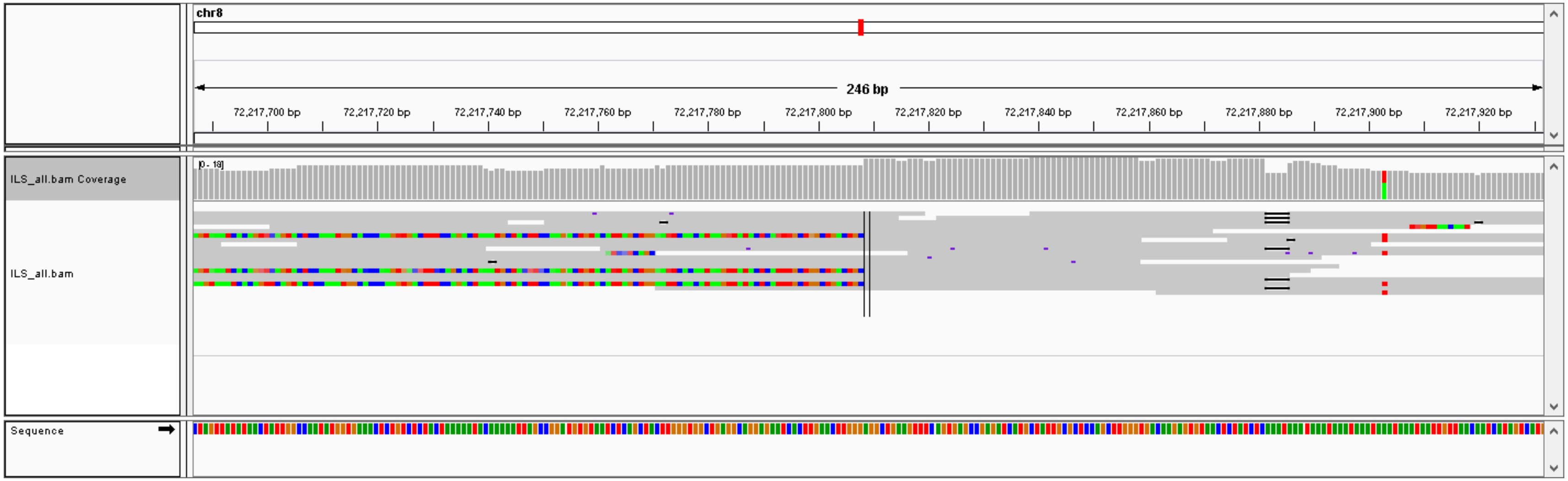

### #48, chr6: 5037249-5039187,  DEL (1937 bp), validation: TP

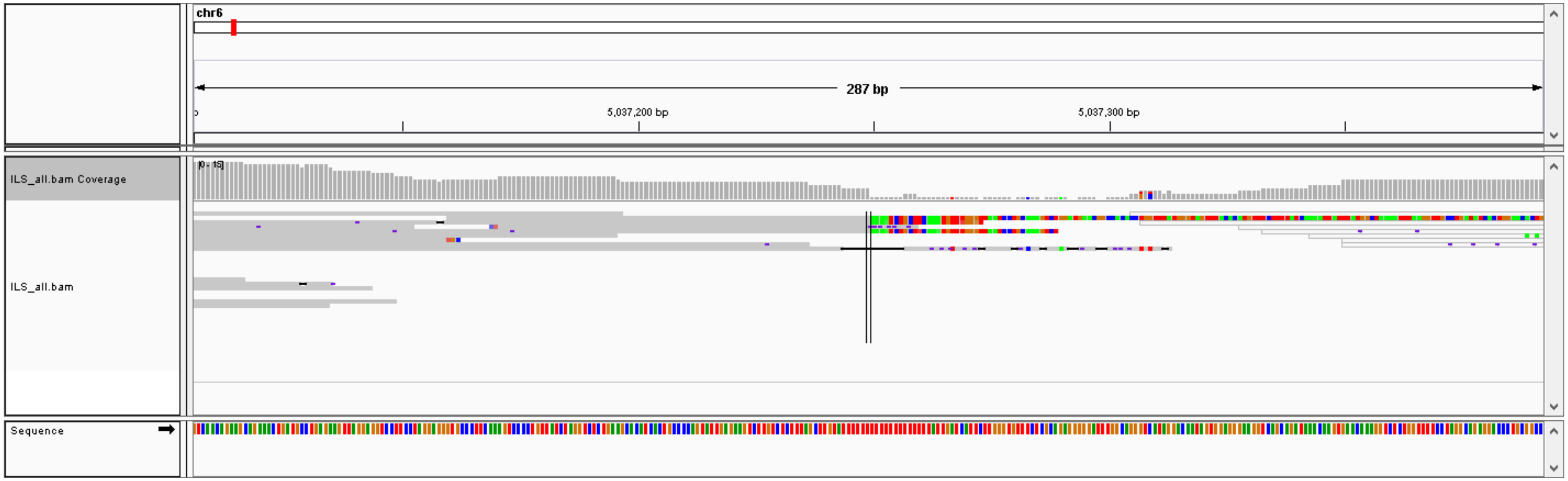


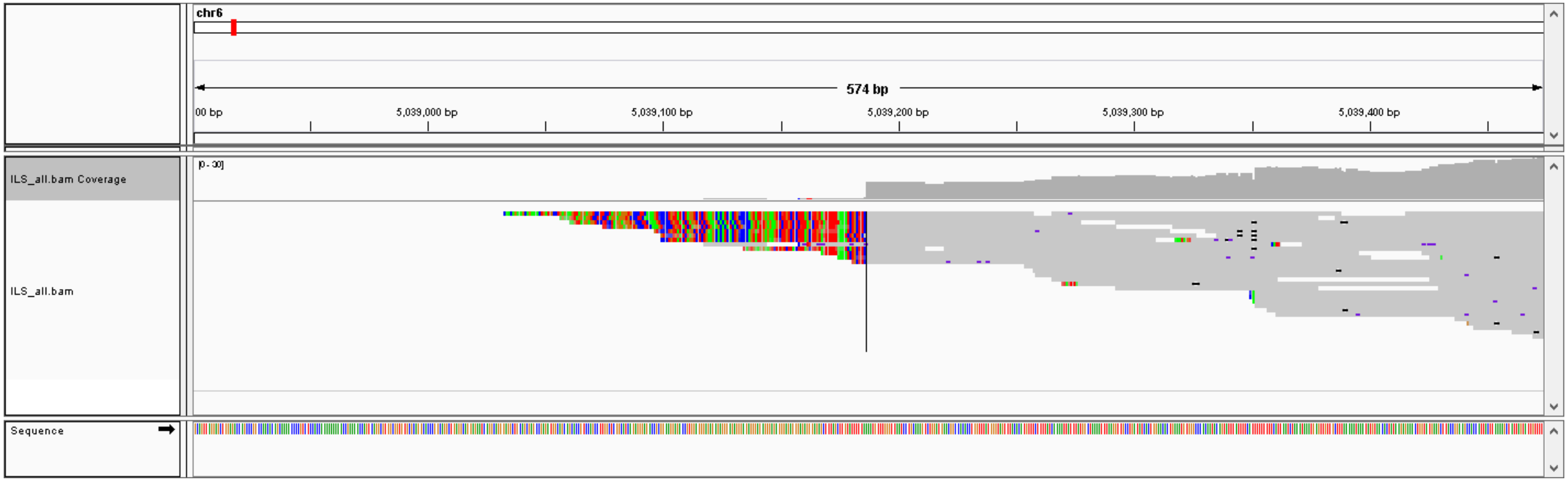

### #49, chr15: 68713338-68713559,  DEL (220 bp), validation: TP

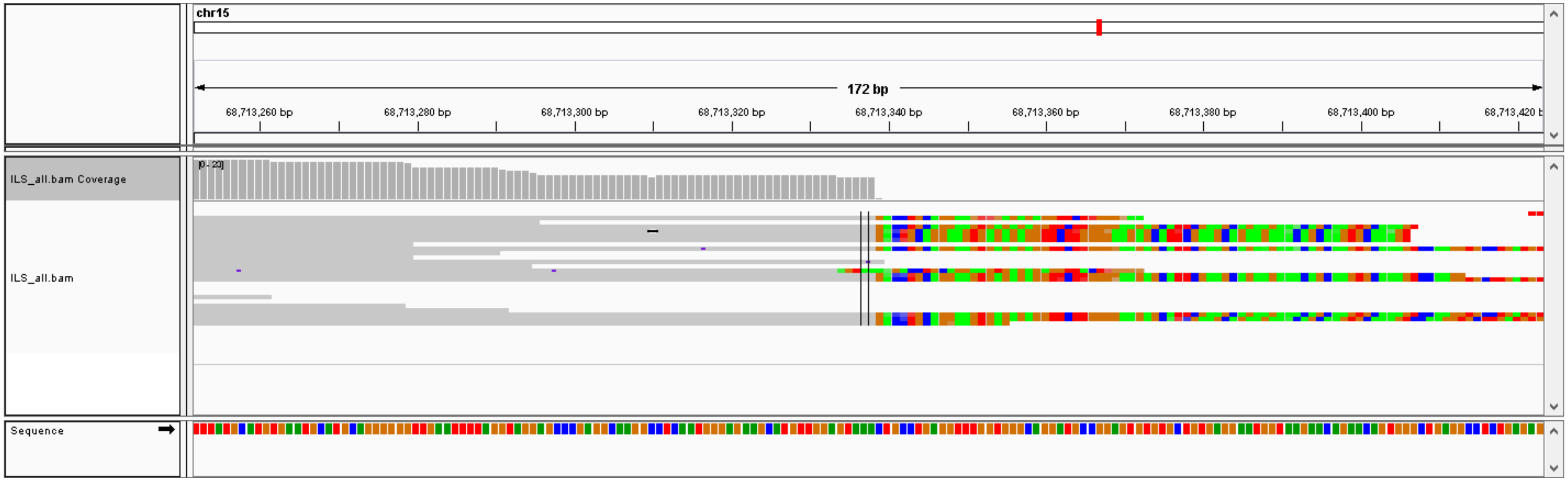


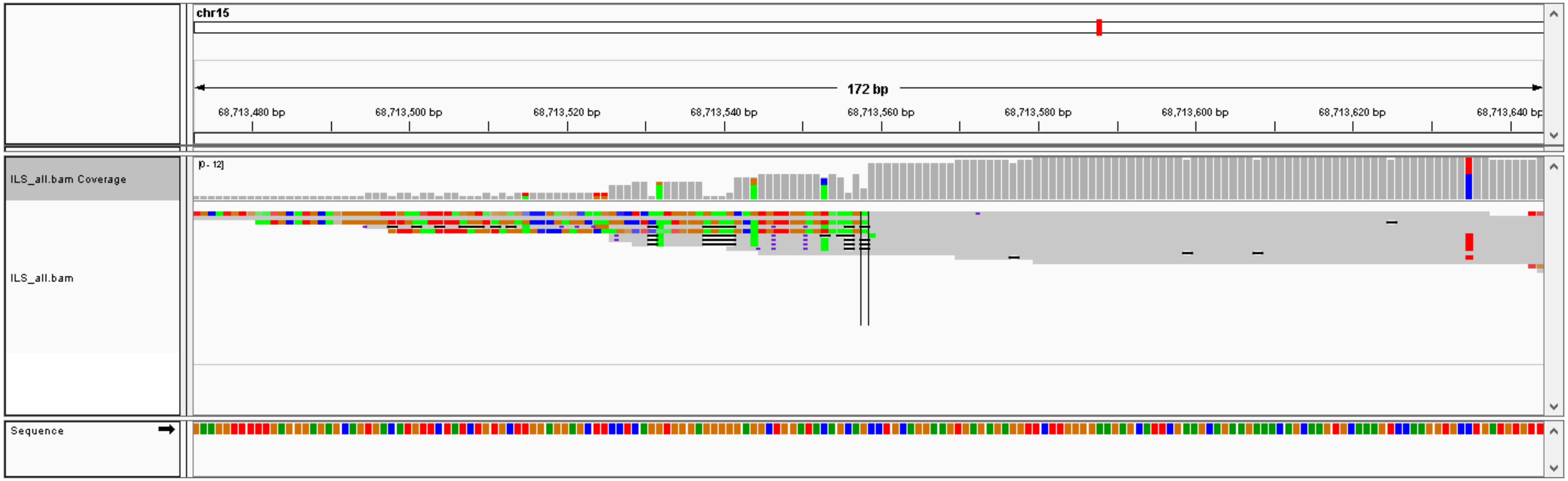

#50, chr10: 12334721-12334747, DEL (25 bp), validation: TP

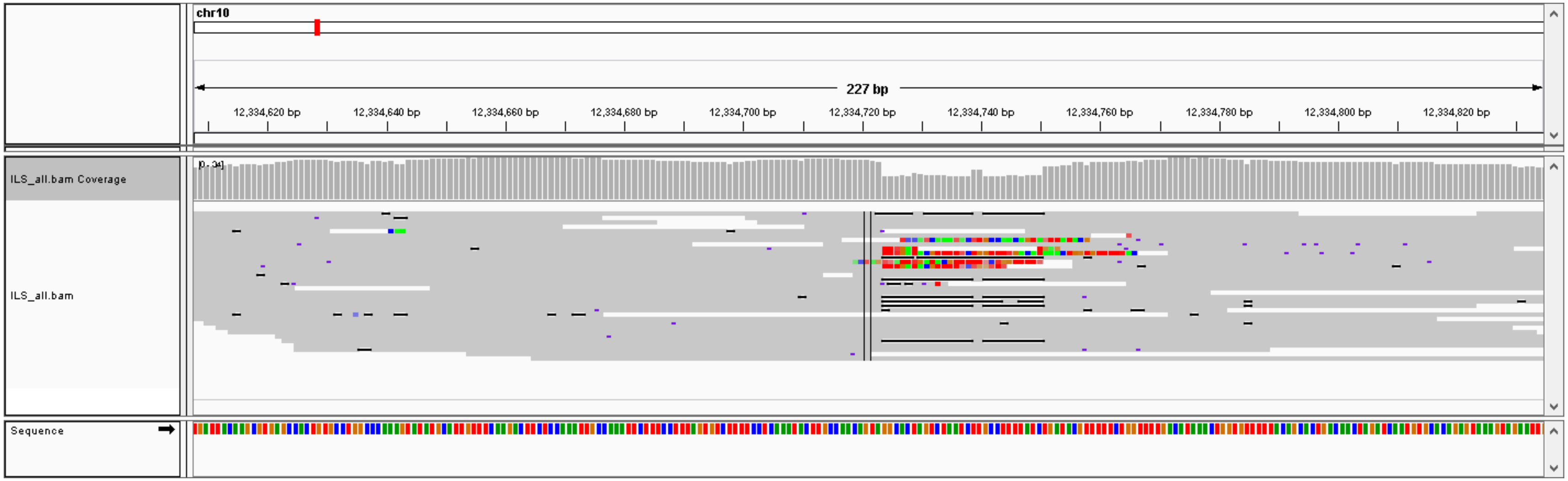

### #51, chr10: 12038776- chr1: 238441058,  CTX, validation: TP

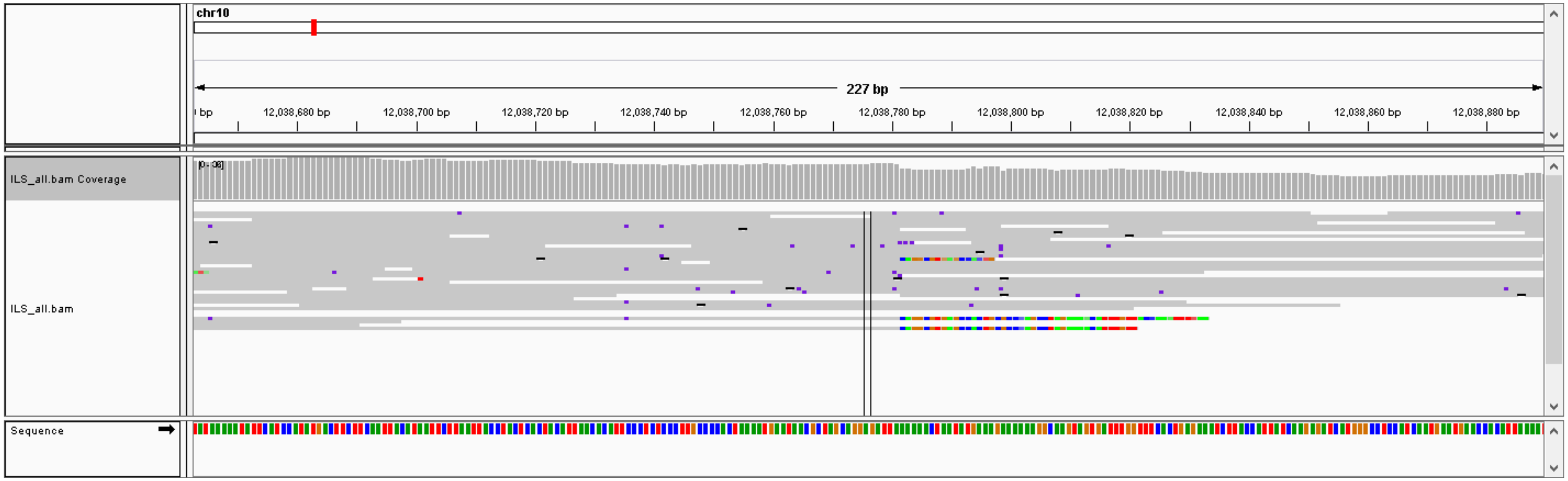


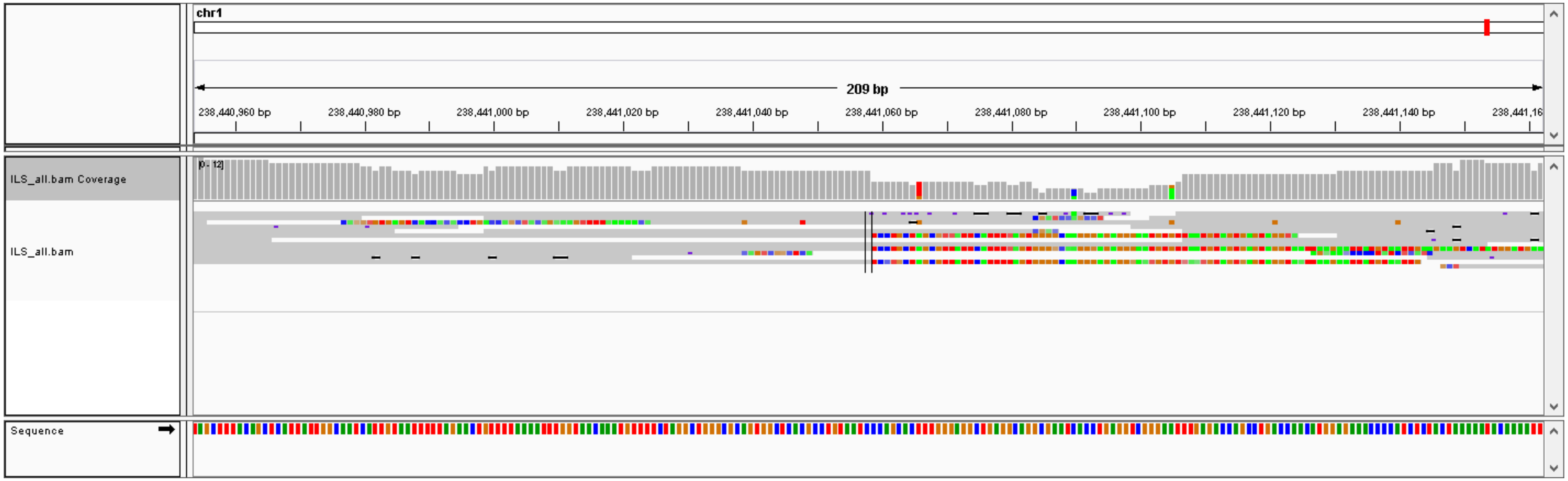

### #52, chr8: 15289367- chr13: 74313862,  CTX, validation: TP

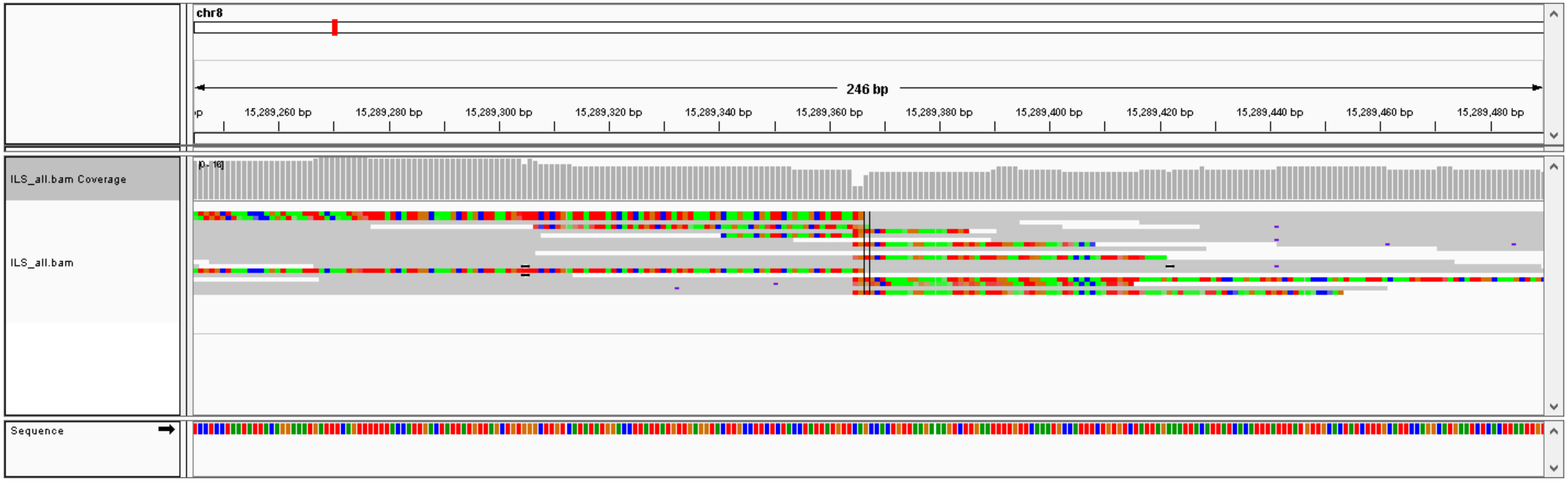


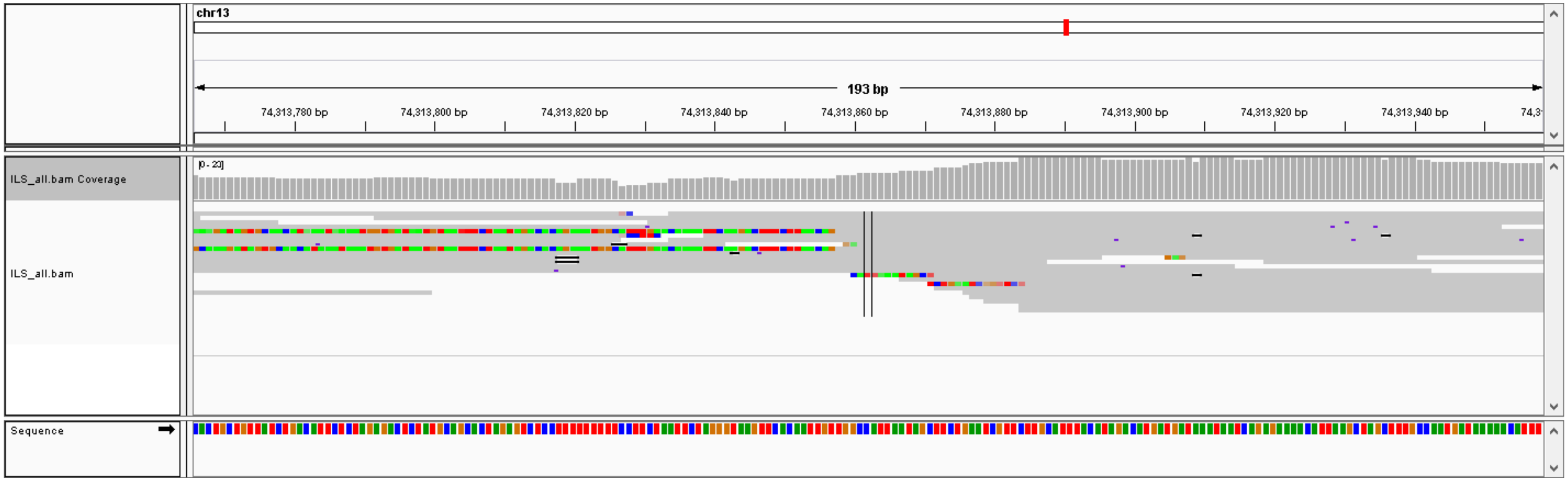

### #53, chr7: 152113758- chr21: 14742956,  CTX, validation: TP

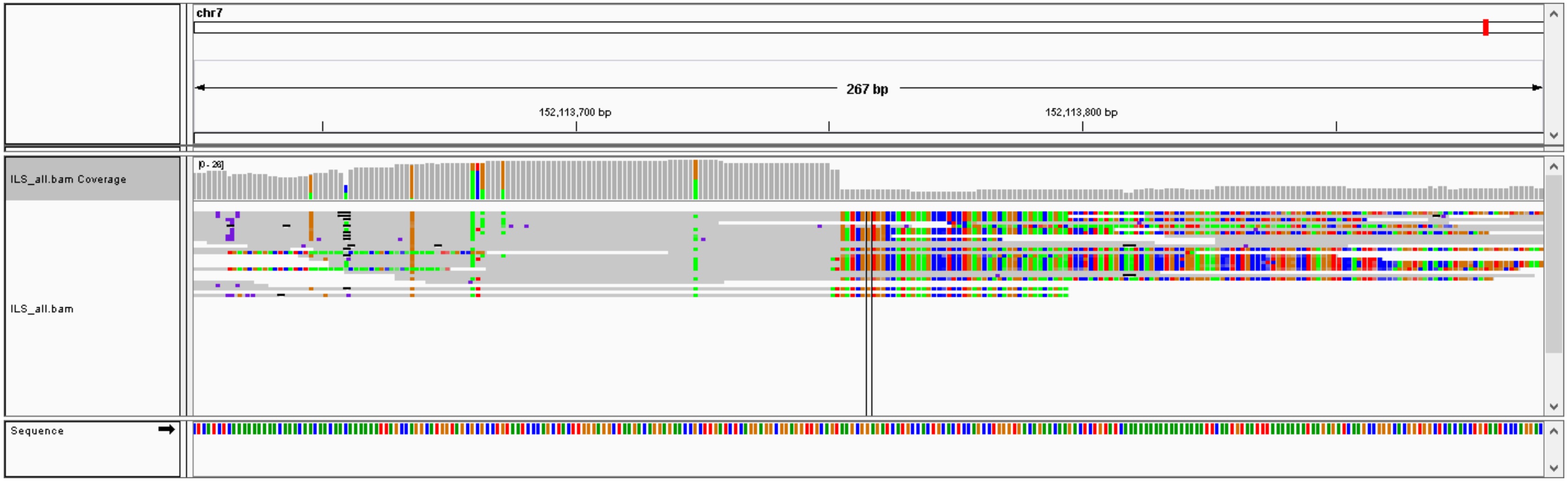


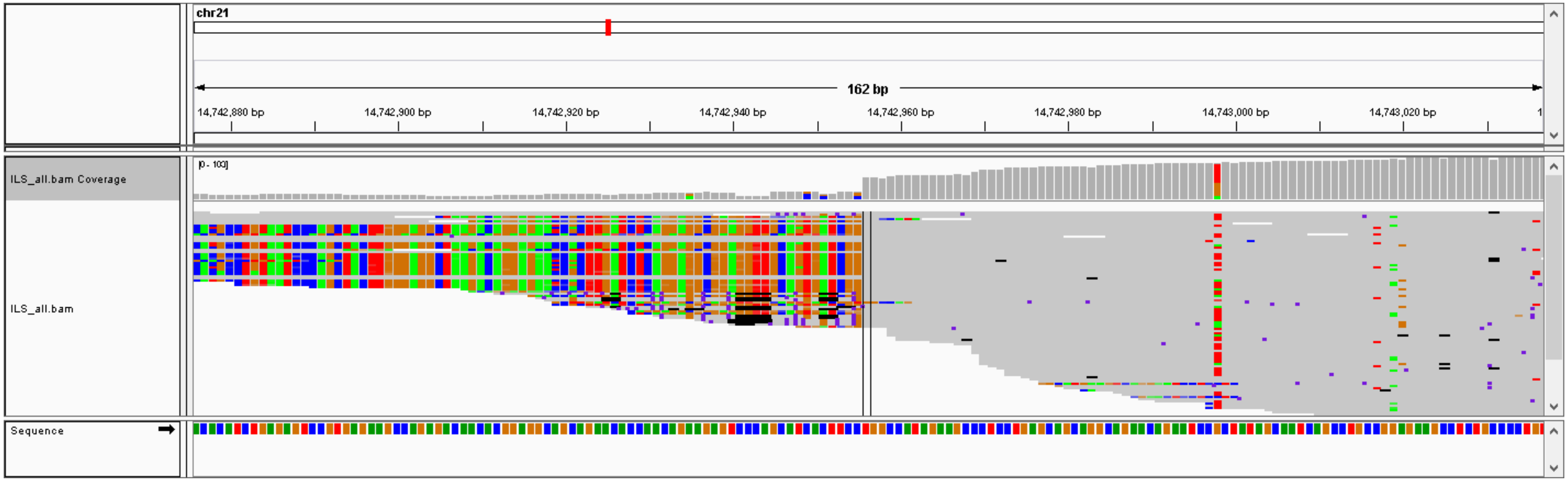

### #54, chr8: 52731478- chr11: 38812658,  CTX, validation: TP

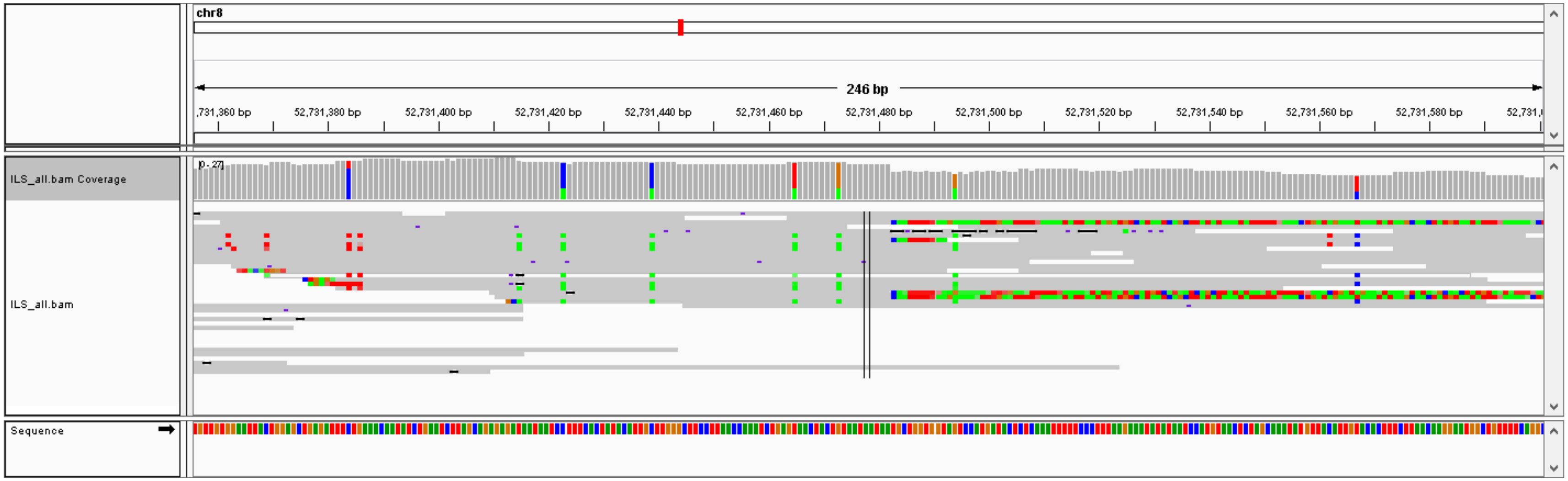


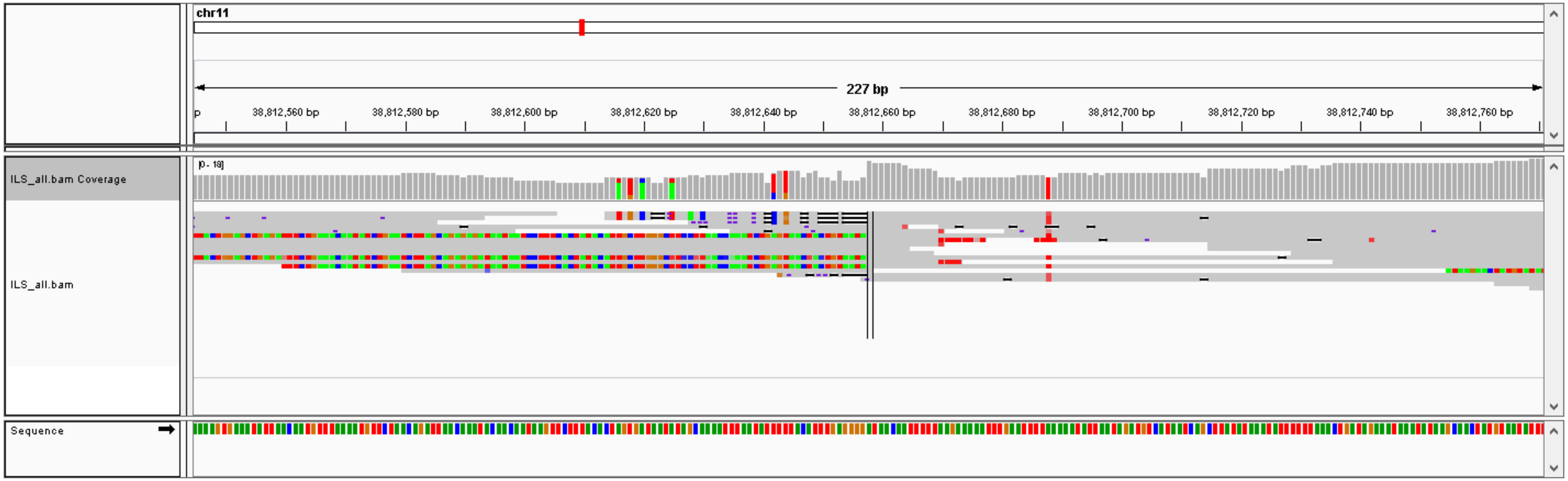

### #55, chr8: 37589180- chr14: 52200267,  CTX, validation: TP

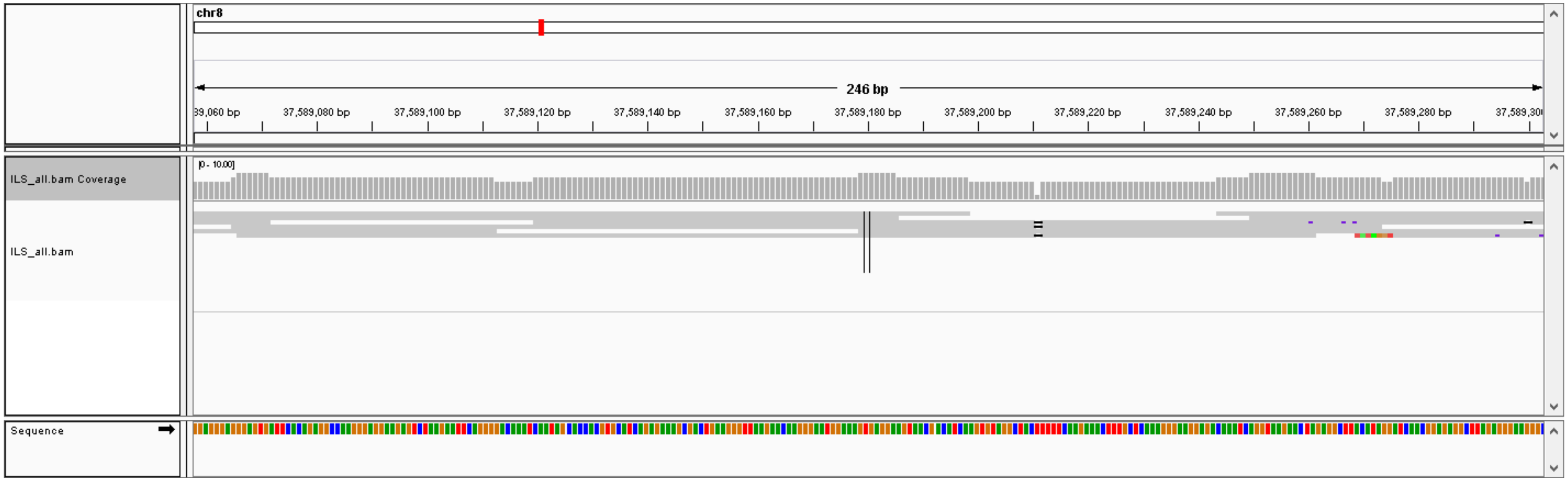


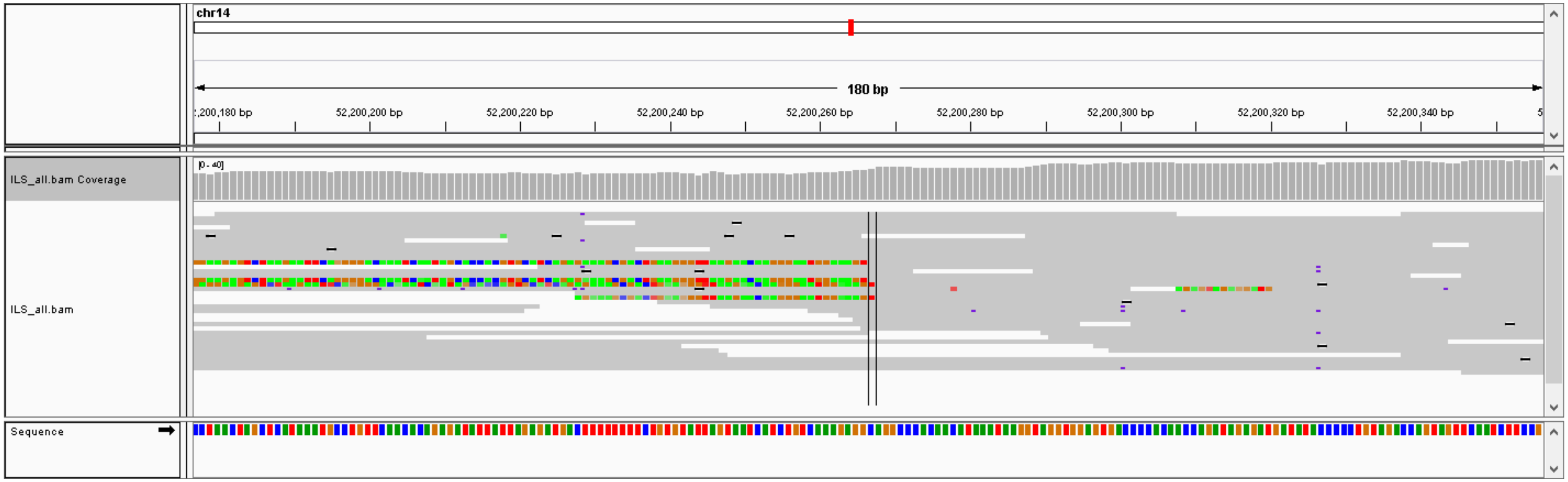

### #56, chr8: 30145402- chr17: 7167959,  CTX, validation: TP

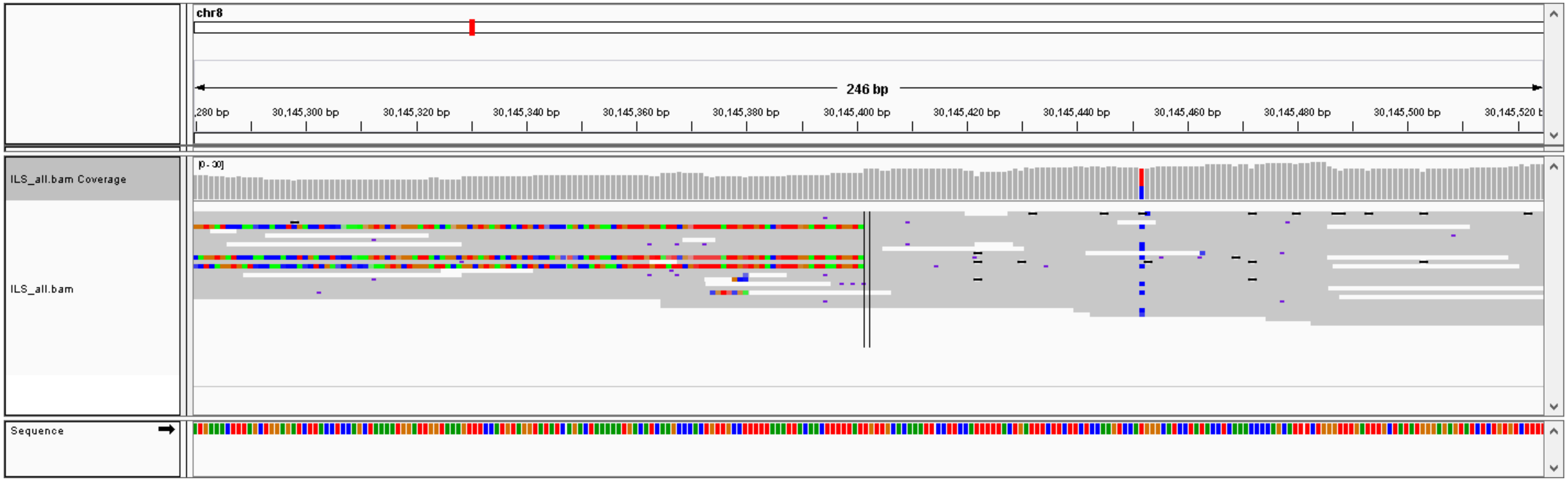


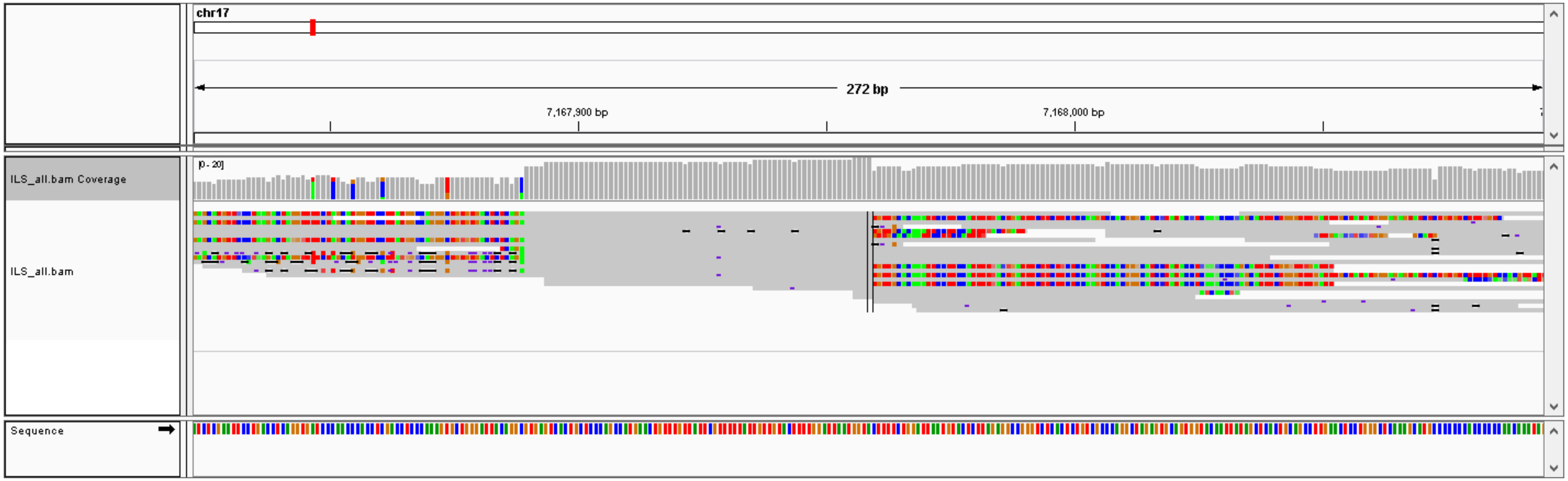

### #57, chr3: 110413395- chr1: 109495133,  CTX, validation: TP

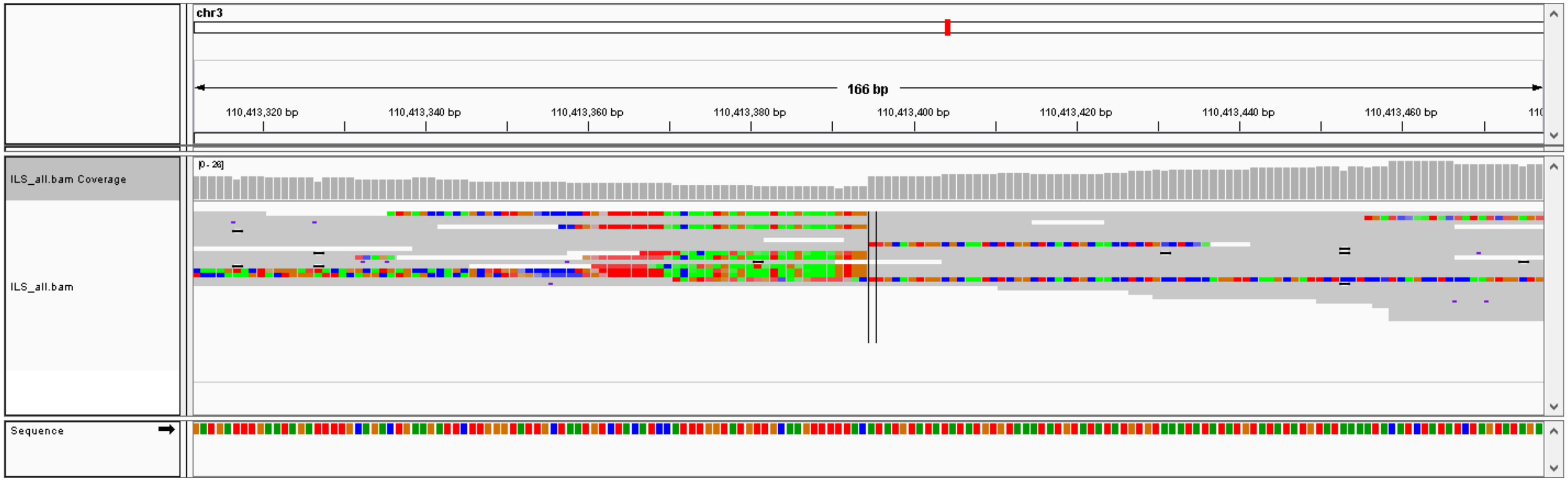


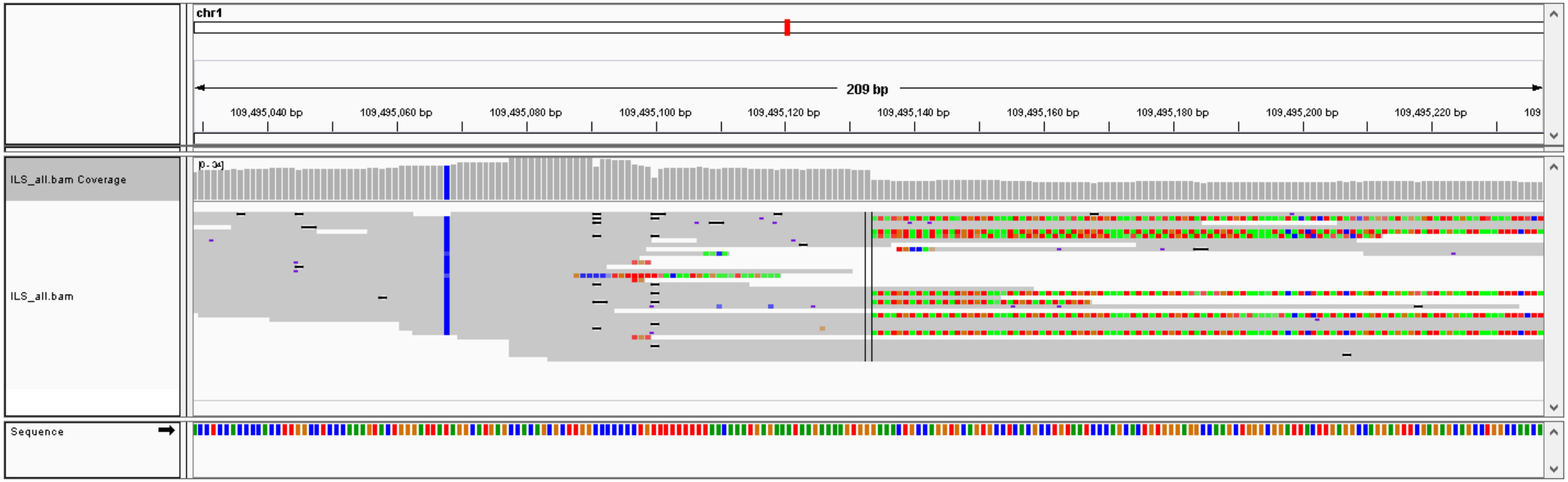

### #58, chr2: 190999445- chr1: 164240436,  CTX, validation: TP

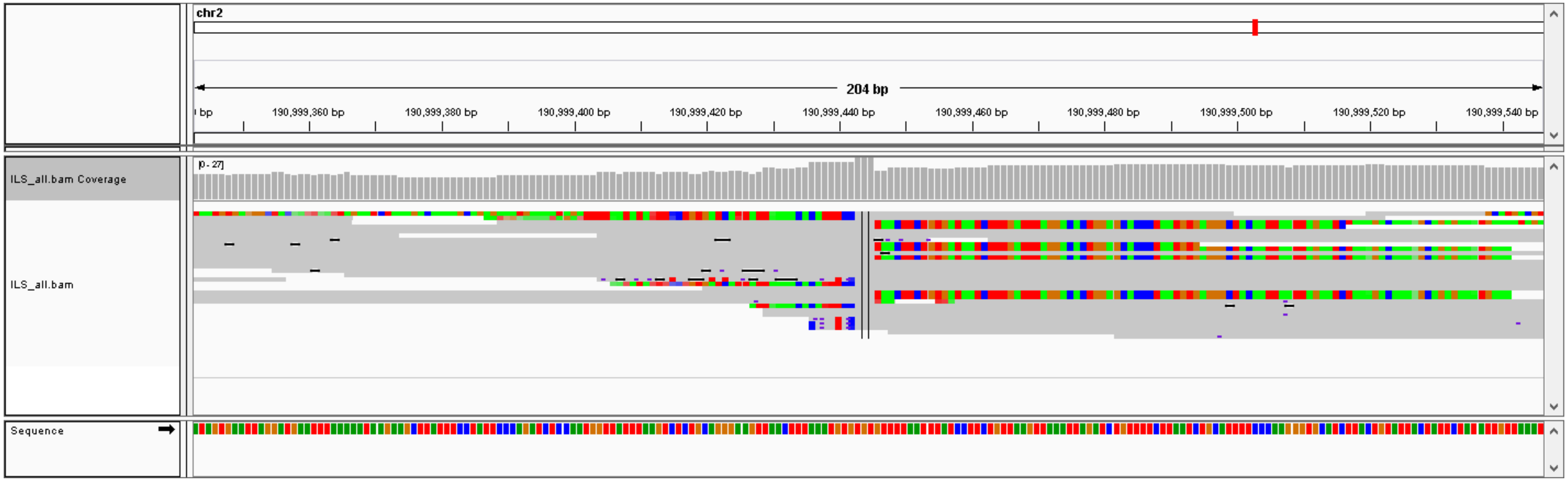


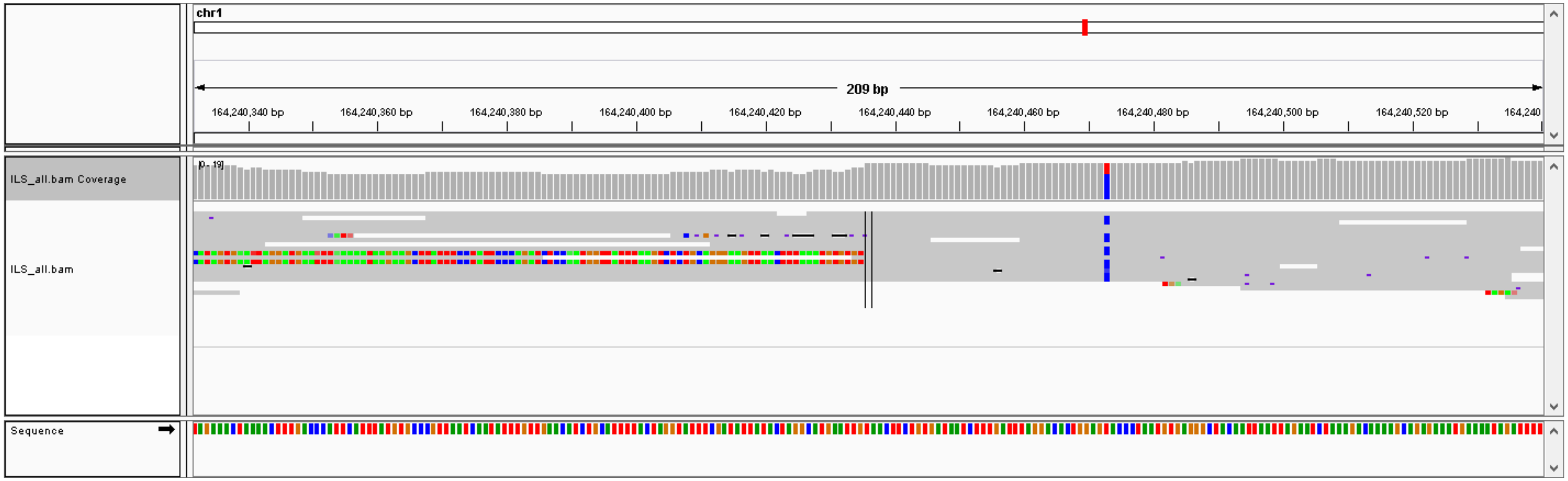

#59, chr4: 147953946- chr20: 54914484,  CTX, validation: TP

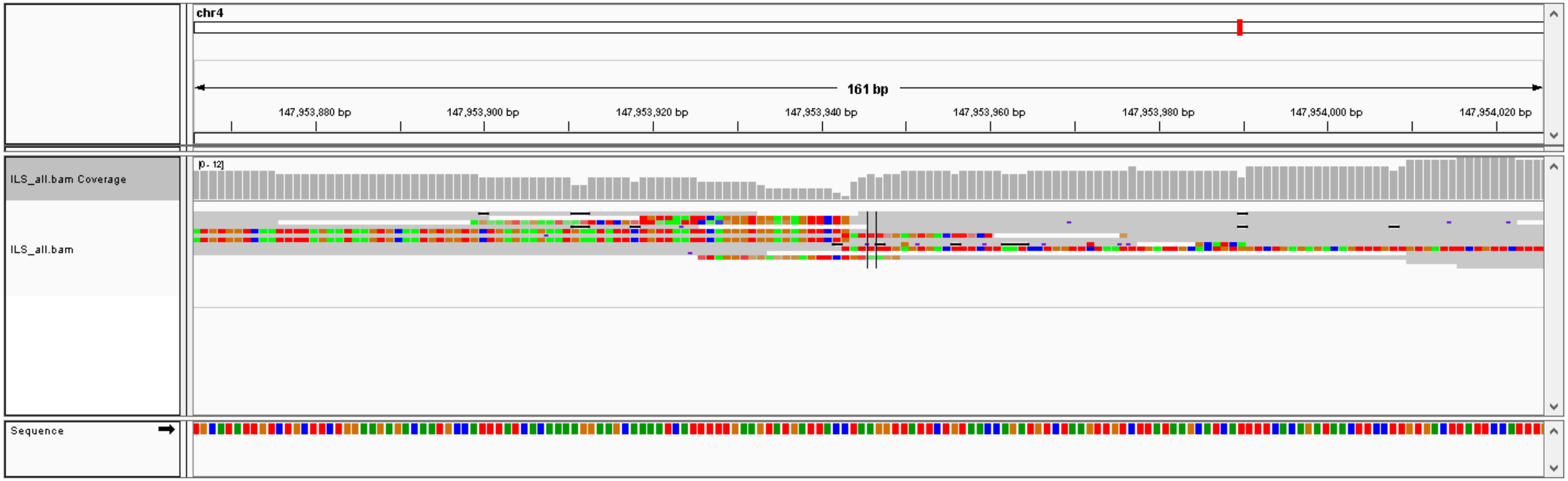


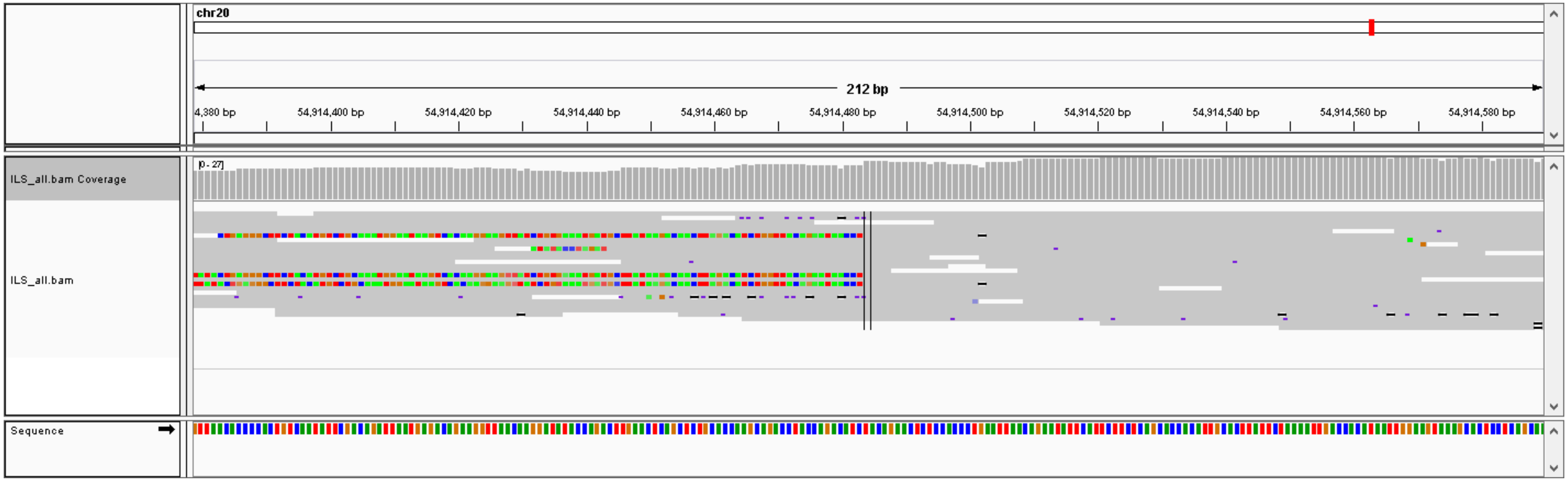

#60, chr7: 40835032- chr20: 49757402,  CTX, validation: TP

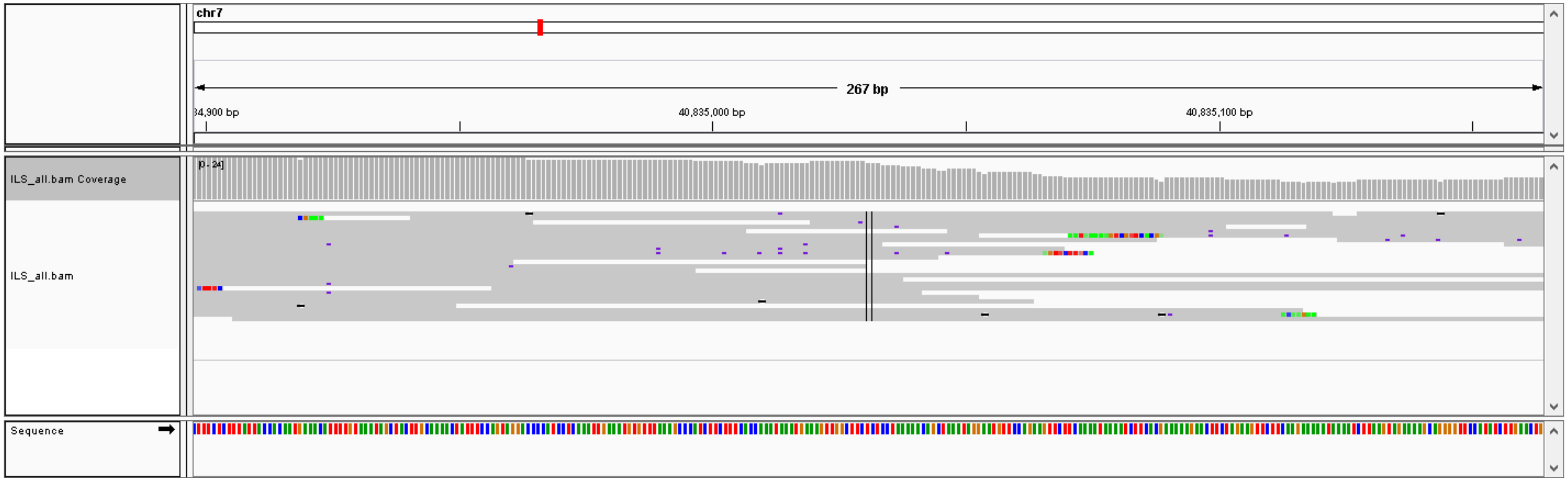


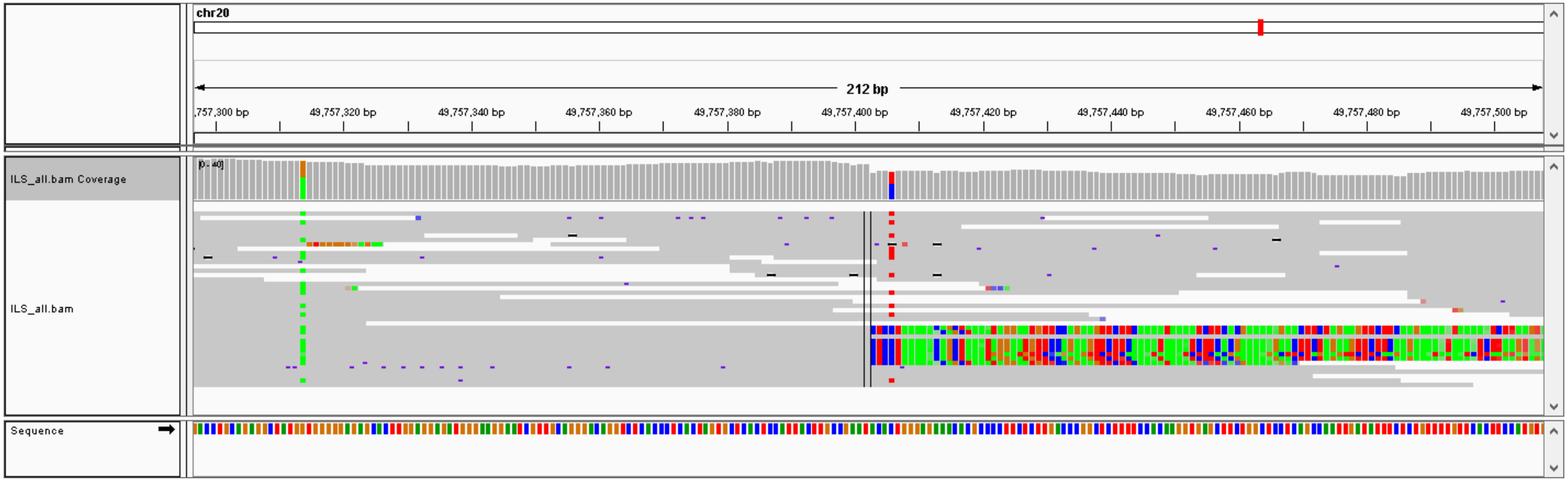
