## Supplementary material for "ReALLEN: structural variation discovery in cancer genome by sensitive analysis of single-end reads": Figure S5

A

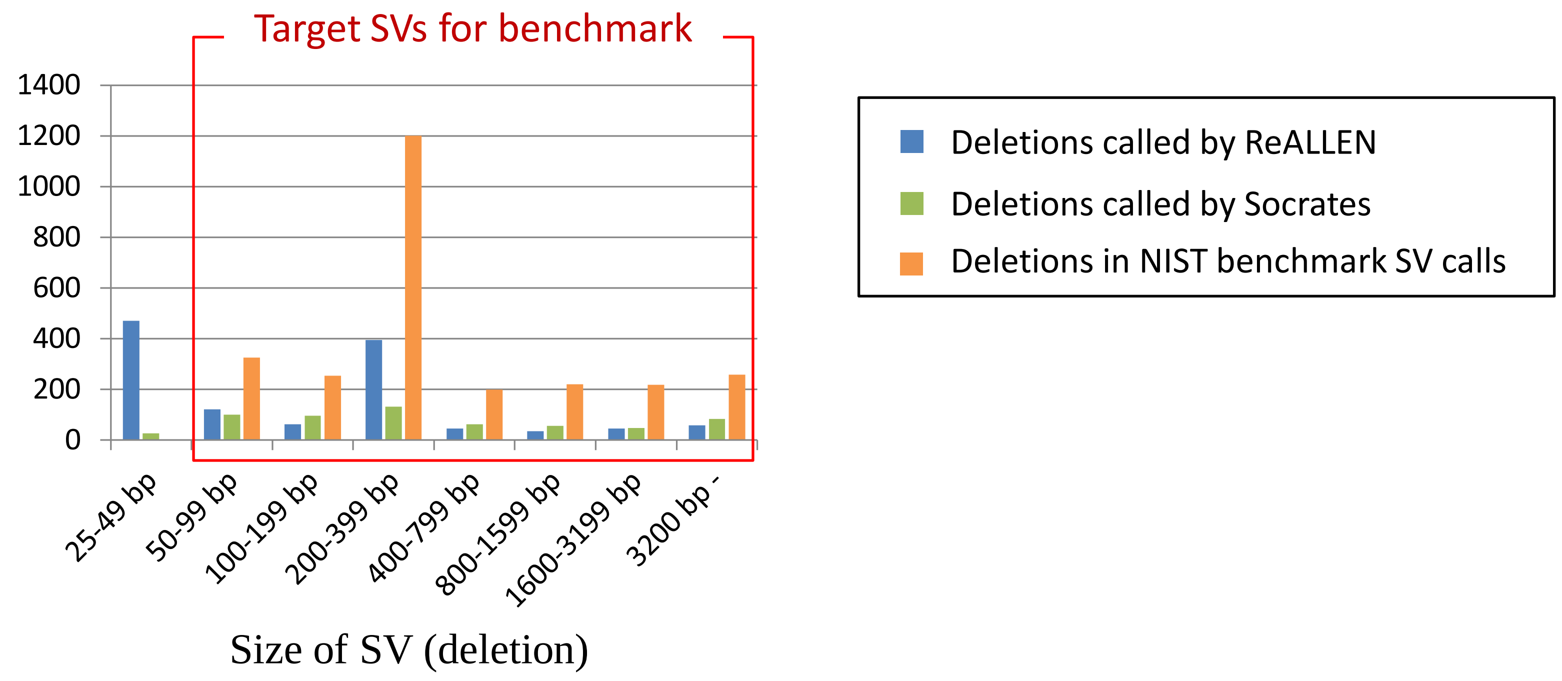


B

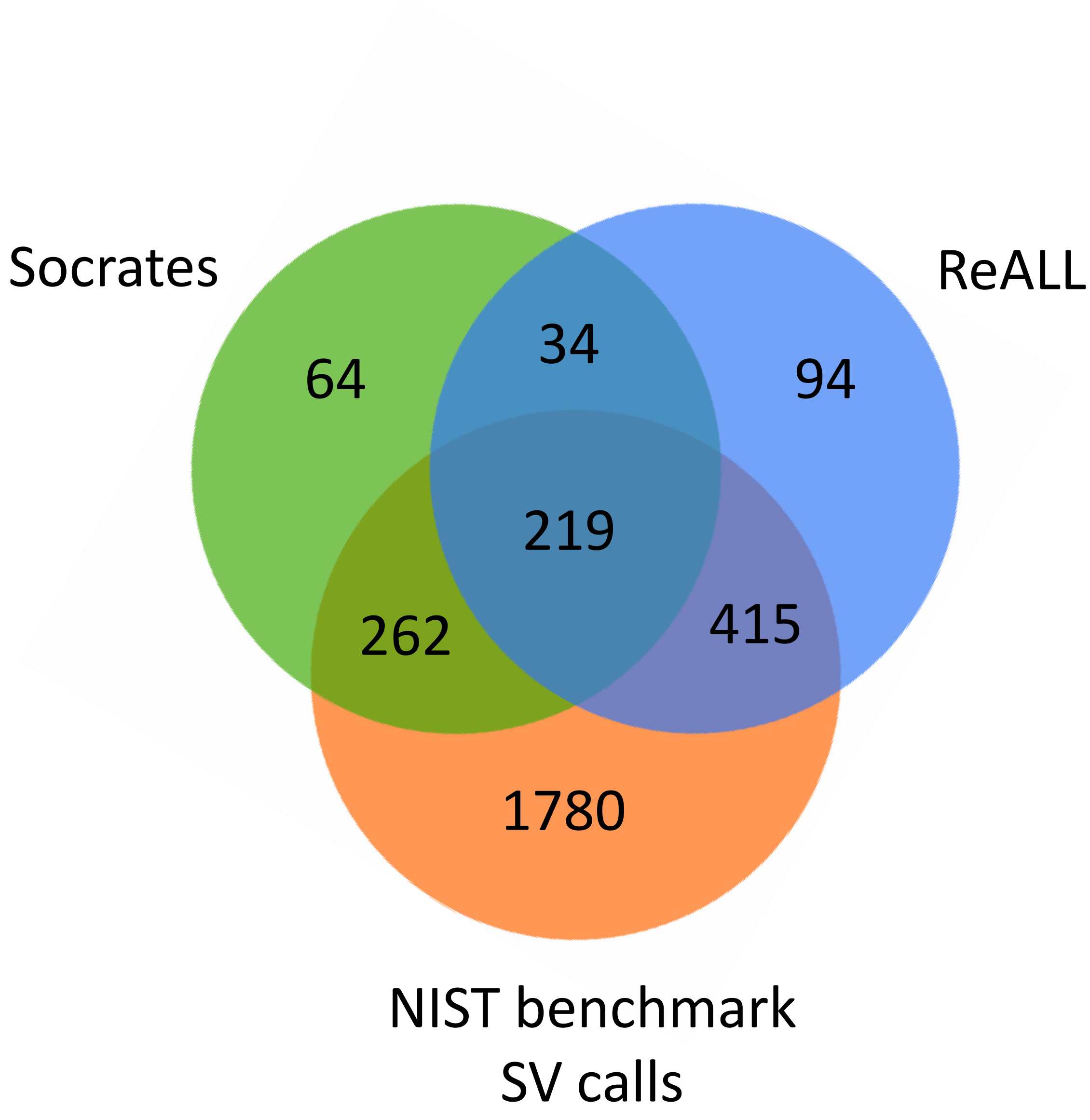


**Figure S5. Evaluation of ReALLEN and Socrates using the NIST benchmark SV calls.**
(A) Number of SVs for each size category included in the NIST benchmark SV calls, the calls by ReALLEN, and the calls by Socrates for NA12878. The deletions more than or equal to 50 bp (surrounded by red frame) were extracted for the comparison. (B) Venn diagram that summarizes the number of deletions in the NIST benchmark SV calls and the calls by ReALLEN and the calls by Socrates for NA12878. Two calls are considered to overlap if they have the same orientation and the locations within 20 bp.
